# CAR T cells recognizing CD276 and Dual-CAR T cells against CD276/FGFR4 promote rhabdomyosarcoma clearance in orthotopic mouse models

**DOI:** 10.1101/2023.09.05.555125

**Authors:** Andrea Timpanaro, Caroline Piccand, Dzhangar Dzhumashev, Stenija Anton-Joseph, Andrea Robbi, Janine Moser, Jochen Rössler, Michele Bernasconi

**Affiliations:** Department of Pediatric Hematology and Oncology, Inselspital, Bern University Hospital, 3010 Bern, Switzerland; Translational Cancer Research, Department for BioMedical Research (DBMR), University of Bern, 3008 Bern, Switzerland; Graduate School for Cellular and Biomedical Sciences, University of Bern, 3012 Bern, Switzerland

**Keywords:** Rhabdomyosarcoma, CAR T cells, CD276, FGFR4, immunotherapy, Dual-CAR T cells

## Abstract

**Background:** Rhabdomyosarcoma (RMS) is the most common soft tissue sarcoma in childhood, whose prognosis is still poor especially for metastatic, high-grade, and relapsed RMS. New treatments are urgently needed, especially systemic therapies. Chimeric Antigen Receptor T cells (CAR Ts) are very effective against hematological malignancies, but their efficacy against solid tumors needs to be improved. CD276 is a target upregulated in RMS and detected at low levels in normal tissues. FGFR4 is a very specific target for RMS. Here, we optimized CAR Ts for these two targets, alone or in combination, and tested their anti-tumor activity *in vitro* and *in vivo*.

**Methods:** Four different single-domain antibodies were used to select the most specific FGFR4-CAR construct. RMS cell killing and cytokine production by CD276- and FGFR4-CAR Ts expressing CD8α or CD28 HD/TM domains in combination with 4-1BB and/or CD28 co-stimulatory domains were tested *in vitro.* The most effective CD276- and FGFR4-CAR Ts were used to generate Dual-CAR Ts. Tumor killing was evaluated *in vivo* in three orthotopic RMS mouse models.

**Results:** CD276.V-CAR Ts (276.MG.CD28HD/TM.CD28CSD.3z) showed the strongest killing of RMS cells, and the highest release of IFN-γ and Granzyme B *in vitro*. FGFR4.V-CAR Ts (F8-FR4.CD28HD/TM.CD28CSD.3z) showed the most specific killing. CD276-CAR Ts successfully eradicated RD- and Rh4-derived RMS tumors *in vivo*, achieving complete remission in 3/5 and 5/5 mice, respectively. In CD276^low^ JR-tumors, however, they achieved complete remission in only 1/5 mice. FGFR4 CAR Ts instead delayed of Rh4 tumor growth. Dual-CAR Ts promoted Rh4-tumors clearance in 5/5 mice.

**Conclusions:** CD276- and CD276/FGFR4-directed CAR Ts showed effective RMS cell killing *in vitro* and eradication of CD276^high^ RMS tumors *in vivo*. CD276^low^ tumors escaped the therapy showing a correlation of antigen density and effectiveness. FGFR4-CAR Ts showed specific killing *in vitro* but could only delay RMS growth *in vivo*. Our results show that combined expression of CD276-CAR with other CAR does not reduce its benefit. Introducing immunotherapy with CD276-CAR Ts in RMS seems to be feasible and promising, although CAR constructs design and target combinations have to be further improved to eradicate tumors with low target expression.

## Background

Pediatric rhabdomyosarcoma (RMS) is the most common soft tissue sarcoma in children and young adults (1), and each year it accounts for 3% of childhood cancers (2). Based on histology, RMS can be classified in four main subtypes: embryonal RMS (eRMS; 60-70%), alveolar RMS (aRMS; 20-30%), pleomorphic RMS (pRMS) and spindle cell/sclerosing RMS (s-scRMS) accounting for 7–15% of the cases (3). RMS classification has been further refined in ‘fusion positive’ (FP) and ‘fusion negative’ (FN) RMS (4), based on two main characteristic chromosomal translocations, the t(2;13)(q35;q14) or the t(1;13)(p36;q14) resulting in the expression of a PAX3-FOXO1 or PAX7-FOXO1 fusion transcription factor, respectively (5–7). The onset of RMS has been attributed mainly to cells of myogenic lineage, but more recently also non-myogenic mesenchymal cells were indicated as possible RMS progenitors (8–11). As a result, RMS can develop not only in skeletal muscles, like head and neck, trunk, and extremities, but also in distant sites, including genitourinary and biliary tract. Despite overall 5-year survival rates have improved with the combined use of surgery, radiation therapy, and chemotherapy, in pediatric patients with metastatic and high-risk disease, such as aRMS, prognosis remains poor (12). Therefore, new therapies are needed for children and young adults with high-risk and recurrent RMS.

Immunotherapies using Chimeric Antigen Receptor (CAR) T cells (13, 14) have shown extraordinary results in patients with B cell malignancies (15), leading to the FDA approval of six CAR T cell therapies so far (16). Target molecules for these CAR T cells are surface proteins on leukemia or lymphoma cell surface, such as CD19 and CD20. However, CAR T cell therapy remains challenging for solid tumors, mainly due to a lack of ideal tumor specific target molecules, and to the immunosuppressive tumor microenvironment (TME) of solid tumors limiting the activity of CAR T cells (17, 18).

Several targets have been investigated pre-clinically and clinically for immunotherapies of pediatric solid tumors (reviewed in (19–21)), including for RMS. More than ten years ago, CAR T cell therapy for RMS has been explored targeting the fetal acetylcholine receptor gamma subunit (fAChRγ) surface protein, which is not present in humans after birth and is therefore highly restricted to RMS (22–24). The fAChRγ-CAR T cells exhibited specific, MHC-independent, *in vitro* lysis and cytokine secretion in response to antigen-expressing target cells and were moderately immunoprotective in murine tumor xenograft models (24). Unfortunately, their activity was not sufficient to eradicated tumors, probably due to fAChRγ low expression levels. Other surface proteins explored for CAR T cell therapy in RMS are IGF1R and ROR1. The corresponding CAR T cells showed effective *in vitro* RMS cell lysis, but have not been tested *in vivo* (25). Other cell surface antigens have been investigated as CAR T cells targets in RMS: PDGFRα-CAR T cells showed good activity *in vitro* and tumor control in a subcutaneous RMS xenograft mouse model (26), as PDGFRα is aberrantly expressed in RMS (27); HER2/ErbB2-CAR-engineered cytokine-induced killer cells showed potent activity on RMS spheroids (28), based on an expression level of the receptor tyrosine kinase HER2/ErbB2 in 10-33% of RMS on the protein level (29, 30). Importantly, in a phase I clinical trial of HER2/ErbB2-CAR T cells (NCT00902044), two complete responses were observed in RMS patients without any safety concerns (31). In particular, one patient with bone marrow metastatic involvement showed complete response after CAR T cell administration (31). Despite all these promising results, more targets and better CAR designs need to be investigated to cover all RMS patients and to avoid relapses mediated by antigen-escape.

In our previous work on identifying potential surface antigens by surfaceome profiling of RMS for an immunotherapy approach (32), we identified two targets that are particularly appealing for a CAR T cell therapy: CD276 and FGFR4.

The type I transmembrane protein CD276 (B7-H3) belongs to the B7 immune co-stimulatory and co-inhibitory family (33), and is expressed on antigen-presenting cells (APC) interacting with immune checkpoint markers, such as CTLA-4, PD-1, and CD28 (34). CD276 has both co-stimulatory and co-inhibitory functions in the regulation of different T cell subsets. CD276 is expressed at high levels in a large proportion of human malignancies, including the pediatric solid tumors RMS (35, 36), osteosarcoma (36, 37), intrinsic pontine glioma (38), medulloblastoma, and Ewing sarcoma (36). Importantly, an immune evasive role for CD276 in the context of RMS has been recently described, further supporting the important impact its targeting might have on disease control by the immune system (39). Moreover, CD276-CAR T cells showed efficacy in preclinical models of pancreatic ductal adenocarcinoma, ovarian cancer, and neuroblastoma without toxicity (40). Of note, a CD276-CAR expressing an antigen-binding domain derived from Enoblituzumab (MGA271), a humanized antibody from MacroGenics recognizing an epitope of CD276 with high tumor reactivity, was particularly effective against pediatric osteosarcoma, Ewing sarcoma and medulloblastoma (36), but has not been tested for RMS *in vitro*, nor *in vivo*.

The other target of interest, the Fibroblast Growth Factor Receptor 4 (FGFR4) is a receptor tyrosine kinase that shows very specific expression in RMS, and in normal tissues is only expressed at low levels in liver and in mature skeletal tissue (41–44). FGFR4-CAR T cells are active *in vitro* against both aRMS and eRMS cell lines (45, 46), and show effective response against disseminated disease *in vivo*, but not in a RMS orthotopic model (47). Interestingly, the combination of FGFR4-CAR T cells with pharmacological inhibition of the myeloid component in stroma allowed FGFR4-CAR T cells to eradicate orthotopic RMS tumors in mice (47).

The design of CAR constructs needs to be optimized to the special situation of RMS. Clearly, each domain of the CAR – the extracellular antigen-binding domain (ABD); the hinge domain (HD); the transmembrane domain (TM); and the intracellular co-stimulatory domains (CSD) - can impact on the activity of CAR T cells (48, 49). The ABD determines the affinity and specificity, while its efficacy is influenced by the target accessibility and density on the surface of tumor cells (50). HD/TM and CSD contribute to regulate the activation threshold (51). The TM domain can affect the stability of CAR immune synapses and regulates the signaling threshold. CD28- or CD8α-derived TM domains are the most frequently used. CAR T cells expressing CD28 TM domain show higher CAR stability on the T cell surface, and a stronger response, but may be prone to overactivation, resulting in activation-induced cell death (AICD) (52). On the other hand, CAR T cells with a CD8α TM domain may have a lower activation threshold, increasing their cytotoxic potential, but release lower levels of TNF-α and IFN-γ (51, 53). The clinically used CSDs are derived from 4-1BB or CD28 proteins. CSDs are necessary for proper activation, expansion, and most importantly, for persistence of CAR T cells. CARs containing a CD28 CSD show high cytokine release upon antigen recognition through activation of the PI3K pathways, promoting the survival of cytotoxic CAR T cells (54, 55); whereas 4-1BB confers higher T cell persistence by promoting central memory T cell differentiation through the activation of the NF-κB signaling pathways (55, 56).

Moreover, previous reports have demonstrated that CD28 co-stimulation enhances exhaustion due to continuous CAR signaling, while 4-1BB co-stimulation mitigates it (57). T cell exhaustion is primarily influenced by the expression of inhibitory receptors. Following acute T cell activation, inhibitory receptors, such as PD-1 and CTLA-4, are upregulated to counteract TCR and co-stimulatory receptor signals, and to prevent excessive immune cell response. However, in the context of cancer, chronic tumor antigen exposure may lead to dysfunctional and exhausted T cells, due to reduced proliferation, cytokine production, increased apoptosis, and expression of inhibitory receptors, such as PD-1, TIM-3, and LAG-3 (58).

PD-1, an early exhaustion marker, is extensively under investigation in this context. It is rapidly upregulated upon T cell activation, but its expression alone is not exclusive to exhausted T cells (59). Alongside PD-1, exhausted T cells exhibit various inhibitory molecules on their cell surface. LAG-3, an intermediate exhaustion marker, shares similarities with CD4 (60) and is found in T cells, NK cells, Tregs, activated B cells, plasma cells, and dendritic cells (61, 62). By contrast, TIM-3 and CD39 are considered terminal dysfunction markers (58). TIM-3, T cell immunoglobulin and mucin domain 3, is predominantly expressed at higher levels by CD8^+^ tumor-infiltrating lymphocytes (TILs) and CD4^+^ Tregs (63). CD39, an ectonucleoside triphosphate diphosphohydrolase expressed on the T cell surface, dephosphorylates ATP released from damaged cells, generating extracellular adenosine and driving the suppression of immune-stimulatory effects downstream (64).

Here, we aimed at selecting *in vitro* the most active CAR construct targeting CD276, FGFR4, or the combination of both, on RMS cells. A combinatorial panel of eight CAR constructs, comprising second- and third-generation constructs, with CD8α or CD28 HD/TM domains and intracellular CSDs from 4-1BB and/or CD28 were produced and tested. The most promising CAR T cells were tested for their activity *in vivo* in orthotopic RMS mouse models.

## Materials and Methods

### Cell culture

Human RMS cell lines RD, Rh4, Rh5, Rh18, Rh28, Rh30, Rh36, JR, RMS, RUCH-3, TTC-442 were kindly provided by Prof. Beat Schäfer (University Children’s Hospital of Zurich). PDXs IC-pPDX-104 from a 7-year-old female with recurrent primary aRMS with PAX3-FOXO1 translocation, IC-pPDX-35 from a 13-year-old male with recurrent metastatic aRMS with PAX3-FOXO1 translocation, were established at the Institut Curie in Paris, France (65, 66); PDX RMS-ZH003 was established from a recurrent aRMS with PAX3-FOXO1 translocation at the University Children’s Hospital Zurich, as described in (65, 66). Immortalized human healthy primary myoblasts KM155C25Dist, kindly provided by the platform for immortalization of human cells MyoLine from the Institut de Myologie (Sorbonne University, Paris, France), referred to as myoblasts, were used as negative controls. All the cell lines, with the exception of human myoblasts, were cultured in DMEM medium (BioConcept, #1-26F01-I), supplemented with 10% fetal bovine serum (FBS, Thermofisher, #10270106), L-glutamine 2 mM (BioConcept, #5-10K00-H) and 100 U/ml Penicillin-Streptomycin (BioConcept, #4-01F00-H) at 37°C and 5% CO_2_ in a humidified incubator. Myoblasts were cultured in Skeletal Muscle Cell Growth Medium (PromoCell, #C-23060) supplemented with Skeletal Muscle Cell Growth Medium SupplementMix (PromoCell, #C-39365). Jurkat cells were cultured in RPMI-1640 medium (Gibco, #42401-018), supplemented with 10% fetal bovine serum (FBS, Thermofisher, #10270106), L-glutamine 2 mM (BioConcept, #5-10K00-H) and 100 U/ml Penicillin-Streptomycin (BioConcept, #4-01F00-H) at 37°C and 5% CO_2_ in a humidified incubator. PDXs were cultured in pre-coated P10 dishes with 1:10 diluted Matrigel (Corning, #354234) in 3 ml of pre-cooled Advanced DMEM/F-12 (Gibco, #12634010), supplemented with Glutamax (Gibco, #35050), 100 U/ml Penicillin-Streptomycin (BioConcept, #4-01F00-H), 1x B-27 (Life Technologies, #17504044), 20 ng/ml bFGF (PeproTech, #AF-100-18B), 20 ng/ml EGF (PeproTech, #AF-100-15), and 1.25 mM N-acetyl-L-cysteine (Sigma, #A9165).

### Plasmids

To express truncated-CD19 protein (tCD19) on RMS cell lines as target for CD19-CAR T cells, the lentiviral plasmid pLenti-fLuc-P2A-tCD19 was generated by excising with AjiI and MluI digestion the Neomycin resistance in the plasmid Lenti-luciferase-P2A-Neo (Addgene, #105621) and replacing it by GIBSON assembly with the tCD19 sequence amplified by PCR from the vector pcDNA3.1+CD19-C-DYK (Genescript, #OHU27320D) using the primers P2A-tCD19 Forward and P2A-tCD19 Reverse (Suppl. Table S1).

### Production of lentiviruses

Lentiviruses were generated by transient co-transfection of Lenti-X 293T cells (Takara, #632180). Briefly, 5×10^6^ Lenti-X 293T cells were seeded onto 10 cm dish, pre-coated with collagen type I rat tail (1:10 diluted in sterile water, Sigma, #9007-34-5), and the day after, transfected with the calcium phosphate method. 10 µg of psPAX2 (Addgene, Plasmid #12260), 4 µg of pVSV-G (Addgene, Plasmid #8454), and 10 µg of lentiviral plasmid of interest were used. The plasmids were mixed in 500 µl 0.25 M CaCl_2_, added dropwise to 500 µl of HBS 2x (NaCl 0.28 M, HEPES 0.05 M, Na_2_HPO_4_ 1.5 mM, pH 7.05), and incubated for 15 min at room temperature. The solution was then dropwise added to Lenti-X 293T cells. Lentiviruses-containing supernatants were harvested 48h post-transfection, filtered with 0.45 µm filters, and concentrated via overnight centrifugation at 4’000xg at 4°C. Lentiviruses were aliquoted and stored at −80°C until use.

### Lentiviruses titration

Titration of Lentiviruses was carried out with Jurkat T cells. 250’000 cells were plated onto 24-well plates and were transduced with ten-fold dilutions of lentiviruses. 48h post-transduction, fluorescent cells were quantified by Flow Cytometry. The titer was calculated with the following formula:

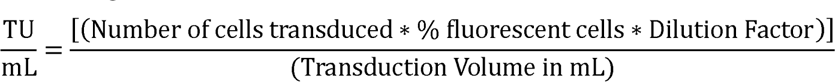

### Firefly Luciferase (fLuc^+^) RMS cell lines

pHIV-iRFP720-E2A-Luc was a gift from Antonius Plagge (Addgene plasmid # 104587) and was used to generate iRFP720^+^fLuc^+^ lentiviruses and to stably transduce RD and Rh4 cell lines at an MOI of 5. iRFP720^+^fLuc^+^ RMS cell lines were sorted 48h after transduction and amplified.

### Truncated CD19 (tCD19^+^) RMS cell lines

The pLenti-fLuc-P2A-tCD19 plasmid was used to generate tCD19^+^fLuc^+^ lentiviruses to stably transduce RD and Rh4 cells at an MOI of 5. 48h after transduction, tCD19^+^fLuc^+^ RMS cell lines were stained with BV421-conjugated anti-CD19 antibody for 30 min at room temperature, sorted by BV421, and amplified.

### CRISPR/Cas9

Two in silico-designed sgRNA sequences targeting CD276 and FGFR4 genes, respectively, were selected by using the Graphical User Interface for DNA Editing Screens (GUIDES), available at guides.sanjanalab.org (67). The two sgRNA sequences (Suppl, Table S2) were cloned into the pLentiguide-PURO plasmid (Addgene, #52963). The plasmid pLentiCRISPR-mCherry (Addgene, #75161) was used to stably express Cas9 nuclease and mCherry in RD and Rh4 cell lines. 48h after lentiviral transduction at an MOI of 5, cells expressing mCherry were sorted by FACS. Cas9^+^mCherry^+^ RMS cells were then transiently transfected with pLentiguide-PURO plasmid and selected with 1 µg/ml Puromycin (InVivoGen, #ant-pr-1) for two days, single cell cultures were then generated by limiting dilutions, and cells were expanded.

### CAR plasmids generation

The lentiviral plasmids (Suppl. Table S3) used for the CAR T cell production were generated with GeneArt Gibson assembly HiFi cloning kit (Thermofisher, #A46624EA). The sequences of FMC63 scFv targeting CD19, CD8αHD, and CD28TM CAR domains were amplified from the plasmids pSLCAR-CD19-hCAR-BB (Addgene, #135992) and pSLCAR-CD19-28z (Addgene, #135991) kindly provided by Dr. Scott McComb (National Research Council of Canada, Ottawa, Canada) (68). The sequence of CD276.MG scFv was kindly provided by Prof. Robbie Majzner (Stanford University, Stanford, CA). To generate the CD276 and FGFR4 CAR constructs, the FMC63 scFv, flanked by Esp3I sites, was swapped with the PCR-amplified sequences of CD276.MG scFv or FGFR4 sdAb (A1-, B1-, B5- and F8-FR4 sdAbs). The swapping reaction was performed by incubating the DNA samples with Esp3I (BsmBI) (Thermofisher, #ER0451) and T4 ligase (Thermofisher, #EL0011) at 37°C for 10 min, followed by 5 cycles at 16°C 10 min and 37°C 60 min and a final enzyme inactivation at 80°C for 5 min (69). The plasmids were then transformed in One Shot TOP10 chemically competent cells (Thermofisher, #C404010), amplified, and purified by NucleoBond Xtra Midi kit (Macherey-Nagel, #740410.50). Purity and concentration were evaluated with Nanodrop. Plasmids were sequenced at Microsynth (Balgach, Switzerland). All plasmids containing the CD19 targeting ABD FMC63 are available from Addgene (#200670-#200681).

### PBMCs and T cells isolation

PBMCs were isolated with Lymphoprep (STEMCELL, #07801) from different donors’ buffy coats obtained from the “Rotkreuz-Organisation für die Menschen im Kanton Bern”, and frozen in FBS (Thermo Scientific, #10270106) with 10% DMSO (Sigma, #D2438). T cells were purified from thawed PBMCs by using EasySep^TM^ Human CD8^+^ T cell Isolation Kit (STEMCELL, #17953) or EasySep Human T cell Isolation Kit (STEMCELL, #17951) following the manufacturer’s instructions, and used for CAR T cell production.

### CAR T cells production

On day 1, 1×10^6^ T cells freshly purified from PBMCs were activated with CD3/CD28 T Cell Activators (STEMCELL, #10971). T cells were expanded for 3 days in XF Expansion medium (STEMCELL, #10981) with addition of 10 ng/ml IL-2 (STEMCELL, #78036.2) and 10 ng/ml IL-21 (STEMCELL, #78082). On day 4, T cells were transduced with Lentiviruses at an MOI of 5, as described below. On day 7, GFP^+^ CAR T cells were sorted by FACS and expanded with addition of 10 ng/ml IL-2, until day 13 for cytotoxicity assays, cytokine production, or *in vivo*. Dual-CAR T cells directed against CD276 and FGFR4 were generated by co-transducing T cells with both CD276- and FGFR4-CARs at the MOI of 2.5 each. The idea was to maintain the same total number of surface CARs expressed on the single antigen-targeting CAR T cells originally transduced at the MOI of 5.

### T cells transduction

250’000 Jurkat cells or purified T cells were seeded onto a 24-well plate and transduced at an MOI of 5 via spin-infection for 1h at 800xg and 32°C in 450 µl RPMI-1640 medium (Gibco, #42401-018) containing 50 µl of 100-fold concentrated lentiviruses. After spin-infection, the cells were diluted 2.5-fold in RPMI-1640 supplemented with 10 ng/ml IL-2 and further incubated for 4h with lentiviruses at 37°C. The next day, the cells were centrifuged and resuspended in fresh cell culture medium. After 48h, transduction efficiency was assessed by Flow Cytometry based on GFP and MycTag expression.

### Bioluminescence killing assays

Bioluminescence killing assays were performed by co-incubating 5’000 iRFP720^+^fLuc^+^ RMS target cells, seeded 24h in advance in bioluminescence 96-well plates (Thermofisher, #136101), with GFP^+^ CAR T cells at the Effector:Target (E:T) ratios of 0.5:1, 1:1, 5:1, and 10:1, in 200 µl total of complete RPMI-1640 medium. Proliferation of RMS cells was monitored for 48h. The supernatant was gently aspirated, and each well was filled with 75µl of fresh medium. The plates were then frozen at −20°C until analysis. The day of the analysis, bioluminescence released by RMS target cells was measured by using ONE-Glo^TM^ Luciferase assay system (Promega, #E6120) with GLO-Max device (Promega). The results were analyzed with Microsoft Excel and GraphPad Prism 9.

### Western blotting (WB)

Cells were detached with a scraper, resuspended in PBS and pelleted by centrifugation at 10’000xg for 3 min, 4°C. Pellets were lysed in RIPA buffer (Thermofisher, #89900) with addition of Halt™ Protease Inhibitor Cocktail (100x) (Thermofisher, #78430) for 20 min on ice. Cells were further homogenized with 1 min sonication (10% duty cycle, Branson Sonifer 250). Protein concentration was measured by Pierce BCA Protein Assay Kit (Thermofisher, #23225). Loading buffer (Invitrogen, #NP0007) was added to 30 μg of protein for each sample. After heating at 65°C for 10 min, samples were loaded on a SurePAGE^TM^, Bis-Tris, 10×8, 4-12% gel (GenScript, #M00654) and separated by MOPS buffer (GenScript, #M0067) at 120V for about 1h. Separated proteins were then transferred to a PVDF membrane in transfer buffer (BioRad, #1610732. 10x: 250 mM Tris base, 1.92 M glycine, 0.1 % SDS) with 10% methanol, at a constant voltage of 100V at 4°C for 1.5 h. PVDF membrane was blocked by 30 min incubation in TBS-T (TBS 10x: 200mM Tris (Sigma, #1185-53-1), 1.37M NaCl (Sigma, #S9888), pH 7.6; supplemented with Tween 0.1%; (Sigma, #P1379)) and 10% skimmed milk. Membranes were then incubated overnight at 4°C with the corresponding primary antibody in TBS-T. After 1h incubation at room temperature with the secondary antibody, the signal was developed with SuperSignal West Femto Maximum Sensitivity Substrate (Thermofisher, #34094) and measured by using ChemiDoc MP Imaging System (Bio-Rad).

### Enzyme-linked immunosorbent assays

ELISAs were performed to assess the amount of cytokines released by CAR T cells during co-incubation with RMS target cells. 25’000 RMS target cells were seeded 24h in advance in transparent 24-well plates (Greiner bio-One, #662160), and CAR T cells were co-cultured at the Effector:Target (E:T) ratios of 5:1 in 500µl total volume of complete RPMI-1640 medium. After 24h, the supernatant was collected, centrifuged at 600xg, transferred to a new plate, and stored at −80°C until use. ELISAs were performed to measure the release of IL-2 (Invitrogen, #88-7025-88), IFN-γ (Invitrogen, #88-7316-88), and Granzyme B (Biozol, #3485-1H-20 for CD276- and CD19-CAR T cells; MabTech, #3486-1H-6 for F8-FR4-CAR T cells) following the manufactures’ instructions. The results were analyzed with Microsoft Excel. The figures were generated using GraphPad Prism 9.

### Antibodies

**Western Blotting** primary antibodies and working dilutions used were: goat anti-CD276 (1 µg/ml; R&D systems, #AF1027); mouse anti-α-tubulin (1:1’000; Cell Signaling, #2144); rabbit anti-MycTag (1:1’000; Cell Signaling, #2278S); mouse anti-human FGFR4 (Santa Cruz, #SC-136988). Secondary antibodies: anti-goat IgG, HRP-linked (1:10’000; Cell Signaling, #HAF109); anti-mouse IgG, HRP-linked (1:10’000; Cell Signaling, #7076); anti-rabbit IgG HRP-linked (1:10’000; Cell Signaling, #7074P2). **Flow Cytometry** primary antibodies and working dilutions: BV421-CD19 antibody (BioLegend, #115537); PE-CD276 (1:100; R&D, #AF1027); PE-FGFR4 (1:100; Biolegend, #324306); CARs expression on CAR T cells was assessed with PE-MycTag (1:100; Cell Signaling, #64025S). CD276 CARs expression on CAR T cells was assessed with human B7H3-Fc chimeric protein (20µg/ml; R&D, #1027-B3) and Goat αHuman IgG (H+L)-Alexa Fluor 647 Ab (1:100; Jackson ImmunoResearch, #109-606-088). CD8/CD4 T cells ratio and activation was assessed with BD Multitest CD3/CD8/CD45/CD4 (1:20; BD Biosciences, #342317) and APC-CD69 (1:100; BD Biosciences, #555533). BV421-CD62L (1:100; BioLegend, #304828); BV510-CD3 (1:100; BioLegend, #563109); BV650-CCR7 (1:100; BioLegend, #353234); APC-CCR5 (1:25; BioLegend, #556903); PerCP-CD4 (1:100; BioLegend, #300528); AF700-CD8 (1:500; BioLegend, #301028); PE-CD95 (FAS) (1:100; BioLegend, #305608); PE-Cy7-CD45RA (1:100; BioLegend, #304108); BV421-CD39 (1:100; Biolegend, #328214); PE-LAG3 (1:100; Biolegend, #369206), PECy7-PD1 (1:100; Biolegend, #329918) and APC-TIM3 (1:100; Biolegend, #345012) were used for phenotype and exhaustion analysis. UltraComp eBeads™ compensation beads (Thermofisher, #01-2222-41) were used for compensation. CD276 and FGFR4 surface copy numbers was calculated with the PE Fluorescence Quantitation Kit (BD Biosciences, #340495) according to the manufacturer’s protocol. **Immunohistochemistry primary** antibodies and working dilutions: mouse monoclonal anti-human FGFR4 (1:50; Santa Cruz, #SC-136988), goat polyclonal anti-CD276 (1 µg/ml; R&D systems, #AF1027), rabbit monoclonal anti-CD3 (1:200; Abcam, #16669). **Immunohistochemistry secondary** antibodies and working dilutions: polyclonal goat anti-mouse (DAKO, #P0447), polyclonal rabbit anti-goat (1:50; DAKO, #P016002), polyclonal goat anti-rabbit (1:100; DAKO, #P0448). The protein expression was detected using the liquid DAB^+^ substrate chromogen system (DAKO, #K3468).

### Immunohistochemistry

Immunohistochemistry analyses were performed on freshly cut, formalin-fixed, and paraffin-embedded tissue slides. Briefly, deparaffinized and rehydrated tissues were treated with 10mM sodium citrate buffer (pH 6) for antigen retrieval. The samples were next blocked with 10% goat or rabbit serum (depending on the host) and 1% BSA for 1h, before being incubated overnight at 4°C with the corresponding primary antibody. Endogenous peroxidase activity was blocked with peroxidase block (DAKO, #K1492). The tissues were incubated with the corresponding secondary antibody for 1h at room temperature, tissues were counterstained in hematoxylin, dehydrated and mounted with Eukitt mounting media (Sigma-Aldrich, #03989). Images were captured with Leica Microsystem DM6000 B microscope.

### Flow Cytometry

Cell staining was performed in 100 µL FACS buffer (PBS + 2% BSA), by diluting the corresponding primary antibodies following the optimized concentrations described above. Flow cytometry experiments were evaluated with CytoFLEX device (Beckman Coulter). The results were analyzed by FlowJo v10.8.1 Software (BD Life Sciences).

### Mice

NOD.Cg-PrkdcscidlL2rgtm1Wjl/SzJ (NSG) (4-6 weeks old) immunodeficient female mice were purchased from Charles River (Germany) and housed at the Central Animal Facility of the University of Bern, under pathogen-free conditions. The experiments were approved by Animal Welfare Office of Canton Bern and in accordance with Swiss Federal Law (Authorization BE22/2022). After 2 weeks of housing, mice were injected in the left gastrocnemius muscle with 50μl of iRFP720^+^fLuc^+^ Rh4, RD, or JR cells (5×10^5^ cells) in sterile PBS. Four days after the injection of iRFP720^+^fLuc^+^ RMS cells, the size of tumors was estimated by bioluminescence imaging with the In Vivo Imaging System (IVIS). On day 5, 5×10^6^ CAR T cells per mouse were resuspended in 100 µl sterile PBS and intravenously injected into the tail vein. Tumor size was measured by using the following formula: Volume = ((short length)^2^ * (long length))/2.

### Single cells isolation from tumors

Expression of CD276 and FGFR4 on RMS cells derived from xenografted tumors was evaluated by Flow Cytometry. Tumors were chopped and incubated in Balanced Salt Solution (HBSS; Thermofisher, #88284) with addition of collagenase III (200 U/ml, corresponding to1 mg/ml, STEMCELL, #07422) and DNAse I (200 U/ml, Thermofisher, #18047019) for 1 h, 37°C. Cells were filtered with 40 µm-strainer and red blood cells were lysed by incubation with ACK lysis buffer (10x: 1.5M NH_4_Cl (Sigma, #213330), 0.1M KHCO_3_ (Sigma, #237205), 0.001M Disodium EDTA (Thermofisher, #AM9260G) for 5 min, room temperature. Samples were then centrifuged at 400xg for 5 min and washed twice before following the Flow Cytometry protocols as described above.

### Statistical analysis

All statistical analyses were conducted using GraphPad Prism 9, incorporating a two-way ANOVA to examine main effects and interactions between variables. The statistical significance of the differences between experimental and control groups was assessed using Dunnett’s multiple comparison test following a two-way ANOVA with levels of significance indicated as follows: p>0.05 (ns), p≤0.05 (*), p≤0.01 (**), p≤0.001 (***), p≤0.0001 (****).

## Results

### Expression of CD276 and FGFR4 in RMS cell lines and PDXs

CD276-CAR T cells have shown excellent pre-clinical activity against several pediatric tumors but have not been tested against RMS (70). In order to verify CD276 expression levels in RMS, we selected six FP-RMS cell lines (JR, RMS, Rh4, Rh5, Rh28, Rh30), five FN-RMS cell lines (RD, Rh18, Rh36, TTC-442, RUCH-3), and three PDXs (IC-pPDX-104, IC-pPDX-35, RMS-ZH003), and quantified total CD276 expression by western blotting (WB) (Fig. 1A) and cell surface expression by FACS (Fig. 1B-E). WB analysis revealed high total expression levels of CD276 in RMS PDXs, and in most RMS cell lines, with the exception of JR, RMS, and Rh18, where lower expression levels were observed. WB confirmed low expression in control myoblasts, PBMCs, and T cells (Fig. 1A). Flow Cytometry revealed high surface expression of CD276 in all the samples (Fig. 1B and 1D), low expression on myoblasts, and no detectable expression in PBMCs and T cells controls. Interestingly, lower levels of surface CD276 were observed for JR, RMS, and Rh18, in agreement with WB results. The number of CD276 on the cell surface were estimated with PE-beads (Fig. 1C, 1E). The most representative RMS cell lines RD, Rh30, and Rh4, displayed around 50’000 copies on their surface. Consistently with the previous analyses, JR, RMS, and Rh18 displayed the lowest number of copies of CD276 on the surface, with ca. 8’000, 10’500, and 25’000 copies, respectively (Fig. 1C). These experiments confirmed consistent expression of CD276 on RMS cell lines and PDXs supporting their targeting by CD276-CAR T cells.

**Fig. 1.**
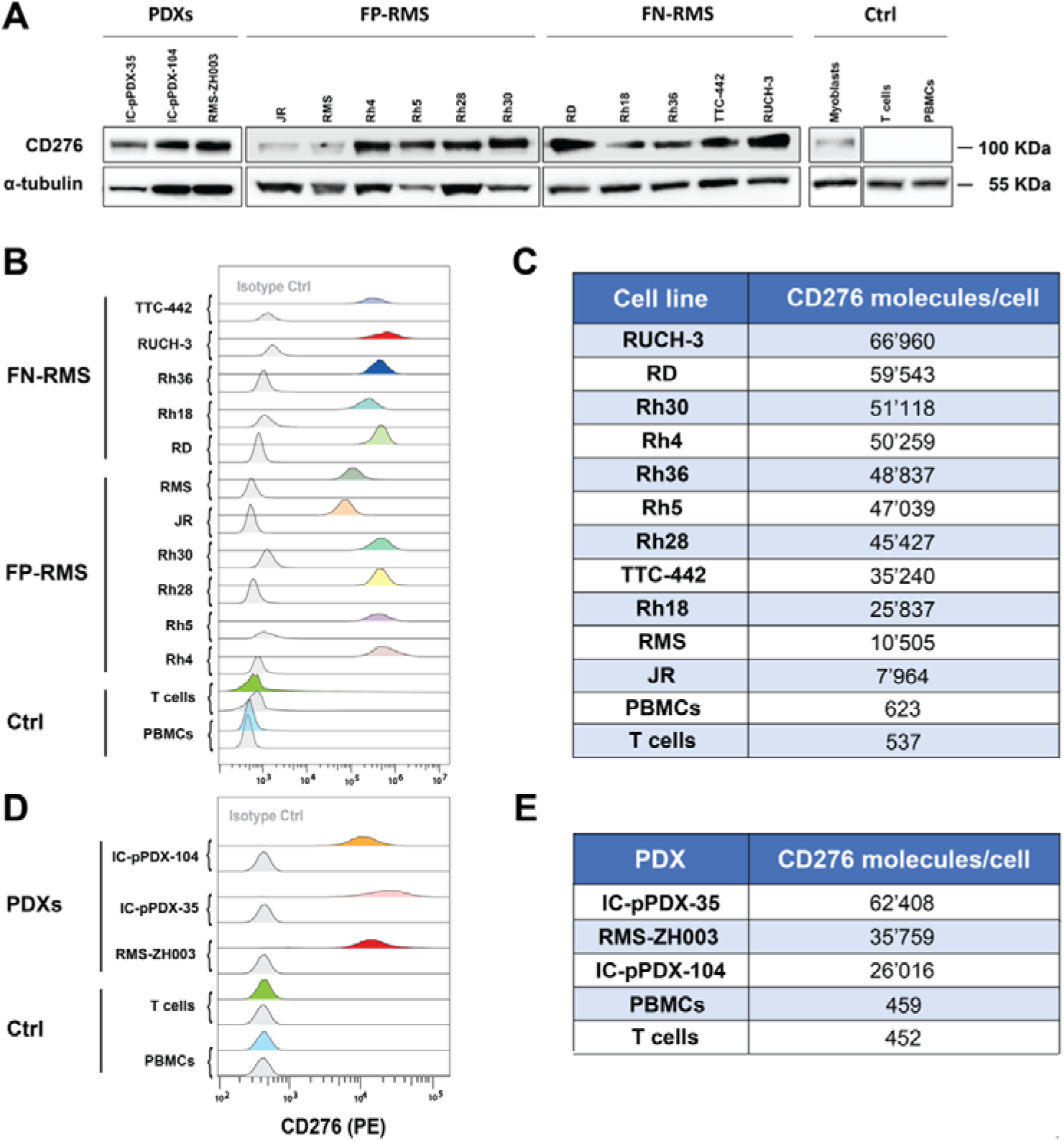
CD276 expression in RMS cell lines and PDXs. (**A**) CD276 expression levels were evaluated in eleven RMS cell lines, three PDXs, myoblasts, PBMCs, and isolated T cells by Western Blotting, revealing high expression in all RMS cell lines, except for the FP-RMS JR, Rh18, and RMS, medium-high levels in the three investigated PDXs, and no detection in the controls. (**B**) Surface CD276 expression on eleven RMS cell lines, and myoblasts, PBMCs, and T cells as controls, was assessed by Flow Cytometry confirming high expression of CD276 in all the considered RMS samples, especially in RUCH-3, RD and Rh4, and no detectable expression in the controls. The correspondent isotype controls are shaded in light grey. (**C**) The number of CD276 molecules on the surface of RMS cells, PDXs, and controls was estimated by Quantibrite PE beads. (**D**) Surface CD276 expression on three RMS PDXs, and PBMCs and T cells, as controls, was assessed by Flow Cytometry confirming high expression of CD276 in all the three PDXs. The correspondent isotype controls are shaded in light grey. (**E**) The number of CD276 molecules on the surface of RMS PDXs and controls was estimated by Quantibrite PE beads.

Antigen loss is a common mechanism of tumor resistance against CAR T cells that can be addressed by CAR T cells able to recognize multiple targets on the surface of tumor cells. Therefore, we next focused on FGFR4, another promising target for RMS, as also recently confirmed by our RMS surfaceome profiling (32). Expression levels of FGFR4 on RMS cell lines and PDXs were assessed by WB and Flow Cytometry. WB put in evidence high expression levels of FGFR4 in Rh4, Rh28, JR, and RMS cell lines, and in the PDXs RMS-ZH003 and IC-pPDX-104; medium FGFR4 expression levels were instead observed in Rh36, RD, and TTC-442; whereas low levels of FGFR4 were detected in Rh18, RUCH-3, Rh5, and Rh30 cell lines, and in IC-pPDX-35 (Fig. 2A). Flow Cytometry analysis performed on live cells confirmed expression of FGFR4 on most of the RMS cell lines (Fig. 2B). No expression of FGFR4 could be detected on PDXs. Quantification of surface FGFR4 copy number revealed a relatively low copy number of FGFR4 on RMS cells (Fig. 2C), compared to CD276. Despite the lower density of FGFR4 on RMS samples compared to CD276, the absence of expression in most normal tissues makes FGFR4 a promising target in RMS for CAR T cell-based strategies (71–73).

**Fig. 2.**
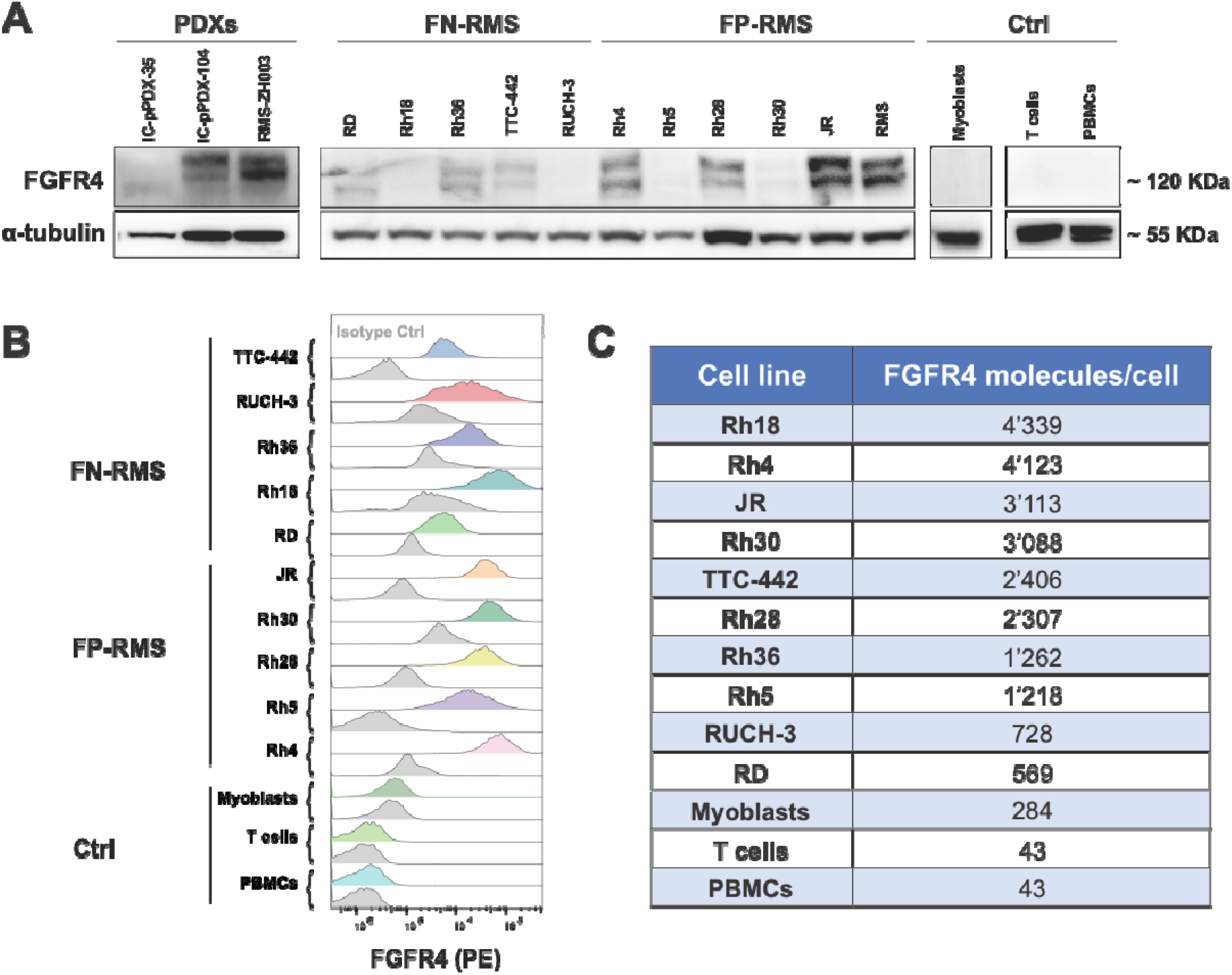
FGFR4 expression in RMS cell lines and PDXs. **(A)** FGFR4 expression levels were evaluated in eleven RMS cell lines, three PDXs, and in the controls, such as Myoblasts, PBMCs, and T cells. WB results revealed high expression in Rh36, Rh4, Rh28, JR, and RMS cell lines, whereas low levels were detected in Rh18, RUCH-3, Rh5, and Rh30. No FGFR4 expression was observed in the controls. (**B**) Flow Cytometry results confirmed expression of FGFR4 on the surface of most RMS cell lines, especially high in Rh4, Rh18, and JR. The corresponding isotype controls are highlighted in grey. (**C**) Estimation of FGFR4 expression on RMS cell lines by Quantibrite PE-Beads showed a surface expression in the range of 600-4000 copies per cell.

### Selection of FGFR4 Antigen Binding Domain

We previously selected four FGFR4-targeting single-domain antibodies (sdAb) A8-FR4, B1-FR4, B5-FR4, and F8-FR4 (71). These were used as ABD to generate lentiviral CAR plasmids and to produce the corresponding CAR T cells (Fig. 3A). After sorting and expansion, T cells expressed the sdAb-FR4-CARs to a similar extent, as monitored by GFP expression and MycTag staining (Fig. 3B). To determine the killing capacity of sdAb-FR4-CAR T cells and to screen for the most specific and effective construct, we performed cytotoxicity assays with fLuc^+^ Rh4 cells. Rh4 were chosen among a panel of RMS cell lines due to the high surface expression of FGFR4 with around 4’000 FGFR4 copies per cell (Fig. 2C). In order to estimate the specificity of the different sdAb, FGFR4-KO Rh4 cells were generated by CRISPR/Cas9 (Fig. S1) and were used as negative controls in cytotoxicity assays. All four sdAb-FR4-CAR constructs were able to trigger more than 80% tumor cell killing by CAR T cells at E:T ratios of 5:1 and 10:1 (Fig. 3C). However, at lower ratios of 0.5:1 and 1:1, B5-FR4-CAR T cells showed the highest cytotoxic effect by eliminating C 40% and C 60% of Rh4 cells, respectively (p<0.001, see Suppl. Table S4). A8-FR4- and F8-FR4-CAR T cells showed a killing capacity of C 20% and C 30% respectively, whereas B1-FR4 was the weakest in terms of cytotoxicity, with no killing at 0.5:1 (Fig. 3C). On the other hand, co-incubation of FGFR4-CAR T cells with fLuc^+^ FGFR4 KO Rh4 cells surprisingly revealed an unspecific activity of CARs A8-FR4, B1-FR4, and B5-FR4, resulting in an effective killing of FGFR4 KO Rh4. F8-FR4 was the only CAR showing no activity against FGFR4 KO Rh4 cells (Fig. 3D). Therefore, F8-FR4 was selected to arm FGFR4-CAR T cells for the next *in vitro* and *in vivo* experiments.

**Fig. 3.**
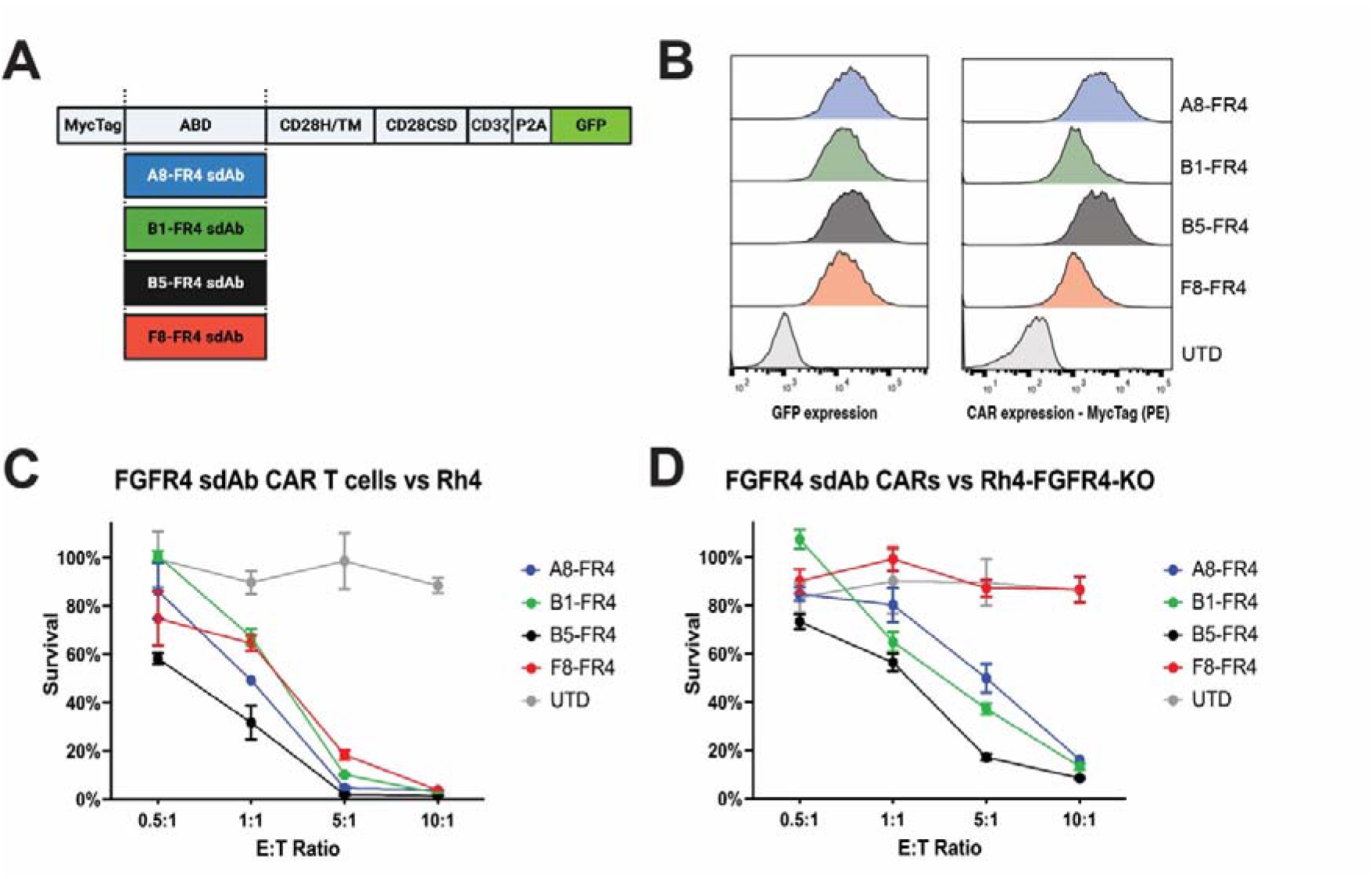
Selection of FGFR4-targeting antigen binding domain. **(A)** Schematic description of the sdAb-FR4-CAR constructs. Created with Biorender.com. (**B**) To select the FGFR4-targeting sdAb that confers the most specific and potent killing capacity to the CAR construct, four different sdAb-FR4-CAR T cells were generated by lentiviral transduction of activated T cells, sorted by GFP expression, and expanded. Expression of CAR constructs was verified by MycTag staining and GFP expression. (**C**) sdAb-FR4-CAR T cells were co-incubated for 48h with fLuc^+^ Rh4 cells at different E:T ratios. RMS cells survival measured by luciferase assays showed effective killing capacity by all the FGFR4-CAR T cells at E:T ratios of 5:1 and 10:1 (p<0.0001, determined using Dunnett’s multiple comparison test following a two-way ANOVA, see Suppl. Table S4). Low E:T ratios put in evidence B5-FR4 sdAb-based CAR as the most effective. (**D**) To verify the specificity of the sdAb-FR4-CARs, CAR T cells were incubated with fLuc^+^ FGFR4 KO Rh4 cells. These experiments showed unspecific killing of the A8-FR4-, B1-FR4-, and B5-FR4-CAR T cells. Nevertheless, F8-FR4-CAR T cells did not kill fLuc^+^ FGFR4 KO Rh4 cells.

### Generation and validation of CD276- and FGFR4-CAR constructs

Aiming at comparing different CAR constructs to identify the CAR conferring the highest cytotoxicity to CAR T cells targeting RMS, we generated a panel of second- and third-generation CARs with HD and TM deriving from either CD8α or CD28, with intracellular CSDs from 4-1BB and/or CD28, and with CD3ζ as SD, as schematically summarized in Fig. 4A. By Gibson assembly, we initially produced eight modular CAR plasmids with the CD19-targeting scFv FMC63 used in the clinic and regarded as the golden standard (CD19.I-VIII, Fig. 4B). These eight CD19-CAR plasmids were then used as negative controls, or as positive controls with RMS cells overexpressing a truncated CD19 (tCD19). CD276-CARs and F8-FR4-CARs were generated by swapping the ABD with the CD276.MG scFv sequence (70) and the F8-FR4 sdAb sequence, respectively. All the CAR plasmids used in this study are summarized in Suppl. Table S3. In order to validate the constructs, Jurkat T cells were initially transduced with lentiviruses coding for CD19- and CD276-CAR constructs, and both correct size and surface expression of the CARs were assessed on CAR Jurkat T cells by WB and Flow Cytometry, respectively. Anti-MycTag detection revealed high expression levels at the expected size for all the CARs (ca. 60 kDa), showing a clear shift from second- to third-generation CARs, due to the additional CSD (Fig. S2A). Flow Cytometry on live cells confirmed high detection of MycTag on the surface of CAR Jurkat T cells, thereby further validating our construct design (Fig. S2B).

**Fig. 4.**
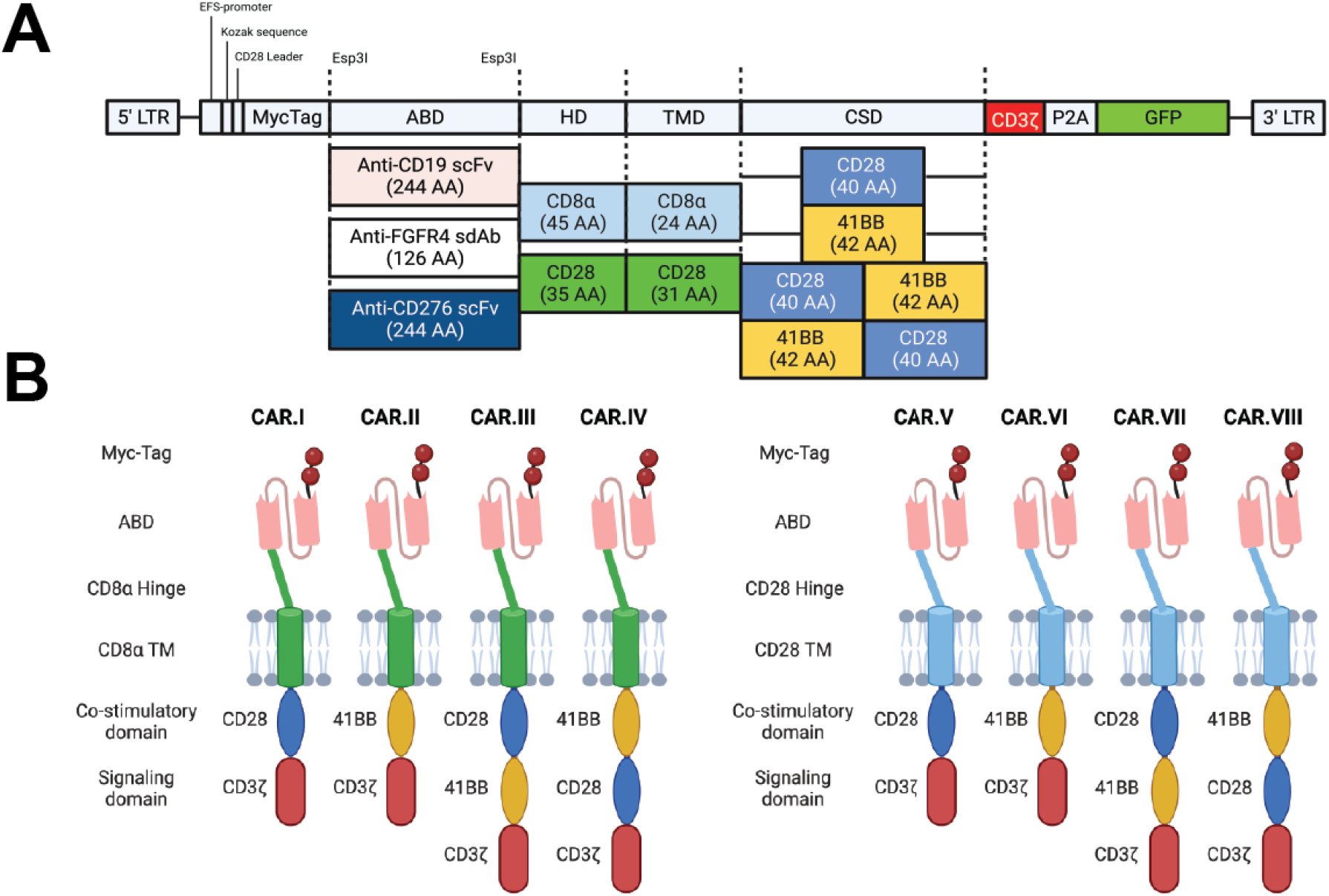
Schematic representation of the investigated CAR constructs. **(A)** The backbone used for the expression of the CARs is characterized by an EFS promoter, followed by a Kozak sequence and a CD28 leader sequence. The ORF is composed of two MycTag regions; an Antigen Binding Domain (ABD; anti-CD19 scFv FMC63 for the initial cloning) flanked by two Esp3I restriction sites for ABD swapping with anti-CD276 scFv (CD276.MG) or with FGFR4-targeting F8-FR4 sdAbs; a Hinge Domain (HD) and a Transmembrane domain (TM) derived from either CD8α or CD28; one or two Co-Stimulatory Domains (CSDs), derived from CD28 or 4-1BB; and a CD3ζ Signaling Domain (SD). Downstream, GFP as reporter is linked to the CD3ζ domain by the self-cleaving peptide P2A. (**B**) Modular CAR structure of the different constructs generated by Gibson assembly with either CD8α HD/TM (green) or CD28 HD/TM (cyan). In each panel, the schematic representation of two second-generation CARs, characterized by a single CSD (CD28 or 4-1BB), and two third-generation CARs, characterized by the expression of both CSDs (CD28-4-1BB or 4-1BB-CD28). Created with Biorender.com.

### Manufacturing and characterization of CD276- and F8-FR4-CAR T cells

T cells isolated from PBMCs were seeded on day 1 in XF expansion medium, supplemented with IL-2, IL-21, and CD3/CD28 activators, as schematically represented in Fig. 5A. Expansion and viability of T cells, and CD4^+^:CD8^+^ ratios were monitored for 14 days prior to both *in vitro* and *in vivo* experiments. Initially, activation of T cells was monitored over one week by measuring CD69 expression. The highest levels of CD69 expression were observed four days after activation (Fig. S3A). T cells were therefore transduced with lentiviruses on day 4 and the efficiency was measured after 48h by GFP and MycTag detection by FACS. Infection efficiency was over 40% for most of the CD276 and F8-FR4 constructs with a mean of 65.5% ± 7.5% by GFP expression and 79.9% ± 16.2% by MycTag expression (Suppl. Table S5, S8). In order to normalize the expression of the CAR constructs, and to allow a precise comparison of their activity, CAR T cells were then sorted by GFP on day 7. IL-2 and CD3/CD28 activators were added fresh, and after further 7 days in culture, the CAR T cells showed 26-fold to 35-fold *in vitro* expansion and a viability of more than 70% (Fig. 5B, 5D). No significant difference in expansion was observed between the different CAR constructs with UTD T cells. At the moment of the co-incubation with target cells, the CD4^+^:CD8^+^ ratio of the amplified CAR T cells was around 1:1 (Fig. 5C, 5E) with a mean of 45% ± 3.7% CD4^+^ and 53.7% ±3.8% CD8^+^ T cells (Suppl. Table S6). No significant difference was observed between the different CAR constructs. CAR size and expression on the surface of CAR T cells were assessed by WB and Flow Cytometry based on MycTag detection. This confirmed the correct expression at ca. 60kDa and difference in size between the second- and third-generation constructs for CD19-CARs (Fig. 6A), CD276-CARs (Fig. 6C), and F8-FR4-CARs (Fig.6E). High surface CAR expression levels were observed in all the samples (Fig. 6B, 6D, 6F). All the experiments were performed with three different donors on day 14 (Suppl. Table S7).

**Fig. 5.**
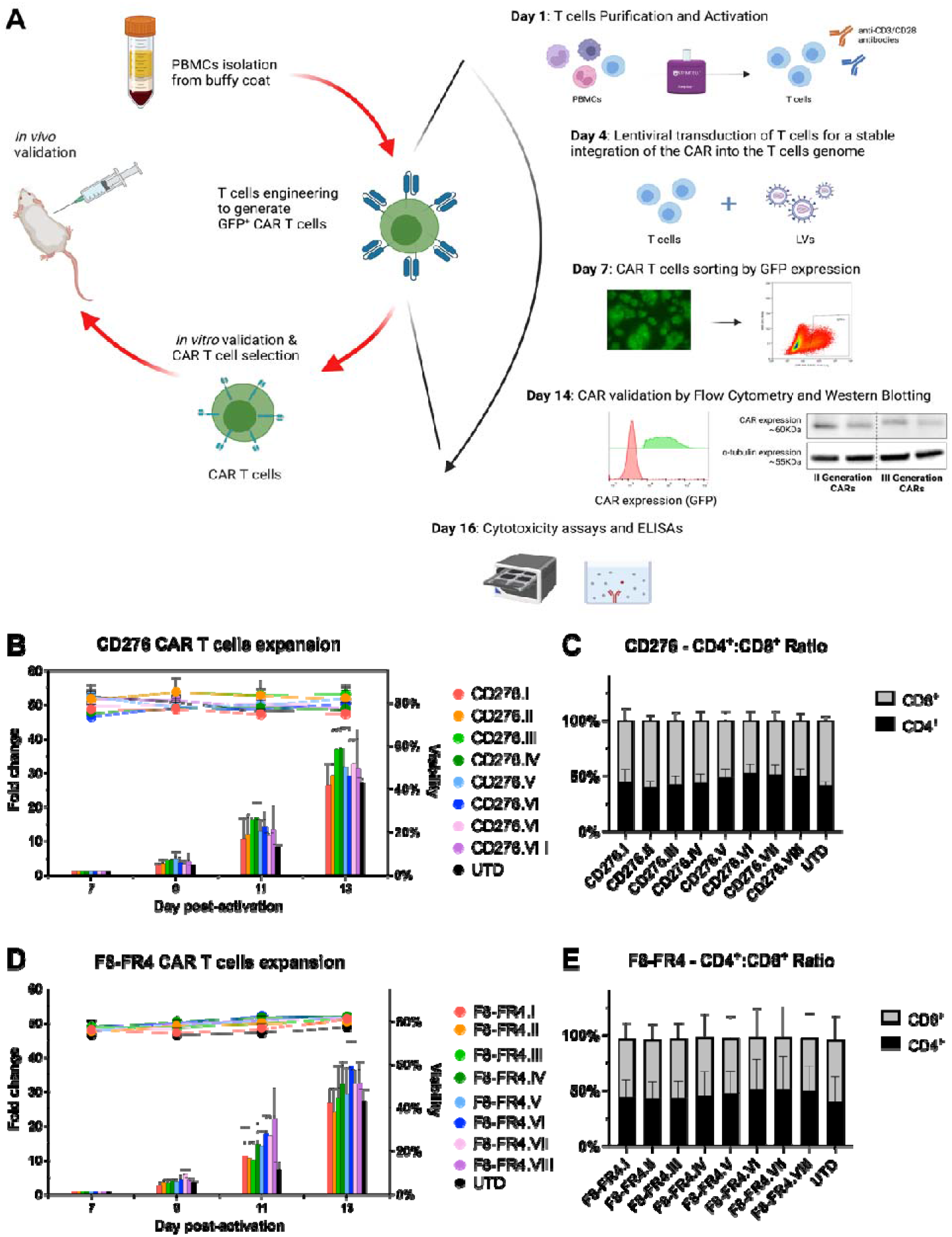
Manufacturing, expansion, viability, and CD4^+^:CD8^+^ ratios of CD276- and F8-FR4-CAR T cells. **(A)** Manufacturing, selection, and testing of CAR T cells. After isolation from PBMCs, T cells were incubated on day 1 with IL-2, IL-21, and anti-CD3/CD28 activators. On day 4, T cells were infected with lentiviruses, CAR T cells were then sorted by GFP expression on day 7 and further expanded with addition of IL-2 for 7 days. Surface CAR expression before cytotoxicity assays were then tested by WB and FACS. On day 14, CAR T cells were co-incubated with RMS cells for killing assays and cytokine release quantification. (**B**) Expansion and viability of CAR T cells were monitored from day 7 to day 13, resulting in up to 35-fold expansion, with a viability >70%. Data shown for three donors. (**C**) CD4^+^:CD8^+^ ratio was assessed on day 14 to rely on a balanced number of helper and cytotoxic T cells. In all the CAR T cells the CD4^+^:CD8^+^ ratio was around 1:1. Data shown for three donors (see Suppl. Table S6). Created with BioRender.com.

**Fig. 6.**
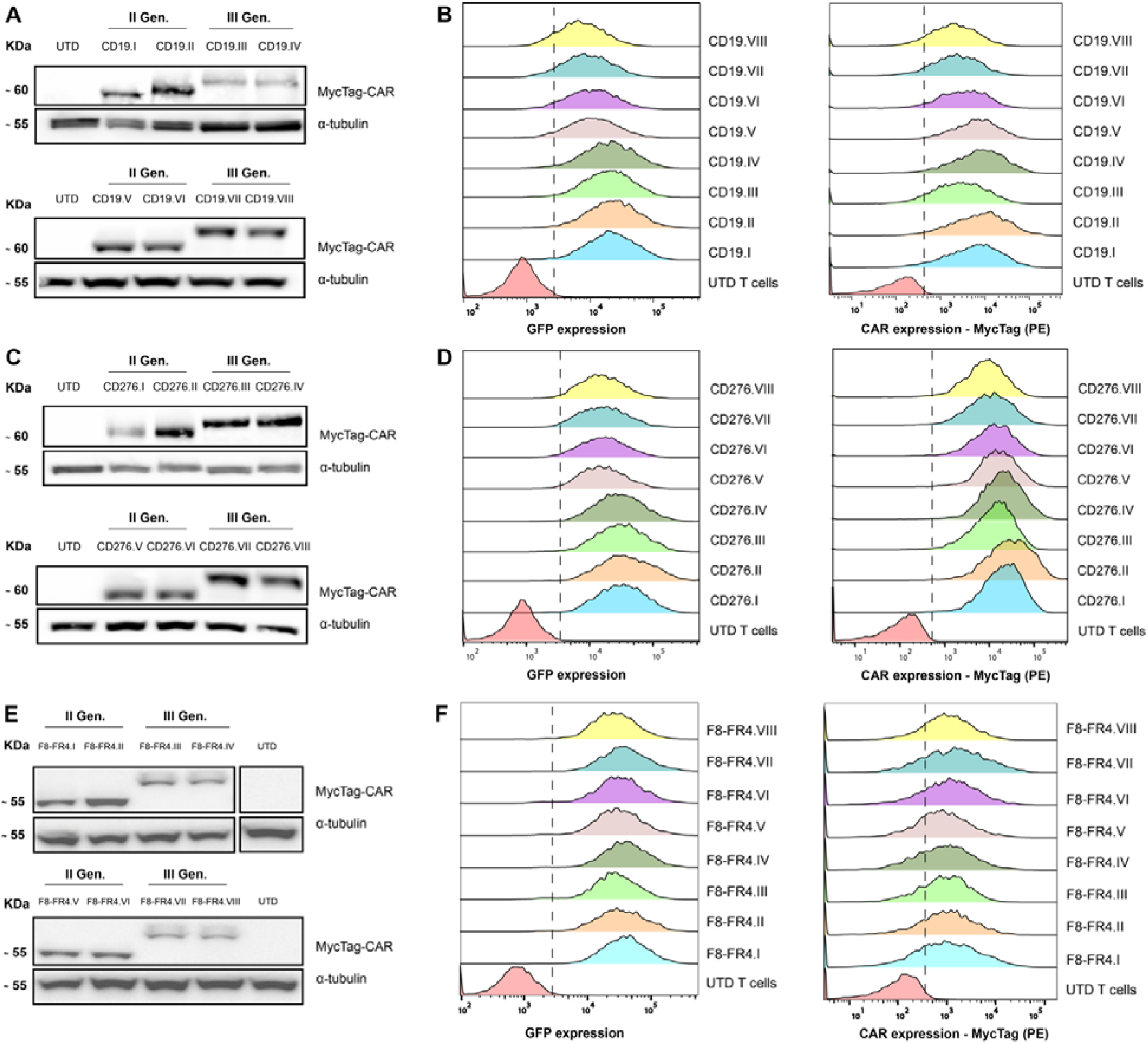
Validation of the CAR constructs expression in T cells. CAR expression in T cells was assessed by Western Blotting (**A, C, E**) and by Flow Cytometry (**B, D, F**) on day 14, after first activation of T cells for CD19-CAR (A, B), CD276-CAR (C, D), and F8-FR4-CAR (E, F). Anti-MycTag antibody was used to discriminate the different CAR generations by size in Western Blotting. The respective bands confirm the predicted CAR size around 60 kDa with a visible shift between the second- and third-generation CARs. (**B**, **D, F**) Surface expression of CAR constructs was detected with an anti-MycTag PE-conjugated antibody by Flow Cytometry, confirming robust expression of the CAR constructs on the surface of T cells. All the experiments were performed in triplicates and with three different donors. Shown are results from one representative donor. Quantification of CAR expression for each donor is summarized in Suppl. Table S7.

### Cytotoxicity and cytokines release of CD276- and F8-FR4-CAR T cells

All the CAR constructs were then tested to determine the CAR constructs promoting the highest cytotoxicity *in vitro*. CD276-CAR T cells were incubated with fLuc^+^ RD and Rh4 cells, both expressing high levels of CD276 on their surface, while F8-FR4-CAR T cells were incubated with fLuc^+^ JR and Rh4 cells, both expressing high levels of FGFR4 on their surface. After 48h co-incubation, survival of RMS cells was determined by luciferase assays. At the effector to target (E:T) ratios of 10:1 and 5:1, all the CD276 experimental groups efficiently killed almost 100% of fLuc^+^ RMS cells (Fig. 7A, 7B). At a lower E:T ratio of 0.5:1 CD276.I (8.8.28.3z, red) and CD276.V (28.28.28.3z, light blue) showed the highest and significant efficacy, killing 43.0% ± 6.7% and 42.7% ± 7.2% of fLuc^+^ Rh4 cells, respectively; and killing 28.9% ± 5.6% (CD276.I) and 39.2% ± 4.7% (CD276.V) of fLuc^+^ RD cells, respectively. Among the FGFR4-CAR experimental group (Fig. 7C, 7D), F8-FR4.I (8.8.28.3z, red), F8-FR4.IV (8.8.BB/28.3z, dark green), and F8-FR4.V (28.28.28, light blue) showed the highest activity, killing more than 95% of JR and Rh4 cells at an E:T ratio of 10:1. All the constructs showed a significant higher activity compared to untransduced T cells (UTD, black). At the E:T ratio of 0.5:1 the same constructs showed a high and significant efficacy against fLuc^+^ JR cells, the highest being F8-FR4.V with 50.7% ± 7.7%, followed by F8-FR4.I with 35.8% ± 6.8%, and F8-FR4.IV with 27.7% ± 15.1%. CD19-CAR T cells, used here as a negative control, showed no significant killing capacity against fLuc^+^ RD and Rh4 cells (Fig. 7E, 7F), whereas when co-incubated with RMS cells expressing truncated CD19 (tCD19^+^ fLuc^+^ RD and Rh4 cells), CD19.V-CAR T cells specifically eliminated most of RMS cells with great efficacy (Fig.7E, 7F, grey). All experiments were performed in triplicates and with three different donors. Donor specific results are shown in Suppl. Fig. S4, and detailed statistical analysis are described in Suppl. Table S9. These results highlight the CAR backbones 8.8.28.3z (I) and 28.28.28.3z (V) as the most effective in killing RMS cells.

**Fig. 7.**
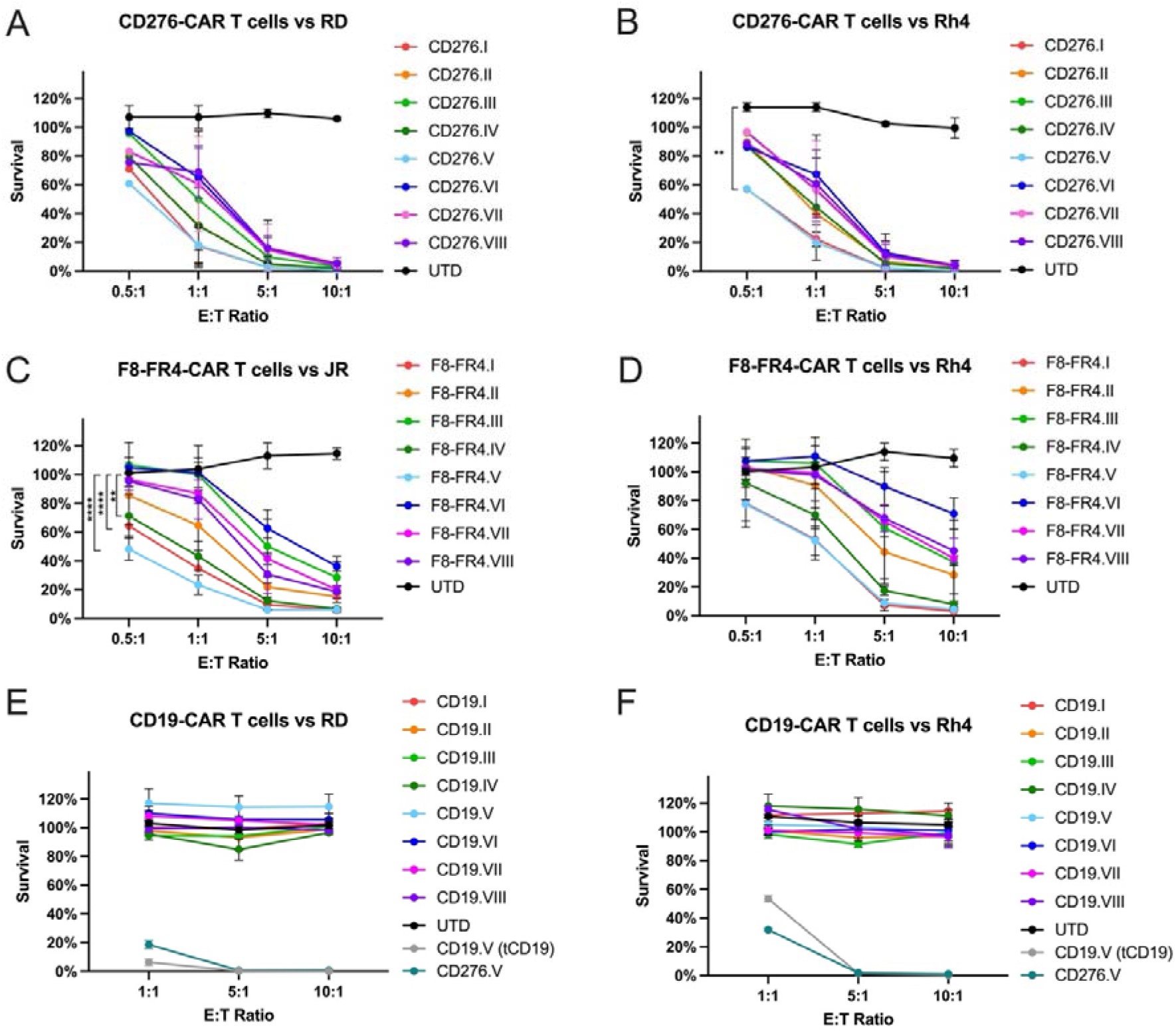
Evaluation of killing capacity by CD276- and F8-FR4-directed CAR T cells after co-incubation with fLuc^+^ RD, Rh4, and JR cell lines. In order to determine the CAR construct conferring the highest cytotoxicity against RMS, CD276-CAR T cells were co-incubated for 48h with fLuc^+^ cells RD (**A**) and Rh4 (**B**) at different E:T ratios. F8-FR4-CAR T cells were co-incubated with fLuc^+^ JR (**C**) and Rh4 (**D**) since RD cells express very low amounts of FGFR4. Luciferase assays showed significant killing capacity by all the generations of CAR T cells at E:T ratios of 5:1 and 10:1, compared to untransduced T cells (UTD) (p<0.0001 for all constructs). Lower E:T ratios highlighted optimal efficacy by CD276.V- and F8.V-CAR T cells, compared to the other constructs. CD19-CAR T cells showed no cytotoxic effect when co-incubated with fLuc^+^ RD (**E**) and Rh4 (**F**) cells. As positive controls, CD19.V-CAR T cells were incubated with fLuc^+^ RD-tCD19 (**E**, grey) and Rh4-tCD19 (**F**, grey) cells. All experiments were performed in triplicates and with three different donors. Donor specific results and detailed statistical analysis are described in Suppl. Fig. S4 and Suppl. Table S9, respectively. The statistical significance of the differences between experimental and control groups was assessed using Dunnett’s multiple comparison test following a two-way ANOVA; p≤ 0.01 (**), p≤0.0001 (****).

Next, the levels of IL-2, IFN-γ, and Granzyme B were measured to quantify cytokine release from the CAR T cells after 24h co-incubation with RMS cells at a 5:1 E:T ratio (Fig. 8 and Suppl. Fig. S5). For the CD276-CAR experimental group, incubated with RD or Rh4 cells (Fig 8A), the results showed highest expression of IL-2 by CD276.I (38.4 ± 5.4 pg/ml) and CD276.V (39.2 ± 16.1 pg/ml) co-incubated with RD cells, and by CD276.I (36.4 ± 25.5 pg/ml) and CD276.V (35.8 ± 18.1 pg/ml) co-incubated with Rh4 cells (Fig. 8A, upper panels). These results were confirmed also in terms of IFN-γ release, with maximum values released by CD276.V-CAR T cells when co-incubated with RD (270.0 ± 154.3 pg/ml) and Rh4 cells (221.0 ± 129.1 pg/ml) and by CD276.I (RD: 136.7 ± 83.2 pg/ml; Rh4: 150.5 ± 60.1 pg/ml) (Fig. 8A, middle panels). Considering the absolute values, Granzyme B was detected at higher levels than IL-2 and IFN-γ, especially when CD276.V-CAR T cells were co-incubated with RD cells (7039.1 ± 1970.6 pg/ml) followed by CARs CD276.I-IV (around 3300 pg/ml) (Fig. 8A, lower panels). The constructs CD276.II and CD276.V were the most effective ones when co-incubated with Rh4 cells, releasing 3011.3 ± 1285.4 pg/ml and 2569.0 ± 1382.3 pg/ml, respectively. For the F8-FR4-CAR experimental group, incubated with JR or Rh4 cells (Fig. 8B), the results showed highest expression of IL-2 by F8-FR4.I (34.5 ± 0.2 pg/ml) and F8-FR4.V (35.9 ± 0.3 pg/ml) co-incubated with JR cells, and by F8-FR4.I, (34.8 ± 0.3 pg/ml) and F8-FR4.V (35.8 ± 0.5 pg/ml) co-incubated with Rh4 cells (Fig. 8B, upper panels). IFN-γ release was highest when F8-FR4.I-CAR T cells were co-incubated with Rh4 cells (132.1 ± 8.0 pg/ml), followed by F8-FR4.V incubated with JR cells (121.5 ± 2.5 pg/ml) and with Rh4 cells (118.4 ± 4.5 pg/ml) (Fig. 8B, middle panels). Considering the absolute values, Granzyme B was again detected at higher levels than IL-2 and IFN-γ, with most constructs releasing more than 5000 pg/ml (Fig. 8A, lower panels). When considering the average of the three donors, only values for Granzyme B were significantly different. However, when comparing released cytokines from the same donor, the values became significant, namely differences between construct V and I (detailed statistical analysis is provided in Suppl. Table S12).

**Fig. 8.**
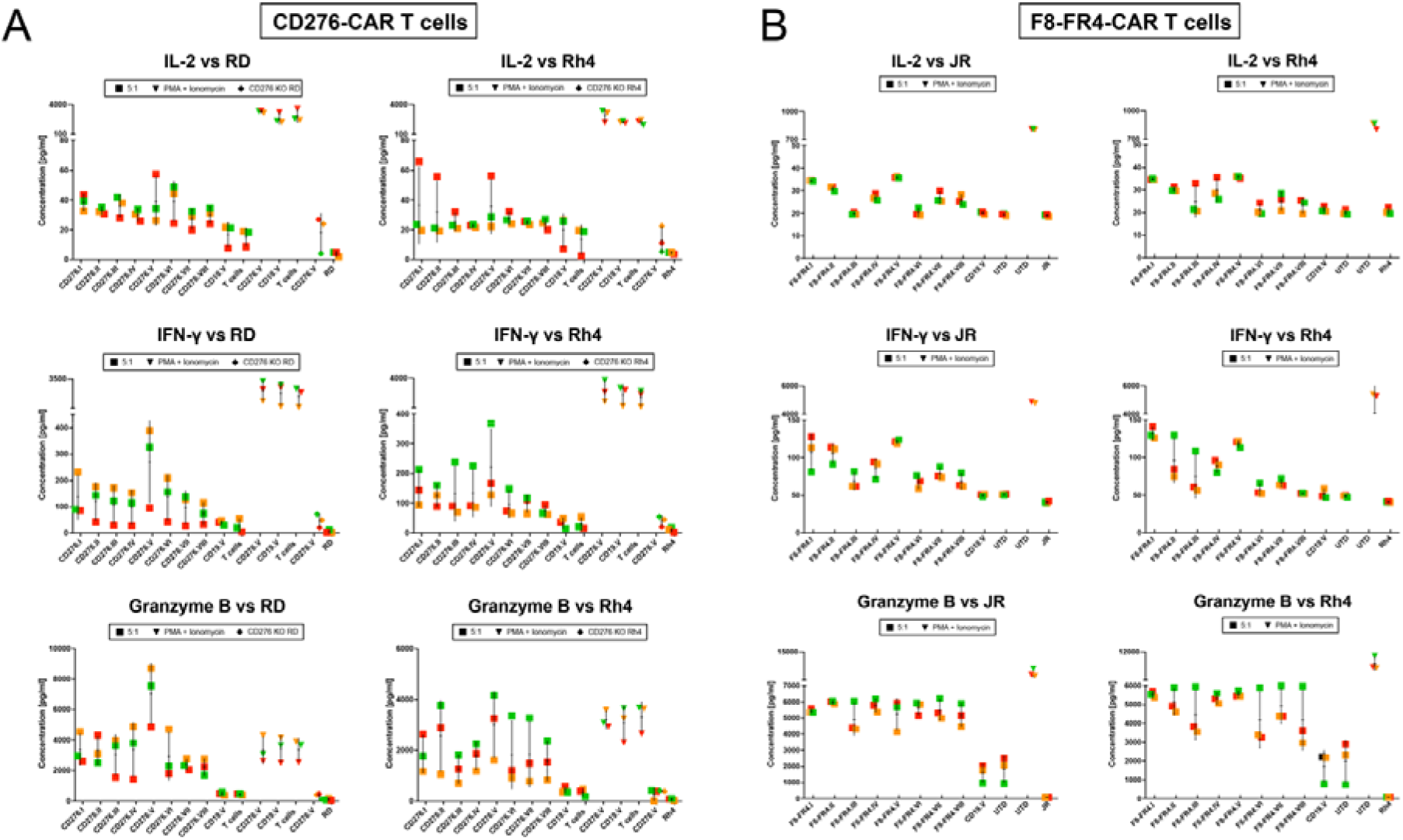
Cytokine release by CD276- and F8-FR4-CAR T cells after 24h co-incubation with RMS cell lines. Concentrations of IL-2, IFN-γ, and Granzyme B in the supernatant released during co-incubation of (**A**) CD276-CAR T cells with RD (left panels) and Rh4 cells (right panels) and (**B**) F8-FR4-CAR T cells with JR (left panels) and Rh4 (right panels) at the E:T ratio of 5:1 were measured by ELISA after 24h. IL-2 measurements revealed CD276.I, CD276.V (**A**), as well as F8-FR4.I, F8-FR4.II and F8-FR4.V (**B**), as the constructs with the highest IL-2 release. CD276.V released maximum levels of IFN-γ with both RD and Rh4 (**A**), while the maximal release of IFN-γ for F8-FR4-CAR T cells was measured for F8-FR4.I (Rh4: 129 pg/ml ± 8 pg/ml) and F8-FR4.V (JR: 121 pg/ml ± 3 pg/ml) (**B**). The highest Granzyme B release was measured during co-incubation of CD276.V with RD cells (7547 pg/ml ± 1970 pg/ml). High values were observed also for CD276.II, CD276.V when co-cultured with Rh4 cells (**A**). For F8-FR4-CAR T cells, the constructs F8-FR4.I, F8-FR4.II and F8-FR4.V showed a robust secretion around 5000 pg/ml with both JR and Rh4 cells. In all the experiments, IL-2, IFN-γ, and Granzyme B levels were at background levels when CD276.V-CAR T cells were co-incubated with CD276 KO RMS cells, similar to background levels detected in RD and Rh4 supernatant alone. UTD: Untransduced T cells. All the experiments were performed in triplicates, and three different donors are shown: donor 1 (red), donor 2 (orange), donor 3 (green).

In conclusion, constructs CD276.V and F8-FR4.V displayed most consistently the strongest killing and highest cytokine release, followed by constructs CD276.I and F8-FR4.I.

### Cytotoxicity of CD276/FGFR4 Dual-CAR T cells against RMS cell lines

Having identified CD276.V and F8-FR4.V as the most effective CAR constructs in terms of cytotoxicity and cytokine release, we generated Dual-CAR T cells directed against CD276 and FGFR4 by co-transducing T cells with both CD276.V and F8-FR4.V CAR lentiviruses. Our aim was to investigate the effect of combined targeting and to develop a strategy to overcome possible antigen loss or antigen escape *in vivo* and to further improve cytotoxicity of the CAR T cells. The quantitative analysis conducted on the experimental CAR T cell groups was performed by measuring GFP and MycTag expression (Fig.9A, 9B), revealing a homogenous population of CAR^+^ cells within samples. More precisely, to discriminate the abundance of CD276-CARs, we relied on the staining with recombinant human CD276-Fc chimeric protein. C 90% CD276-CARs were detected on CD276.V-CAR T cells, whereas only C 35% on Dual.V-CAR T cells, confirming as expected a successful C 2-3-fold reduction in the expression levels. By contrast, no binding of rhCD276-Fc to F8-FR4.V-CAR T cells was observed (Fig.9C).

**Fig. 9.**
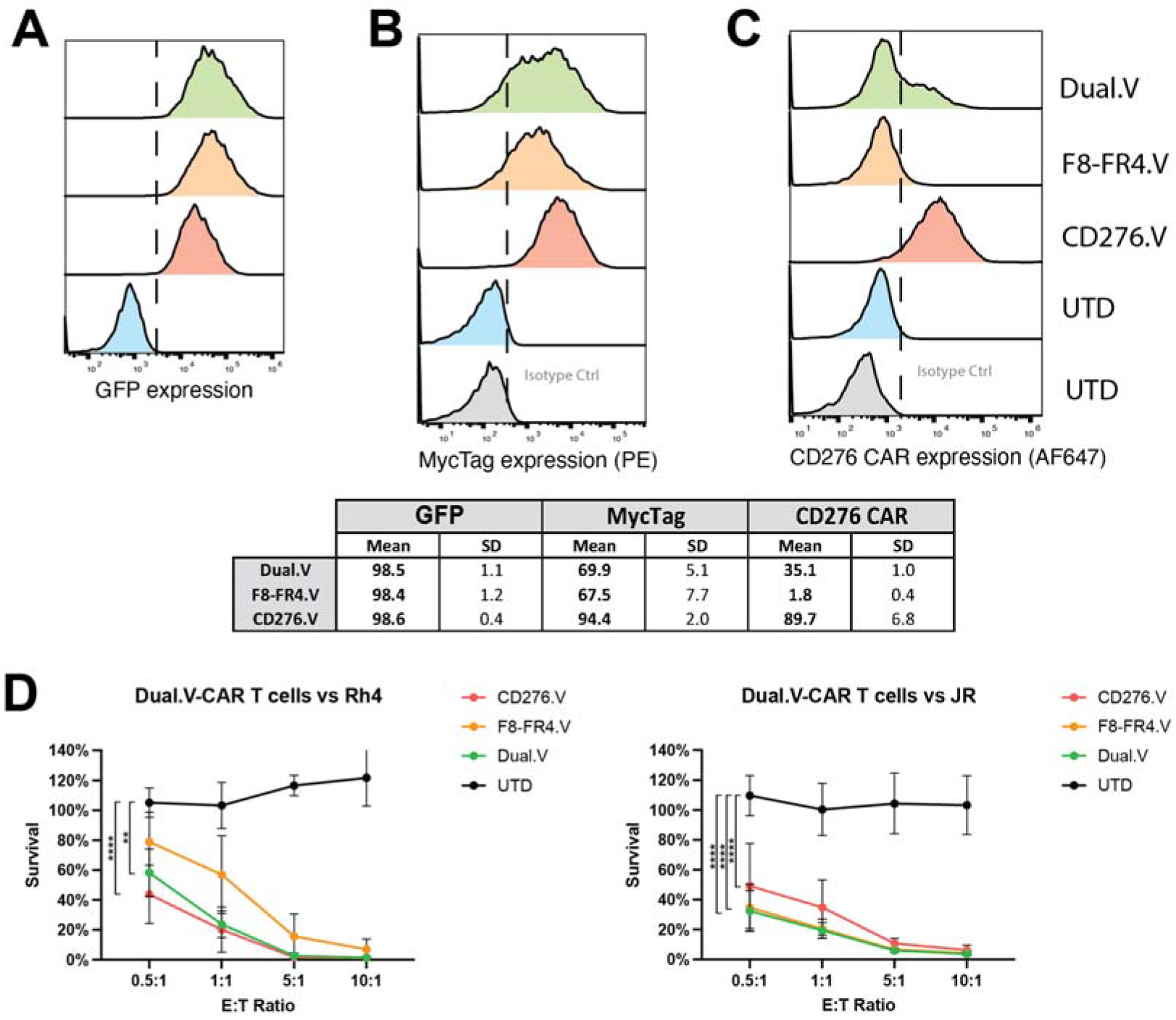
Generation and characterization of Dual.V-CAR T cells killing capacity. Dual.V-CAR T cells were generated by simultaneous transduction of activated T cells with lentiviruses encoding for CD276.V-CAR and F8-FR4.V-CAR. (**A-C**) The quantification of the surface CAR abundance on CD276.V-, F8-FR4.V, and Dual.V-CAR T cells is based on GFP expression (**A**), MycTag expression (**B**), and staining with soluble CD276 (**C**), revealed a significant difference in the CD276.V-CAR detection within the three experimental groups. While the GFP signal shows uniform expression, the staining with the soluble CD276 protein shows 90% of CD276-CAR expression on the CD276.V-CAR T cells, LJ40% on the Dual.V-CAR T cells, and no expression was detected on F8-FR4.V-CAR T cells. (**B**) Killing assays with CD276/FGFR4 Dual.V-CAR T cells were performed with three different donors. The killing capacity of Dual-CAR T cells was evaluated by co-incubation experiments with Rh4 and JR cell lines for 48h at different E:T ratios. Luciferase assays showed effective killing capacity by CD276.V-, F8-FR4.V-, and Dual.V-CAR T cells on Rh4 cells (CD276^high^/FGFR4^high^) at E:T ratios of 5:1 and 10:1. Low E:T ratios exhibited high efficacy by CD276.V- and Dual.V-CAR T cells, compared to F8-FR4.V-CAR T cells (left panel). Killing assays performed with JR cells (CD276^low^/FGFR4^high^) showed high killing capacity by F8-FR4.V- and Dual.V-CAR T cells, but moderate efficacy by CD276.V-CAR T cells (right). Untransduced T cells showed no significant killing against both RMS cells. p≤0.01 (**), p≤0.0001 (****).

CD276.V-, F8-FR4.V-, and Dual.V-CAR T cells were co-incubated for 48h with Rh4 cells, defined as CD276^high^FGFR4^high^; and JR, defined as CD276^low^FGFR4^high^. The results from cytotoxicity assays indicate that, when co-incubated with Rh4, despite the lower CD276-CAR expression, Dual.V-CAR T cells showed a killing efficacy comparable to CD276.V-CAR T cells. By contrast, we observed only a moderate killing of Rh4 cells of around 20% by F8-FR4.V-CAR T cells at the E:T ratio of 0.5:1 (Fig. 9D, left panel). Experiments performed with JR cell line, instead, exhibited a cytotoxic activity of Dual.V-CAR T cells analogous to the F8-FR4.V-CAR T cells, highlighting a highly consistent killing by the Dual.V-CAR T cells regardless of the targeted RMS cell line. As expected, CD276.V-CAR T cells showed a lower activity against JR, likely due to the lower surface expression of CD276 (Fig. 9D, right panel).

### Memory phenotype and exhaustion status of CAR T cells

In order to characterize the CAR T cell subsets enriched during manufacturing and to assess any variation following co-incubation with tumor cells, the T cell phenotype from three different donors was investigated by Flow Cytometry. On day 1, T cells were purified from PBMCs, stimulated with CD3/CD28 activators, and cultured in XF expansion medium supplemented with IL-2 and IL-21. On day 4 (pre-transduction), the subsets of activated T cells (Fig. 10A, lower panels) were compared to untreated T cells (Fig. 10A, upper panels).

**Fig. 10.**
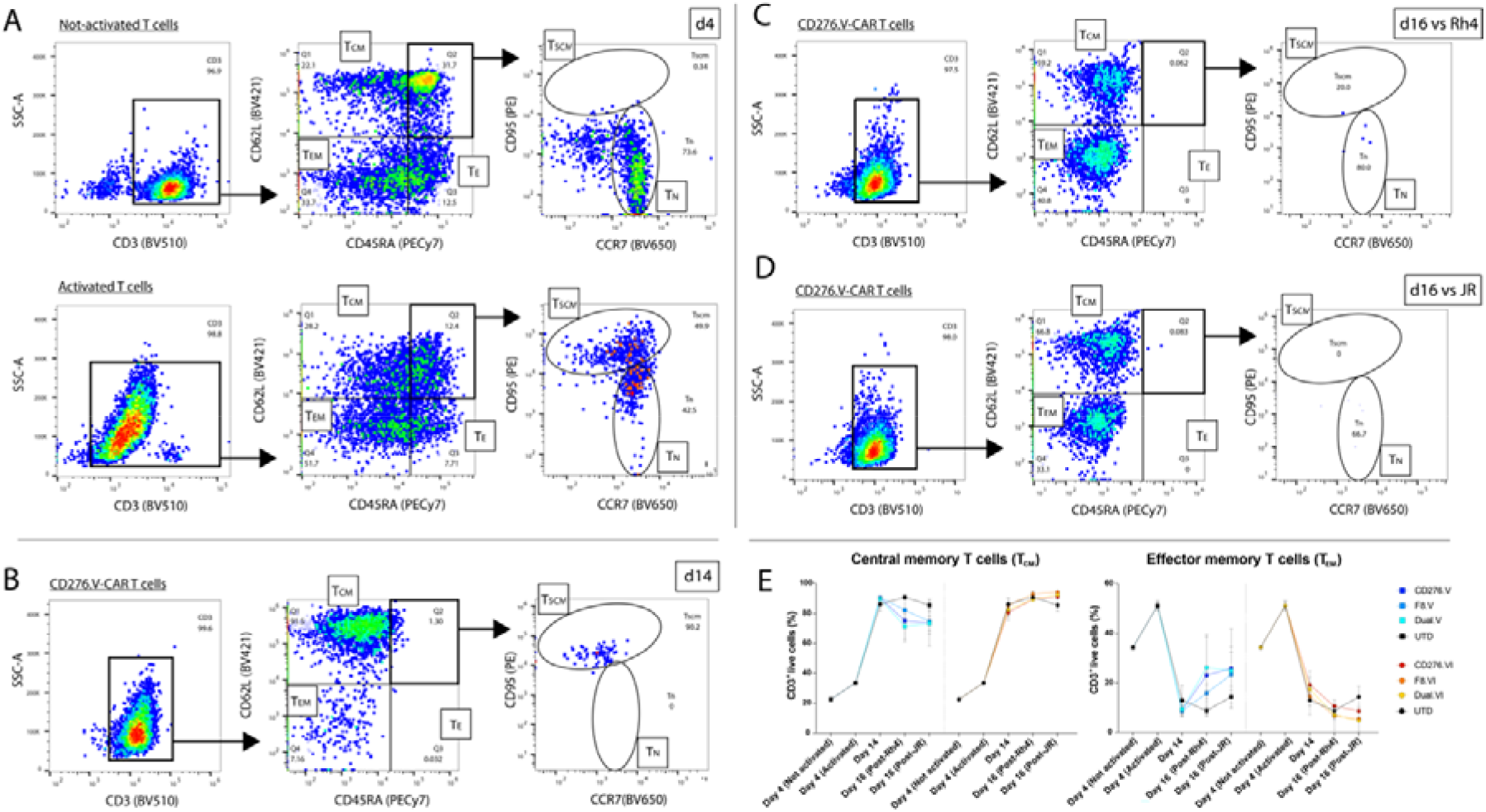
Phenotypic characterization of CAR T cells during expansion and co-incubation experiments. Flow Cytometry during CAR T cells manufacturing was used to quantify percentages of cell memory (T_CM_), effector memory (T_EM_) and effector (T_E_) T cells on CD3^+^ cells by CD45RA and CD62L staining. Naïve (T_N_) and stem cell-like (T_SCM_) T cells were quantified on CD3^+^/CD45RA^+^/CD62L^+^ by CD95 and CCR7 staining. (**A**) On day 4, not-activated (upper panels) and activated (lower panels) T cells were compared. Not-activated T cells showed lower percentages of TCM, TEM, and TE cells than activated T cells (central panels). By contrast, based on CD95 and CCR7 staining, a homogenous population of TN cells could be discriminated in not-activated T cells (right panel), whereas an evident transition towards TSCM could be appreciated in T cells after 4d of activation. Of note, CD3 staining was weaker in activated T cells, due to the presence of CD3/CD28 cross-linking activators. (**B**) On day 14, before co-incubation assays activated CAR T cells showed very high percentage of TCM cells and lower percentages of the other cell populations, compared to day 4. (**C**) On day 16, after co-incubation assay with Rh4 cells, almost 60% of activated CAR T cells had a TCM phenotype, and ca. 40% a TEM phenotype. (**D**) On day 16, after cytotoxicity experiments with JR, almost 70% were TCM cells, whereas ca. 30% were TEM cells.

Of note, we detected a lower CD3 expression in activated T cells compared to not-activated controls (Fig. 10A, left panels), due to the presence of the CD3/CD28 activators competing anti-CD3 staining. The analysis conducted on singlet-gated not-activated CD3^+^ T cells revealed a population of CD62L^+^/CD45RA^+^ naïve T cells (T_N_) of 22.6% ± 0.9% (Fig. 10A, central panels), in contrast to the smaller population (5.0% ± 0.4%) observed in activated T cells. Interestingly, during the first four days of expansion, T_N_ cells showed a transition into CD95^+^/CCR7^-^ stem cell-like memory T cells (T_SCM_) reaching ∼50%, which corresponds to ∼5.9% ± 0.8% of CD3^+^ live cells (Fig. 10A, right panels). No T_SCM_ cells were observed in not-activated T cells. Considering the mean of three donors (Fig. 10E, and Suppl. Table 10), the percentages of CD62L^+^/CD45RA^-^ central memory T cells (T_CM_) were higher in activated T cells (33.5% ± 0.9%) than in not-activated T cells (∼22.6% ± 0.6%). Likewise, CD45RA^-^/CD62L^-^ effector memory T cells (T_EM_) were more abundant in activated T cells (34.4% ± 0.6% vs 51.2% ± 1.7%). To remark, the percentage of effector (T_E_) T cells was lower in activated T cells (3.3% ± 0.5%) compared to not-activated T cells (12.0% ± 0.7%).

Several factors can impact the memory development of CAR T cells. Here we focused on the impact of CD28 and 4-1BB CSDs on memory development of CAR T cells, which have been shown to have a distinct effect on CD19-CAR T cells (74). We performed a direct comparison between the CAR constructs V (28.28.28), containing CD28 CSD, and VI (28.28.BB), containing 4-1BB CSD. After lentiviral transduction on day 4 and sorting on day 7, CD276-, F8-FR4-, and Dual-CAR T cells were expanded until day 14, when they were co-incubated for 48h with Rh4 and JR tumor cells. On day 14, as expected from removal of the CD3/CD28 activators, CD3 levels detected in CAR T cells were comparable to those of untransduced T cells (Fig. 10B, left panel; Suppl. Fig.S6, left panels). The analysis of three different donors showed a uniform CAR T cell differentiation into T_CM_ cells in all the samples (Fig. 10B, central panel; Suppl. Fig.S6, central panels). In fact, both the experimental groups and the respective controls exhibited a consistently high percentage, in the range of 86.5% ± 3.9% T_CM_ cells (Fig. 10E, Suppl. Table S10). A slight difference was instead observed in the number of T_EM_ cells between the constructs of the CD28 group and the 4-1BB group. CD276.V-, F8-FR4.V-, and Dual.V-CAR T cells showed an increase in T_EM_ cells counts compared to CD276.VI-, F8-FR4.VI-, and Dual.VI-CAR T cells (Fig. 10E, Suppl. Fig. S6, S8). In detail, 13.4% ± 9.6% of CD276.V vs 10.7% ± 1.1% of CD276.VI, 16.9% ± 11.2% of F8-FR4.V vs 13.6% ± 1.6% of F8-FR4.VI, and 19.5% ± 7.9% of Dual.V vs 13.9% ± 0.6% Dual.VI, supporting the idea that the backbone V (28.28.28) might induce a differentiation of CAR T cells into T_EM_ after 14 days of culture. No significant differences were noticed in the other T cell subsets within the examined groups, with negligible populations of T_N_/T_SCM_ detected (∼1-2%, Suppl. Fig. S6, S8, Suppl. Table S10).

On day 16, CAR T cells were collected after co-incubation with RMS cells and analyzed. Considering the three donors, CD276-, F8-FR4-, and Dual-CAR T cells showed a consistent profile (Fig. 10C, 10D, Suppl. Fig.S7, Suppl. Table S10). Compared to the untransduced T cells (88.2 ± 4.1%), the majority of detected CAR T cells was classified as T_CM_ (83.3% ± 10.6%), regardless of the type of RMS cell line encountered (Fig. 10E, Suppl. Fig. S7, S8). Percentages in the range of ∼5-25% (15.7% ± 10.8%) were found for T_EM_ cells within all the experimental groups (Fig. 10E, Suppl. Table S10), whereas minimal numbers of T_N_/T_SCM_ and T_E_ were observed. However, the main differences were observed between the constructs V and VI in each condition. In fact, post co-incubation with Rh4, CD276.V-, F8-FR4.V-, and Dual.V-CAR T cells showed a significant increase up to 25% of T_EM_ cells, whereas CD276.VI-, F8-FR4.VI-, and Dual.VI-CAR T cells showed the same values in the three donors, as in the untransduced T cells (6%-10%; Fig. 10E, Suppl. Fig.S7, S8; Suppl. Table S10). This was confirmed also post co-incubation with JR. As shown in Fig.10D and Suppl. Fig. S7, the number of T_EM_ cells was in the range of 24.8% ± 1.2% in CD276.V-, F8-FR4.V-, and Dual.V-CAR T cells. By contrast, T_EM_ cells were 7.1% ± 1.4% in CD276.VI-, F8-FR4.VI-, and Dual.VI-CAR T cells. As observed on day 14, no substantial differences were noticed in the other T cell subsets within the studied groups.

Next, to assess the effect of the manufacturing protocol on CAR T cells exhaustion, and to investigate possible differences between CAR constructs containing CD28 CSD or 4-1BB CSD, the two groups of CAR T cells were analyzed for their expression of CD39, LAG-3, PD-1, and TIM-3 (Fig. 11). CD39 expression was low at day 4 and day 7 of the production, averaging 4% ± 0.2% on day 4 and 6% ± 2% on day 7 (detailed values are reported in Suppl. Table S11), and increased at day 14 ranging 19% ± 4% for the CD28 group and 32% ± 3% for the 4-1BB group. After co-incubation with tumor cells, the CD28 group showed a decreased CD39 expression for F8-FR4.V and Dual.V CAR T cells (16 ± 1% for both), while CD276.V-CARs remained at higher levels of 22% ± 5% when co-incubated with Rh4 and of 21% ± 5% when co-incubated with JR. In the 4-1BB group expression decreased to 25% ± 8% for T cells but remained high at around 33% ± 2%for the CAR constructs. Despite some donor-specific differences, the 4-1BB group surprisingly showed higher expression of CD39 compared to the CD28 group. No significant difference was instead observed between CD276- and F8-FR4-CAR T cells (Fig. 11A).

**Fig. 11.**
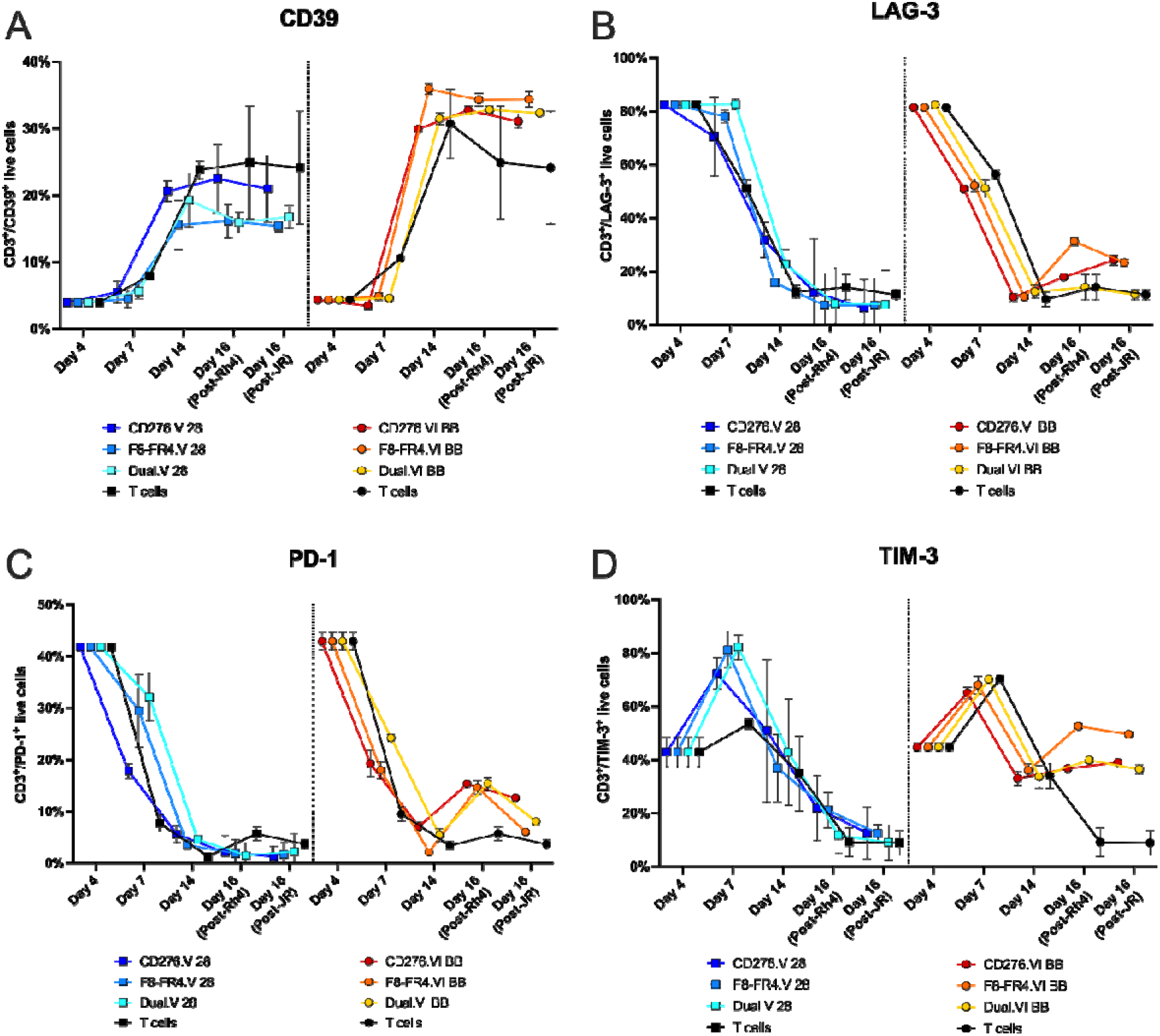
Exhaustion marker expression on CAR T cells during expansion and co-incubation experiments. Expression of the exhaustion markers CD39 (**A**), LAG-3 (**B**), PD-1 (**C**) and TIM-3 (**D**) was measured by FACS in T cells during the manufacturing process and after co-incubation of CAR T cells with RMS cells. T cells were activated on day 1 with IL-2, IL-21 and CD3/CD28 T cells activators. Measurements were performed before lentiviral infection (day 4), after FACS sorting (day 7), at the end of production before co-incubation experiments (day 14), and after two days of co-incubation with RMS cells at E:T ratio of 5:1 (day 16). To compare the impact of CD28 CSD and 4-1BB CSD, two groups of CAR T cells were generated and analyzed: CD276.V-, F8-FR4.V-CAR (with the backbone CD28HD-CD28TM-CD28CDS-CD3ζ) were used to co-transduce T cells and generate Dual.V-CAR T cells; and CD276.VI-, F8-FR4.VI-CAR (with the backbone CD28HD-CD28TM-41BBCDS-CD3ζ) were used to co-transduce T cells and generate Dual.VI-CAR T cells. T cells were treated like CAR T cells, except that they were not transduced on day 4 nor FACS sorted on day 7. Shown are the mean values of three donors with standard deviations.

LAG-3 expression was high in all samples at day 4, 82% ± 1%, as a result of LAG-3 expression induced by IL-2 in activated T cells (75). At day 7, three days after lentiviral transduction, expression in the CD28 group remained high at around 80% (CD276.V: 70% ± 14%; F8.FR4.V: 78% ± 2%; Dual.V: 82% ± 2%), while it decreased in the 4-1BB group to around 50% (CD276.VI: 51% ± 1%; F8.FR4.VI: 52% ± 2%; Dual.VI: 50% ± 3%). At day 14, LAG-3 expression was decreased in both the CD28 groups (average 23% ± 8%) and the 4-1BB groups (average 11% ± 2%), suggesting a recovery of CAR T cells. Upon co-incubation with RMS cells, LAG-3 expression remained stable in the CD28 groups with an average of 8% ± 1%, and was increased in the 4-1BB groups to an average of 20% ± 1% (Fig. 11B).

Mean expression of PD-1 of all the groups was 42% ± 1% at day 4 of CAR T cells production, right before transduction with lentiviruses. At day 14, there was a consistent decrease of PD-1 in all the groups, reaching 4% ± 2%. Upon co-incubation with RMS cells, however, a clear increase was visible in the 4-1BB groups co-incubated with Rh4 cells, where PD-1 expression reached 15% ± 1%, while in the CD28 groups PD-1 expression was 2% ± 2%. In untransduced T cells, PD-1 expression was 5% ± 1%. Activation of T cells clearly induced a transient exhausted phenotype, from which CAR T cells can recover. Surprisingly, co-incubation with RMS cells induced a stronger increase of PD-1 expression in the 4-1BB group than in the CD28 group (Fig. 11C).

In activated T cells, at day 4, expression of TIM-3 was 43% ± 4%, similarly to PD-1. At day 7, after FACS sorting, TIM-3 expression was increased, 70% ± 9%, with the highest levels observed for the CD28 group (79% ± 7%). At day 14, after seven days of expansion in culture, TIM-3 showed an overall decrease, ranging from 33% ± 3% for CD276.VI, 33% ± 4% for Dual.VI, 37% ± 12% for F8-FR4-V and 36% ± 2% forF8-FR4-VI, 51%± 26% for CD276.V. In general, TIM-3 expression was therefore lower in 4-1BB groups compared to CD28 groups at the end of manufacturing. However, upon co-incubation with RMS cells, TIM-3 expression was clearly decreased in the CD28 group, 15% ± 5%, while it slightly increased to 42% ± 1% in the 4-1BB groups (Fig. 11D).To conclude, during the first four days of expansion we observed a differentiation of activated T cells from T_N_ to T_SCM_, corresponding to a very high expression of LAG-3 (>80%) and a strong expression of PD-1 and TIM-3 (40%). On day 14, in each group the majority of CAR T cells uniformly converted into T_CM_ cells, showing a decreased expression of LAG-3, PD-1, and TIM-3, in both CD28 and 4-1BB groups. Measurements performed pre- and post-co-incubation experiments revealed no crucial phenotype alteration within the experimental groups, except for a significant 2-fold increase in the number of T_EM_ cells in the constructs V, compared to the constructs VI. Surprisingly, exhaustion expression of CD39, LAG-3, PD-1, and TIM-3 was increased for all the markers investigated in the 4-1BB after coincubation with RMS cells.

### CD276-CAR T cells eradicate RMS tumors in an orthotopic mouse model

Next, *in vivo* experiments were performed in immunodeficient NSG mice carrying orthotopic RD-derived tumors (CD276^high^FGFR4^low^), with the aim to assess the *in vivo* activity of CD276.V-CAR T cells. RD was selected as a representative FN-RMS cell line presenting high CD276 surface copy number (59’543, Fig.1C). On day 0, four treatment groups were defined: 1) PBS; 2) Untransduced T cells (referred to as UTD); 3) CD19.V-CAR T cells (referred to as CD19); 4) CD276.V-CAR T cells (referred to as CD276). We used PBS, UTD, and CD19 as negative control groups. 0.5×10^6^ fLuc^+^ RD cells were orthotopically injected into the left gastrocnemius muscle of five mice per group, and on day 4, the tumor size was estimated by bioluminescence to prove the uniform tumor growth in all the groups (Fig.12A). On day 5, 5×10^6^ untransduced T cells (UTD), or 5×10^6^ CAR T cells were injected intravenously. Tumor growth was then monitored weekly by bioluminescence (Fig. 12B, 12D). During the third week after RD injection, we observed RMS tumor eradication in 3/5 mice in the CD276 group. By contrast, in the three control groups, RD tumors grew regularly. At week 6, fLuc^+^ RD cells were still undetectable by bioluminescence in 3/5 CD276 mice, indicating a maintained complete remission, whereas in the other two CD276 mice the tumor growth was controlled until week 6, when bioluminescence measurements were stopped for technical reasons. The mice were followed up to day 110, and the tumors measured by caliper, until the tumors in the control group reached the maximal allowed size of 800 mm^3^ (Fig. 12C). At the end of the experiment, 3/5 CD276 mice were still tumor-free. This experiment revealed a promising activity of CD276.V-CAR T cells in promoting clearance of orthotopic RMS tumor. No drop in mouse weight nor evident side effects were observed in the CD276 treated group. Notably, alopecia-like phenotype appeared only in the CD19.V-CAR T cells group during week 5 and persisted until the end of the experiment. All the controls showed a similar tumor growth during the whole experiment. These results revealed a good activity of CD276.V-CAR T cells against CD276^high^ FN-RMS cells, leading to RMS eradication in 3/5 mice and growth reduction in 2/5.

**Fig. 12.**
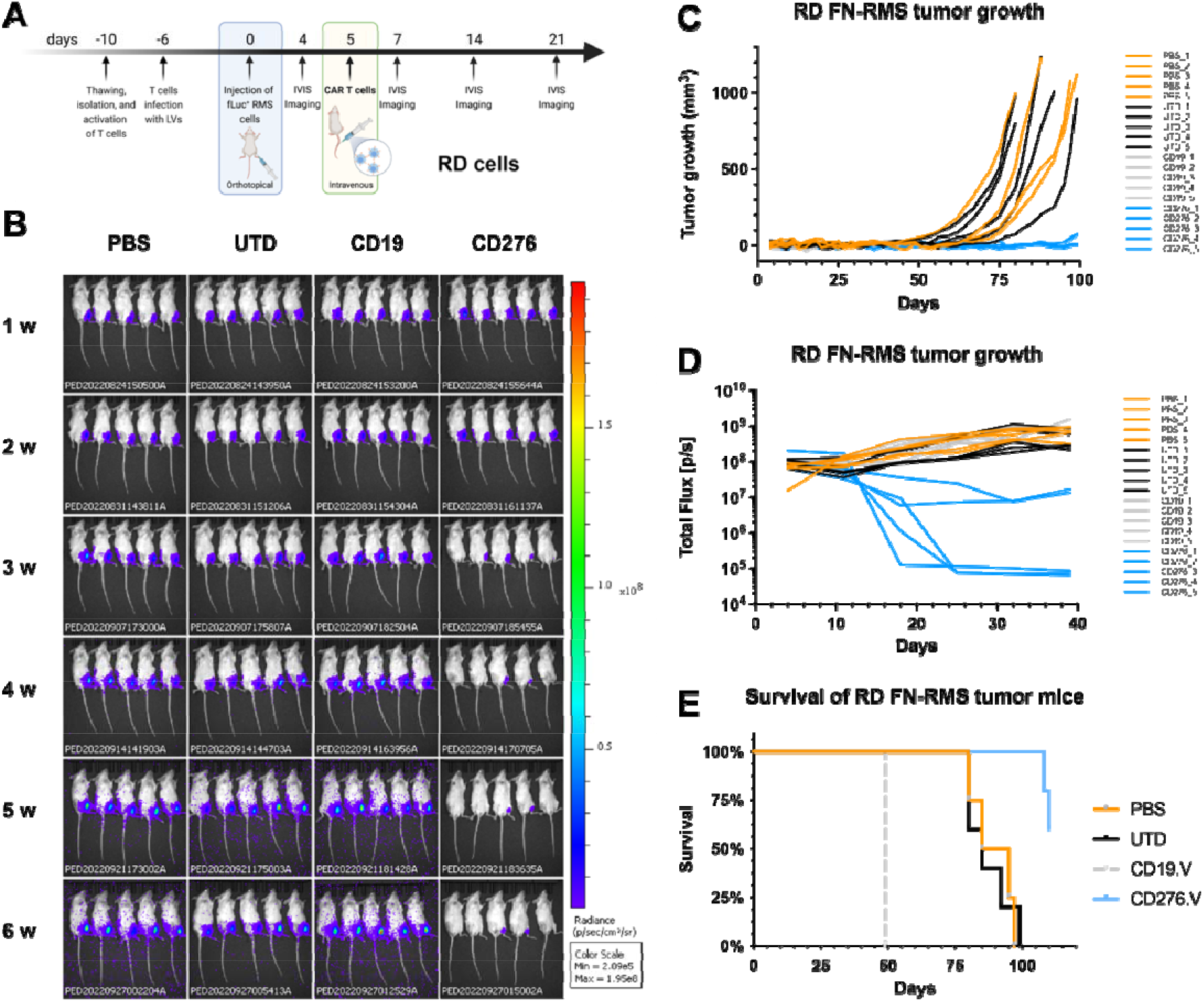
*In vivo* experiment with CD276.V-CAR T cells in mice orthotopically engrafted with RD cells. **(A)** Schematic overview of the experimental design. On day −10, PBMCs were thawed, T cells were purified and activated. On day −6, CAR T cells were produced and expanded until day 5. In the meantime, on day 0, 0.5×10^6^ fLuc^+^ RD cells were orthotopically injected into the left gastrocnemius muscle of mice. After 5 days, CAR T cells were injected into mice by tail vein injection. (**B**) Tumor progression was monitored by bioluminescence photometry. Efficient tumor eradication by CD276.V-CAR T cells was observed in 3/5 mice after three weeks from the injection of RD cells on day 0, whereas CD276.V-CAR T cells showed significant tumor burden control in 2/5 mice. Maintained complete remission was confirmed in 3/5 mice in the group treated with CD276-CAR T cells until the end of the experiment (day 110). Tumor growth curves for the investigated groups was measured by caliper (**C**) and by bioluminescence (**D**). Humane endpoint was set to tumor volume of 800 mm^3^. (**E**) Kaplan-Meier plot showing survival of treated and control mice. On day 50, all the mice from the CD19.V control group had to be removed from the study because of a bacterial infection non-related to the treatment.

### CD276- and CD276/F8-FR4 Dual-CAR T cells induce RMS tumor clearance

Next, we tested in immunodeficient NSG mice the activity of CD276.V-CAR T cells against orthotopic tumors derived from Rh4 cells (CD276^high^FGFR4^high^), expressing high levels of CD276 (50’259, Fig. 1C) and FGFR4 (4’123, Fig. 2C). Here, we also tested F8-FR4.V-CARs against FGFR4, as well as Dual-CAR T cells targeting both CD276 and FGFR4. On day 0, five treatment groups were defined: 1) Untransduced T cells (referred to as UTD); 2) CD19.V-CAR T cells (referred to as CD19); 3) CD276.V-CAR T cells (referred to as CD276); 4) F8-FR4.V-CAR T cells (referred to as F8-FR4); 5) CD276/F8-FR4 Dual-CAR T cells (referred to as Dual). On day 0, 0.5×10^6^ fLuc^+^ Rh4 cells were engrafted into NSG mice and, on day 5, 5×10^6^ untransduced (UTD) T cells, and 5×10^6^ CAR T cells were intravenously injected in the mice (Fig.13A). At week 2, 2/5 mice from the CD276 group and 2/5 from the Dual group showed a significantly smaller tumor burden, compared to the controls (Fig.13B), visible also in the delayed tumor volume growth (Fig. 13C), and stagnation of bioluminescence signal (Fig. 13D). At week 3, all the mice of both CD276 and Dual groups presented a complete remission, which lasted until the end of the experiment (day 59). The F8-FR4 group exhibited a delayed tumor growth in 4/5 mice, compared to the controls until week 6 of the treatment (Fig. 13C, 13D), but failed to completely eradicate the tumors. In the control mice, instead, rapid growth of Rh4-derived xenograft was observed from week 3, so that CD19 mice had to be sacrificed on day 39, due to high tumor burden (>800 mm^3^ tumor size, Fig.13C). For the same reason, after 53 days, all the control and the F8-FR4.V mice had to be sacrificed. At the end of the experiment (day 59), all the mice from CD276.V and Dual-CAR T cells groups were tumor free (Fig. 13D) and did not show any obvious side effect. No drop in mouse weight was observed in the treated groups. These results confirm the excellent activity of CD276.V-CAR T cells against CD276^high^ tumors, leading to complete sustained remission in 5/5 mice, while F8-FR4.V-CAR T cells showed a limited activity, leading to growth delay in 4/5 mice. Interestingly, Dual-CAR T cells showed the same activity as CD276.V-CAR T cells, completely eliminating tumors in 5/5 mice.

**Fig. 13.**
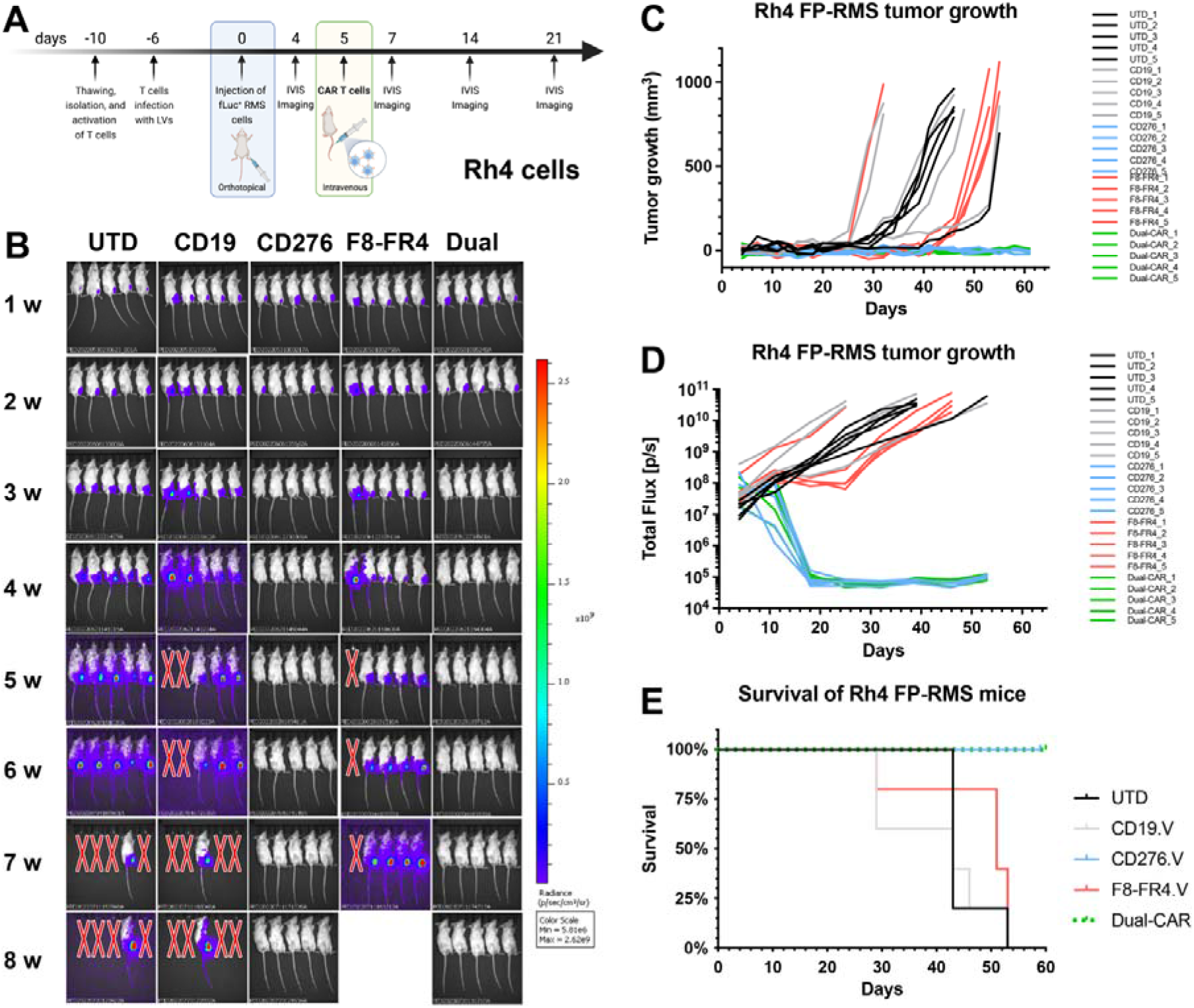
*In vivo* experiment with CD276-CAR T cells, F8-FR4-CAR T cells, and CD276/F8-FR4 Dual-CAR T cells in mice orthotopically engrafted with Rh4 cells. **(A)** Schematic overview of the experimental design. On day −10, PBMCs were thawed, T cells were purified and activated. On day −6, CAR T cells were produced and expanded until day 5. In the meantime, on day 0, 0.5×10^6^ fLuc^+^ Rh4 cells were orthotopically injected into the left gastrocnemius muscle of mice. After 5 days, CAR T cells were injected into mice by tail vein injection. (**B**) Tumor progression was monitored by bioluminescence photometry. Efficient tumor eradication by CD276.V-CAR T cells and Dual-CAR T cells was observed in 5/5 mice 3 weeks after Rh4 cells injection on day 0. Maintained complete remission was confirmed in both groups until the end of the experiment (day 60). Tumor growth curves for the investigated groups was measured by caliper (**C**) and by bioluminescence (**D**). (**E**) Kaplan-Meier plot showing survival of treated and control mice. Humane endpoint was set to tumor volume reaching 800 mm^3^.

### Limited eradication in CD276^low^ RMS tumors

In order to test CD276-CAR T cells activity *in vivo* against tumor cells expressing lower levels of CD276, and to investigate the effect of combining CD276/F8-FR4.V Dual-CAR T cells, the same experimental groups of CAR T cells described above (Fig. 13) were investigated in JR cells (CD276^low^FGFR4^high^) expressing low levels of CD276 (7’964, Fig. 1C) and medium-high levels of FGFR4 (3’133, Fig. 2C). On day 0, six treatment groups were defined: 1) Untreated (referred to as PBS); 2) Untransduced T cells (referred to as UTD); 3) CD19.V-CAR T cells (referred to as CD19); 4) CD276.V-CAR T cells (referred to as CD276); 5) F8-FR4.V-CAR T cells (referred to as F8-FR4); 6) CD276/F8-FR4 Dual-CAR T cells (referred to as Dual) (Fig.14A). We used CD19 as negative control group. On day 0, 0.5×10^6^ fLuc^+^ JR cells were engrafted into NSG mice and, on day 5, 5×10^6^ untransduced (UTD) T cells, and 5×10^6^ CAR T cells, were intravenously injected into the mice. During week 2, bioluminescence imaging revealed a reduction in tumor size in 1/5 mice from the CD276 group and in 1/5 mice from the F8-FR4 group, compared to the controls (Fig. 14B, 14D). However, from week 3, only one mouse (nr. 2) from the CD276 group exhibited a complete remission, which lasted until the end of the experiment (Fig.14B-D). Unfortunately, none of the mice from the other experimental groups showed tumor eradication, nor a significantly delay in tumor growth. Instead, all the mice showed a rapid and homogenous tumor increase (Fig. 14C), and the mice were euthanized during week 6, having reached the termination criteria. We did not observe any drop in mouse weight, except for one mouse (nr. 3) of the PBS group, which was sacrificed once it lost 15% of the initial weight. Mice of the CD19 group exhibited fur loss on the back of the neck, but no further side effects were observed. These results revealed the limit of CD276.V-CAR T cells against tumors expressing lower levels of CD276 and confirmed the limited activity of F8-FR4-CAR T cells.

**Fig. 14.**
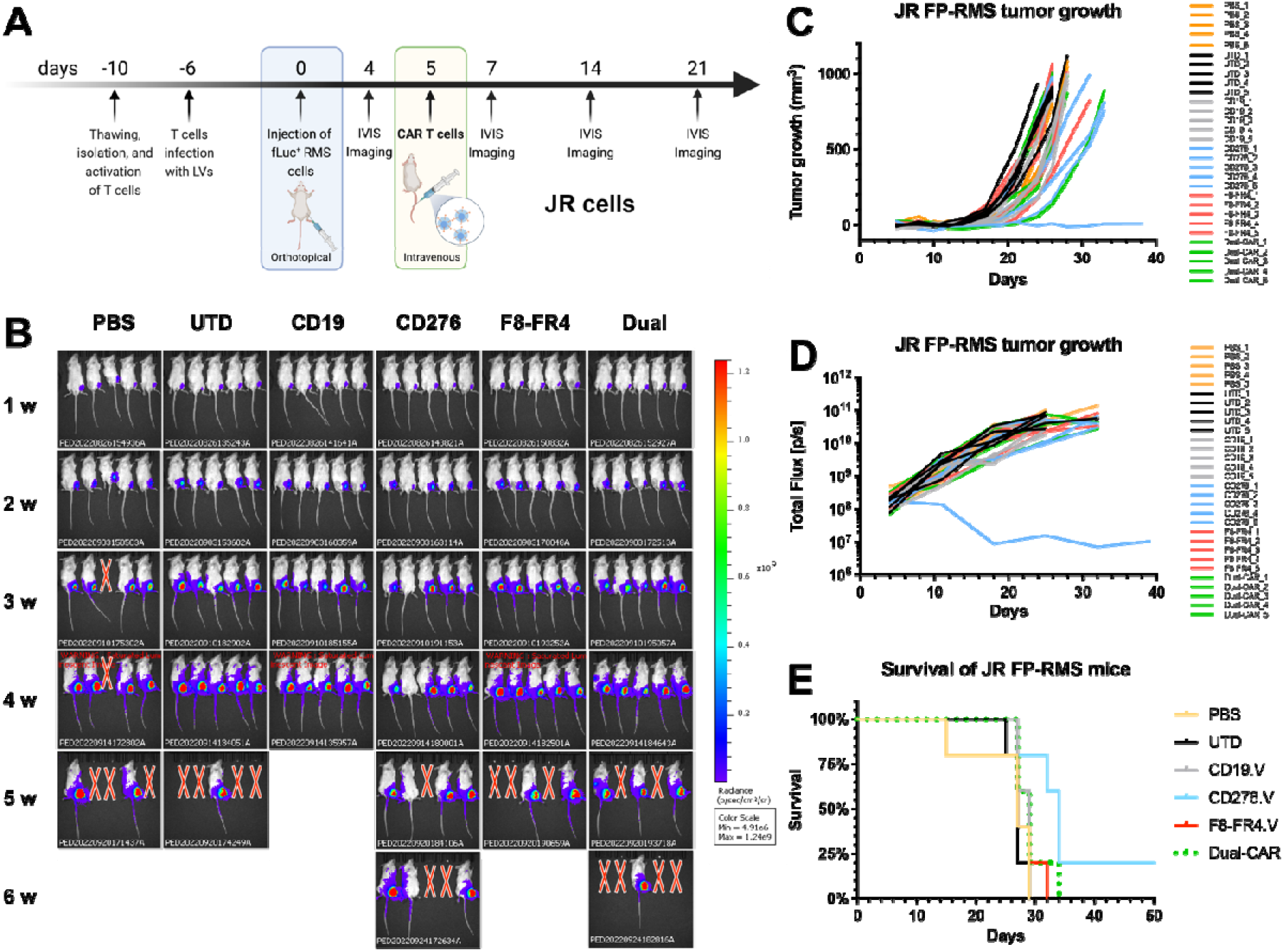
*In vivo* experiment with CD276-CAR T cells, F8-FR4-CAR T cells, and CD276/F8-FR4 Dual-CAR T cells in mice orthotopically engrafted with JR cells. **(A)** Schematic overview of the experimental design. On day −10, PBMCs were thawed, T cells were purified, and activated. On day −6, CAR T cells were produced and expanded until day 5. In the meantime, on day 0, 0.5×10^6^ fLuc^+^ JR cells were orthotopically engrafted into the left gastrocnemius muscle of mice. After 5 days, CAR T cells were injected into mice by tail vein injection. (**B**) Tumor progression monitored by bioluminescence photometry. Tumor eradication was observed in 1/5 mouse in the CD276.V-CAR T cells treated group, which was maintained until the end of the experiment on day 50. Tumor growth curves for the investigated groups was measured by caliper (**C**) and by bioluminescence (**D**). CD276.V-CAR T cells and Dual-CAR T cells delayed tumor growth in 3/5 and 1/5 mice, respectively. (**E**) Kaplan-Meier plot showing survival of treated and control mice. Humane endpoint was set to tumor reaching 800 mm^3^.

### Decreased expression of CD276 and FGFR4 in tumors compared to cultured cells

To investigate if expression of the CAR targets in orthotopic tumors corresponded to what observed on cultured cell lines, CD276 and FGFR4 expression was assessed by Flow Cytometry on single cells extracted from Rh4 tumors (Fig. 15). Flow cytometry results showed a lower CD276 expression on tumor xenografts-derived Rh4 cells, with a fluorescence intensity 14-fold higher than the control, compared to cultured Rh4 cells (600-fold over control) (Fig. 15A). Interestingly, a much lower expression of FGFR4 was observed in tumor xenografts-derived Rh4 cells (2-fold over isotype control) compared to cultured wild-type Rh4 cells (97-fold over control) (Fig. 15B). An estimation of the magnitude of the decrease of CD276 and FGFR4 based on the fluorescence intensity (geometric mean), indicates a similar decrease in tumors, 40-fold (CD276) and 50-fold (FGFR4), compared to cultured cells.

**Fig. 15.**
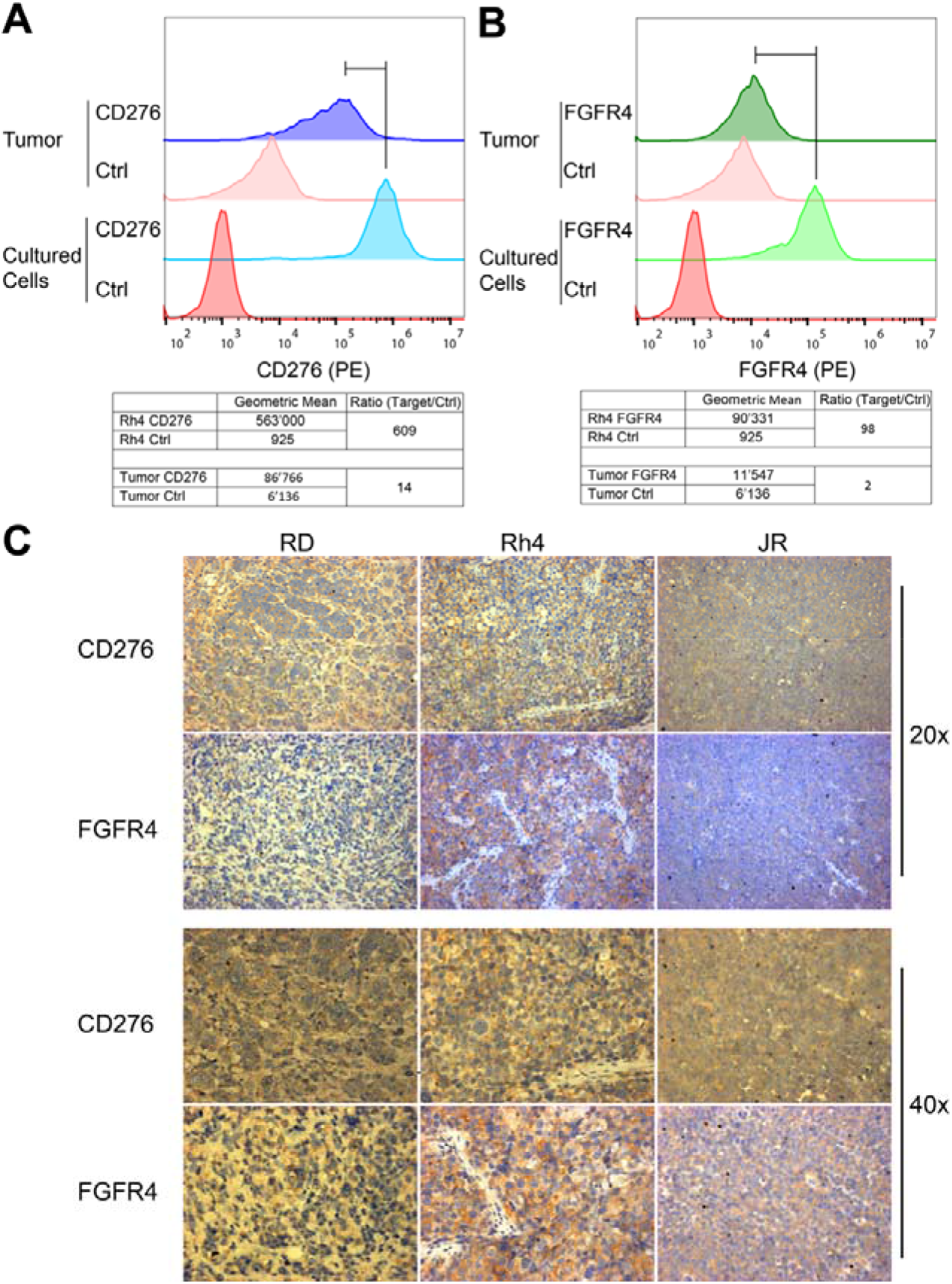
Expression of CD276 and FGFR4 on RMS-derived xenografts. **(A)** Expression of CD276 was assessed by Flow Cytometry on untreated Rh4-derived xenograft tumors and compared to cultured wild-type Rh4 cells by using a PE-conjugated anti-CD276 antibody. CD276 expression was calculated dividing the geometric mean of antibody-stained cells by the geometric mean of the controls. CD276 expression showed a lower CD276 expression on Rh4 tumor-derived cells, compared to the cultured Rh4 cells. (**B**) Expression of FGFR4 was assessed by Flow Cytometry on Rh4-derived xenograft tumors from untreated mice and compared to cultured wild-type Rh4 cells by using a PE-conjugated anti-FGFR4 antibody. FGFR4 expression was calculated dividing the geometric mean of antibody-stained cells by the geometric mean of the isotype controls. The results indicated a strong decrease in FGFR4 expression in Rh4 tumor xenograft-derived cells. (**C**) Assessment of CD276 and FGFR4 expression in tumor tissue xenografts by IHC. After treatment, CD276 is still highly detectable on RD- and Rh4-derived tumor xenografts, but low staining detection was observed in JR-derived tumors. FGFR4 seems expressed at low levels on RD, at medium levels on JR, and at high levels on Rh4.

To confirm these results, IHC staining of CD276 and FGFR4 was performed on formalin-fixed and paraffin-embedded sections of RD-, Rh4-, and JR-derived tumor xenografts from control mice (Fig. 15C). IHC staining results on tumors confirmed uniform and higher expression of CD276 in RD and Rh4 derived tumors, compared to JR derived tumors (Fig. 15C). FGFR4 was detected at high levels in Rh4-, at medium levels in JR-, and at low levels in RD-derived tumors (Fig.15C). Taken together, these results indicate that both targets are clearly detectable in Rh4 mouse tumor xenografts, but they are expressed at different levels in RD and JR. In conclusion, CAR targets seem to be expressed in tumors at lower levels compared to cultured cell lines, but their relative expression seems to be maintained.

### CAR T cells persistence in mouse spleen

CAR T cells persistence *in vivo* is an important aspect that determines the therapeutic impact. To investigate this, we performed CD3 staining on JR-derived tumors collected from CD276.V-CAR T cells-treated mice at the end of the experiment, after 5 weeks from tumor cells implantation, and from Rh4-derived tumors collected from F8-FR4.V-CAR T cells-treated mice at the end of the experiment, after 7 weeks from tumor cells implantation. IHC CD3 staining revealed the presence of CAR T cells in both tumors (Fig. 16A). In JR tumors CAR T cells were visible only in structures that appear to be vessels (Fig. 16A, black arrowheads) and CAR T cells seemed not to have penetrated the tumor tissue, whereas in Rh4 tumors CAR T cells appeared to have infiltrated in the tumor. Since Rh4-derived tumors were cleared by CD276.V- and Dual-CAR T cells, we investigated if persistent CAR T cells could be still detected in the spleen of mice treated with CD276.V-CAR T cells. IHC analyses detected T cells in the spleen of Rh4-bearing mice injected with control untransduced T cells (UTD control) and in the spleen of CD276.V-CAR T cells treated mice, suggesting persistence of CAR T cells even after tumor eradication (Fig. 16B). IHC analyses on normal tissues from control mice and from Rh4-tumor bearing mice were performed to evaluate any sign of toxicity caused by CAR T cells. Hematoxylin and Eosin staining in liver, muscle, brain, heart, and spleen of untreated and treated mice showed similar phenotype (Suppl. Fig. S9). No evident necrosis nor inflammation were observed in the tissues analyzed, suggesting that CD276.V- and Dual-CAR T cells specifically recognize the targets of interest without damaging the investigated normal mouse tissues.

**Fig. 16.**
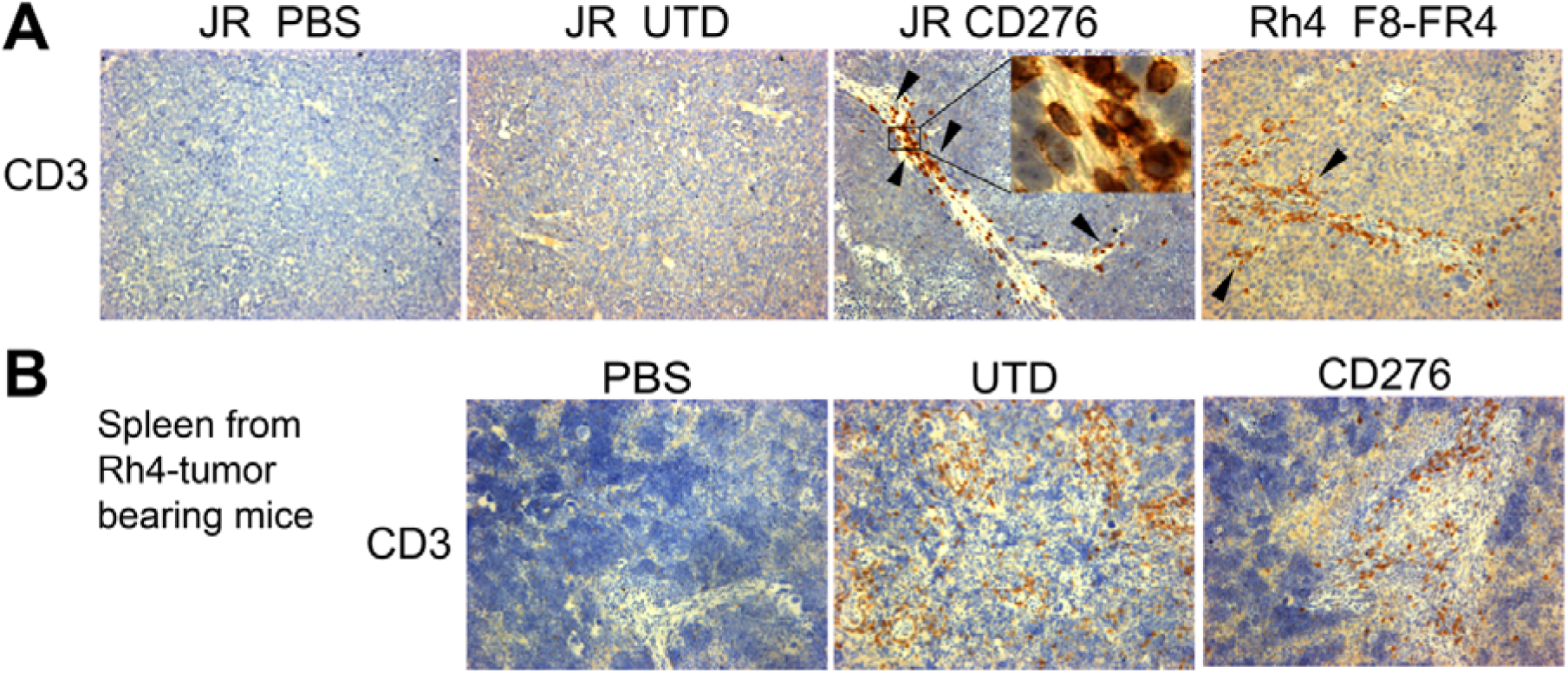
T cell detection in tumors and spleen. **(A)** Detection of T cells in orthotopic tumors derived from JR or Rh4 cells. Tumors were collected at the end of the experiment, after 5 weeks from JR implantation, and after 7 weeks from Rh4 implantation. CD3 staining revealed presence of CAR T cells in JR- and Rh4-derived xenografts. However, in JR xenografts CAR T cells were only detectable in the vessels (black arrows and inset), whereas T cells seemed to have infiltrated Rh4 tumors. (**B**) Evidence of CAR T cells persistence in the spleen CD3 staining revealed presence of T cells in the spleen of mice bearing Rh4 orthotopic tumors and injected with untransduced T cells (UTD) and CD276.V-CAR T cells at the end of the experiment, after 8 weeks and 9 weeks from Rh4 implantation, respectively.

In conclusion, these results indicate that even after unsuccessful treatment, both CD276.V- and F8-FR4-CAR T cells can be detected in JR tumors and suggest that CD276-CAR T cells might be limited in reaching tumors cells. In Rh4-tumor mice, where tumors could be eradicated, CAR T cells could be detected even several weeks after tumor clearance, indicating a good persistence potential.

## Discussion

In this study, we investigated the impact of specific domains included in the CAR constructs and adopted a pragmatic approach, by *in vitro* screening of a combinatorial pool of eight CD276-directed CAR constructs, with CD8α or CD28 HD/TM domains and one or two intracellular CSDs from 4-1BB and/or CD28. CD276.V-CAR T cells (276.MG.28HD/TM.28CSD.3z), were the most effective in killing FN-RMS RD and FP-RMS Rh4 cells *in vitro* and released the highest levels of IFN-γ and Granzyme B. CD276.V-CAR T cells were able to eradicate CD276^high^ Rh4 tumors *in vivo* and persisted even after tumor elimination.

These results confirm recent studies demonstrating that CAR T cells with CD28 HD/TM domains show more reactivity to low-antigen density tumors, faster killing, and higher levels of IL-2 release, when compared to CARs with CD8α HD/TM domains (51). Superior anti-tumor efficacy of CD28 HD/TM was also reported in HER2-positive breast cancer and CD276-positive neuroblastoma (51). On the other side, CD8α HD/TM domains are associated with a lower production of cytokines, such as IL-2, IFN-γ, and TNF-α, when compared to CD28 HD/TM domains. However, despite these reduced levels, CD8α-mediated cytokine-release syndrome (CRS) was still observed in patients (76). In this context, we chose the killing capacity of CAR T cells and the release of cytokines, such as IFN-γ, as predictive readouts, due to the important role played by IFN-γ in targeting solid tumors (77).

From investigations of different co-stimulatory domains (CSDs) in mouse models, we know that CD28 CSD is associated with a higher release of inflammatory cytokines and with an exhausted phenotype (56). Moreover, an augmented CRS occurrence was observed in patients treated with CD19-CD28 CAR T cells, compared to CD19-4-1BB CAR T, highlighting a more cytotoxic effect by CD28 CSD (78, 79). Our results are in good agreement with a stronger activity conferred by CD28 CSD.

Our results on successful use of CD276.V-CAR T in CD276^high^ RMS tumors are in line with the results obtained with other CAR constructs targeting CD276 in preclinical studies against several types of tumors: 276.8αHD/TM.28CSD.BBCSD.3ζ against glioblastoma (80); 276.MG.8αHD/TM.BBCSD.3ζ against medulloblastoma (70), neuroblastoma (70, 81), osteosarcoma (70), and Ewing sarcoma (70); 276.8H9S3.3.8α.28TM.BBCSD.3ζ against gastric cancer (82); and 276.8αHD.28TM.28CSD.3ζ and 276.8αHD/TM.BBCSD.3ζ against esophagus squamous cell carcinoma (83). All these CAR constructs generally allowed T cells to increase cytokine release *in vitro*, to prolong survival, and to persist *in vivo*.

In our screening, second-generation CAR T cells with one CSD, outperformed third-generation CAR T cells with two CSDs. This result was surprising, since several preclinical studies with third-generation CD19-CAR T cells hinted at a higher intracellular signaling activity compared to the second-generation CAR constructs (84). Co-expression of both CD28 and 4-1BB CSDs was reported to compensate the possible limitations of single CSD, improving expansion and showing superior persistence (85). For these reasons, we expected that third-generation CAR T cells would show a more effective killing capacity and a higher cytokine release. On the other side, some studies revealed lower surface expression of third-generation CARs and inferior persistence, compared to second-generation (86).

In this work, we established a 14-day long manufacturing protocol able to isolate from PBMCs a main population of T_N_ cells (∼20%) and to promote their transition into T_SCM_ within day 7. On day 14, we observed an enrichment for a large population of CAR T cells (>70%) reflecting the phenotype of central memory T cells (T_CM_). This profile was observed in all the tested conditions and groups, revealing no inter-group nor intra-group phenotypical difference. In the context of CAR T cells, using T_N/SCM_-derived T_CM_ CAR T cells might be clinically beneficial, since T_CM_ cells represent a less differentiated grade of memory T cells and exhibit a potent anti-tumor ability (87–89). Also, it was demonstrated that T_CM_ CAR T cells can persist longer *in vivo* than T_N_ or T_EM_ cells (90) and present a good safety profile (91, 92). Recent studies have shown that T_SCM_ play a central role in the early anti-leukemia response and late immune surveillance (93), suggesting that producing CAR T cells deriving from T_SCM_ subsets may represent a potential therapeutic advantage. However, analyses conducted post-co-incubation with RMS cells, on day 16, exhibited a differentiation of ∼5-10% T_CM_ into T_EM_ CAR T cells exclusively for the groups CD276.V- and Dual.V-CAR T cells, which expressed CD28 as CSD and showed high killing efficacy *in vitro*. Interestingly, this change was not observed in CAR T cells with 4-1BB as CSD that eliminated RMS cells with lower efficacy, suggesting a link between the high cytotoxicity observed for the groups CD276.V-, F8-FR4.V-, and Dual.V-CAR T cells, and their reduced memory phenotype.

Exhaustion analysis of untransduced T cells and CD276-, F8-FR4-, and Dual-CAR T cells revealed variable expression of PD-1, LAG-3, CD39, and TIM-3 markers during the manufacturing protocol and *in vitro* tests. As extensively reported in the literature, PD-1 is considered as an early T cell exhaustion marker. Our results showed transient and moderate expression levels (∼40%) of PD-1 on day 4, which drastically decreased along T cell expansion, reaching negligible levels on days 14. Before co-incubation with RMS cells, no significant difference was observed between the experimental CAR T cell groups and the control. However, after two days co-incubation with RMS cells, higher levels of PD-1 (∼15%) were detected in the 4-1BB CSD group CD276.VI-, F8-FR4.VI-, and Dual.VI-CAR T cells, compared to the CD28 CSD group CD276.V-, F8-FR4.V-, and Dual.V-CAR T cells (<5%). LAG-3 is an intermediate exhaustion marker. LAG-3 was detected at very high levels (∼80%) in all the investigated samples on days 4 and 7, but similarly to PD-1, its expression reached less than 10% on day 14. Interestingly, after co-incubation with RMS cells, on day 16, the 4-1BB CSD group displayed >2.5-fold LAG-3 expression than the CD28 CSD group (∼20% vs ∼8%). CD39 and TIM-3 are considered terminal exhaustion markers and exhibited an opposite trend compared to PD-1 and LAG-3. After T cell activation, CD39 showed very low levels (∼5%) in all the experimental groups, and no significant difference was observed with the untransduced T cells. On day 14, CD39 levels increased up to ∼15-20% for the CD28 CSD group, and ∼35% for the 4-1BB CSD group, maintaining a comparable expression even after co-incubation with RMS cells. Likewise, TIM-3 exhibited a moderate expression (∼40%) on day 4, reached a peak on day 7 (∼80%), and constantly decreased to ∼10% in the CD28 CSD group. On the other hand, the 4-1BB CSD group followed this behavior only until d14, when the levels of TIM-3 were at ∼40%. However, after co-incubation with Rh4 and JR cells, TIM-3 levels remained moderate at ∼40%. To conclude, the main differences we observed were between the CAR T cells expressing the backbone 28.28.3z and those expressing 28.28.BB, with the latter ones displaying higher exhaustion at each time-point.

These results indicate that IL-2 and CD3/CD28 activators induce an initial stress on T cells. However, CAR T cells were capable to recover and properly expand during manufacturing until d14, after which they engaged with high efficacy RMS cells. To note, in contrast with what is described in the literature, our CD28 CSD group CD276.V-, F8-FR4.V-, and Dual.V-CAR T cells outperformed their 4-1BB CSD counterparts CD276.VI-, F8-FR4.VI-, and Dual.VI-, respectively, showing high killing capacity *in vitro* and a less exhausted profile pre- and post-co-incubation.

*In vivo*, CD276.V-CAR T cells successfully eradicated Rh4-derived tumors at week 2 of treatment with 100% efficiency whereby in 5/5 mice the tumor did not relapse. The efficacy against RD-derived tumors was of around 60%, with 3/5 mice showing a clear maintained complete remission. Notably, the other two mice showed an initial significant treatment response, but not sufficient to achieve a complete remission. Since Rh4 and RD cells lines show similar CD276^high^ surface density *in vitro*, a plausible explanation of the different efficacy might be found in the different growth rate: slow growing FN-RMS RD possibly impede CD276-CAR T cells activity due to the small size of the RD-derived tumors that might not provide sufficient stimulation to activate CAR T cells and to allow their expansion and persistence. In fact, RD tumors could only be monitored by bioluminescence, and not measured by caliper. This might theoretically also reduce homing and decrease the probability by CD276-CAR T cells to encounter the target, protecting RD tumor cells during the first week of experiment. Future experiments with either implantation of more RD cells, or with CAR T cells injection at a later time point with a larger RD-tumor size, might answer this question. CD276.V-CAR T cells failed to control growth ofCD276^low^ JR-derived tumors.

CD276.V-CAR T cells were detectable in tumors after 5 weeks, indicating that a lack of persistence might not be the reason for the limited activity. This lack of activity was disappointing, since CD276.V killing of JR cells was comparable to Rh4 and RD cells *in vitro*, but not surprising, since the dependence of CARs on antigen density for effective activity has already been recognized (51, 70, 94). On the same line, F8-FR4-CAR T cells failed to eradicate Rh4-derived tumors and the strongest effect observed was just a 1-week delay in tumor expansion. F8-FR4-CAR T cells target FGFR4 which is expressed at lower levels than CD276 on Rh4 cells, with C 3’000 FGFR4 copies per cell vs C 50’000 of CD276. Therefore, a possible explanation for the lack of tumor eradication might simply be an insufficient activation of F8-FR4-CAR T cells. However, recent work published by Sullivan et al. (47) showed that a combination of FGFR4-CAR T cells with pharmacologic inhibition of the myeloid component in stroma in Rh30 RMS-derived tumors allowed FGFR4-CAR T cells to eradicate orthotopic RMS in mice (47). It is interesting to note that Rh30 cells were reported to express 2’700 FGFR4 molecules per cell, while we measured 2’000 molecules/cell in the Rh30 cell line in our laboratory and 3’700 molecules per cell in Rh4 cells. These results suggest that it is possible to overcome the limitations posed by a low-density target by manipulating the myeloid compartment in RMS. Alternatively, expression of c-Jun gene in F8-FR4.V-CAR T cells, which have the same CAR backbone used to target low GD2 expressing cells, could lead to *in vivo* eradication of tumors with low antigen density without side effects (95).

To increase CAR T cells activity, and to overcome a possible antigen loss, we investigated a Dual-CAR T cell strategy by targeting CD276 and FGFR4 at the same time. Dual-CAR T cells co-expressing on their surface equivalent numbers of CD276.V-CARs and F8-FR4.V CARs were tested *in vitro* against Rh4 cells showing a slightly better killing capacity compared to CD276.V-CAR T cells, while a lower cytotoxicity was observed for F8-FR4.V CAR T cells. In Rh4-tumor bearing mice, CD276/FGFR4 Dual-CAR T cells showed a similar complete eradication as CD276.V-CAR T cells. This is an important observation in the optic of combining CD276 with other CARs, since it implies that they are still fully active even if the number of CD276 CARs on the surface of T cells is reduced. Unfortunately, Dual-CAR T cells failed to show an improved effect, and were not effective against CD276^low^FGFR4^high^ JR cells, just like F8-FR4.V- and CD276.V-CAR T, indicating a direct correlation between target surface density and efficacy of CAR T cells. In fact, despite the high efficacy shown with RD and Rh4, CD276.V-CAR T cells exhibited reduced killing capacity against JR.

Currently, there are three ongoing CD276 clinical trials: NCT04691713 is recruiting patients with advanced CD276-positive solid tumors; NCT04432649 is a Phase 2 study investigating the activity of 4SCAR-276 CAR T cells in brain tumors and Ewing sarcoma; NCT04185038 is a Phase I clinical trial for Diffuse Intrinsic Pontine Glioma/Diffuse Midline Glioma and recurrent or refractory pediatric central nervous system tumors. Our results support the inclusion of patients with CD276-positive RMS in current clinical trials.

## Conclusions

CD276.V-CAR T cells containing CD28-derived HD/TM and CSD are able to control and eradicate orthotopic RMS tumors, when their target is expressed above a critical threshold. FGFR4-targeted CAR T cells can only partially control RMS growth, possibly because of low target expression and immunosuppressive environment. CD276/FGFR4 Dual-CAR T cells showed the same activity as CD276-CAR T cells, suggesting that dilution of CD276-CAR does not detrimentally impact their activity. Targeting multiple antigens, increasing signal strength through optimized CAR design, remedying the limitations imposed by the immunosuppressive TME will be necessary to achieve consistently complete remission.

### List of abbreviations

ABD: antigen binding domain; AICD: activation-induced cell death; APC: antigen presenting cells; aRMS: alveolar RMS; CAR: chimeric antigen receptor; CSD: co-stimulatory domain; EFS: elongation factor 1α short promotor; eRMS: embryonal RMS; FN-RMS: fusion negative RMS; FP-RMS: fusion positive RMS; HD: hinge domain; IHC: immunohistochemistry; LTR: long terminal repeat; scFv: single chain variable fragment; PDX: patient derived xenograft; RMS: rhabdomyosarcoma; sdAb: single-domain antibodies; TM: transmembrane domain; TME: tumor microenvironment; UTD: untreated; WB: Western blotting.

## Declarations

### Ethics approval and consent to participate

The mice experiments were approved by Animal Welfare Office of Canton Bern and in accordance with Swiss Federal Law (Authorization BE22/2022).

### Consent for publication

Not applicable

### Availability of data and materials

The datasets used and/or analyzed during the current study are available from the corresponding author on reasonable request.

### Authors’ contributions

AT: Design and execution of experiments, supervision, analysis, interpretation of data, writing, and funding acquisition. CP: Design and execution of experiments, analysis and interpretation of data, critical reading of the manuscript. DD: Execution of experiments, analysis and interpretation of data. SAJ: Execution of experiments, analysis and interpretation of data. AR: Execution of experiments, analysis and interpretation of data. JM: Execution of experiments, analysis and interpretation of data. JR: Supervision, funding acquisition, and critical reading of the manuscript. MB: Design of the study, interpretation of data, supervision, writing, and funding acquisition. All authors read and approved the final manuscript.

## Acknowledgements

We thank the Institute of Myology, Paris France, for providing humanized myoblasts. We are also very grateful to Dr. Stephan Müller and the Flow Cytometry and Cell Sorting (FCCS) facility of the DBMR (University of Bern) for their support. We are grateful to Prof. Andreina Schöberlein and her lab for their help with immunohistochemistry.

**Authors’ information (optional)**

