## Supplementary Material for "CAR T cells recognizing CD276 and Dual-CAR T cells against CD276/FGFR4 promote rhabdomyosarcoma clearance in orthotopic mouse models"

**Additional Files**

**Supplementary Tables:**

### Supplementary Table S1. Primers used in this study.

| Primer name | Sequence |
| --- | --- |
| P2A-tCD19 Forward | CGGCGCCACCAACTTCAGCCTGCTGAAGCAGGCCGGCGACGTGGAGGAGAACCCCGGCCCCCCACCTCCTCGCCTCCTC |
| P2A-tCD19 Reverse | TTCACAAATTTTGTAATCCAGAGGTTGATTGTCGACTTAATCACAGGACCAGGGCTCTT |

### Supplementary Table S2. sgRNA sequences used to target CD276 and FGFR4.

| Direction | Target | Guide Sequence (5' -> 3') | PAM | Exon |
| --- | --- | --- | --- | --- |
| + | CD276 | GCTGGTGCACAGCTTTGCTG | AGG | 8 |
| + | FGFR4 | TGGTGGCCACTGGTACAAGG | AGG | 6 |

### Supplementary Table S3. List of the CAR plasmids used for the production of CAR T cells.

| Plasmid | CAR | Addgene # |
| --- | --- | --- |
| MB0106 | CD19.8h.8TM.28.3z | 200670 |
| MB0107 | CD19.8h.8TM.BB.3z | 200671 |
| MB0108 | CD19.8h.8TM.28.BB.3z | 200672 |
| MB0109 | CD19.8h.8TM.BB.28.3z | 200673 |
| MB0110* | CD19.8h.28TM.28.3z | 200674 |
| MB0111* | CD19.8h.28TM.BB.3z | 200675 |
| MB0112* | CD19.8h.28TM.28.BB.3z | 200676 |
| MB0113* | CD19.8h.28TM.BB.28.3z | 200677 |
| MB0114 | CD19.28h.28TM.28.3z | 200678 |
| MB0115 | CD19.28h.28TM.BB.3z | 200679 |
| MB0116 | CD19.28h.28TM.28.BB.3z | 200680 |
| MB0117 | CD19.28h.28TM.BB.28.3z | 200681 |
| MB0118 | CD276.8h.8TM.28.3z |  |
| MB0119 | CD276.8h.8TM.BB.3z |  |
| MB0120 | CD276.8h.8TM.28.BB.3z |  |
| MB0121 | CD276.8h.8TM.BB.28.3z |  |
| MB0126 | CD276.28h.28TM.28.3z |  |
| MB0127 | CD276.28h.28TM.BB.3z |  |
| MB0128 | CD276.28h.28TM.28.BB.3z |  |
| MB0129 | CD276.28h.28TM.BB.28.3z |  |
| MB0130 | F8-FR4.8h.8TM.28.3z |  |
| MB0131 | F8-FR4.8h.8TM.BB.3z |  |
| MB0132 | F8-FR4.8h.8TM.28.BB.3z |  |
| MB0133 | F8-FR4.8h.8TM.BB.28.3z |  |
| MB0134 | F8-FR4.28h.28TM.28.3z |  |
| MB0135 | F8-FR4.28h.28TM.BB.3z |  |
| MB0136 | F8-FR4.28h.28TM.28.BB.3z |  |
| MB0137 | F8-FR4.28h.28TM.BB.28.3z |  |

*not used in this study

### Supplementary Table S4. Statistical analysis of sdAb-FR4-CAR T cells killing capacity.

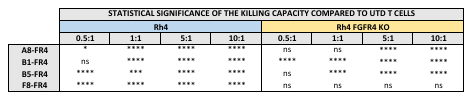

The statistical significance of the differences between experimental and control groups was assessed using Dunnett’s multiple comparison test following a two-way ANOVA (p>0.05 (ns), p ≤ 0.05 (*), p ≤ 0.01 (**), p ≤ 0.001 (***), p ≤ 0.0001 (****)).

### Supplementary Table S5. T cells infection efficiency assessed at day 6 by GFP and CAR expression assessed by MycTag expression.

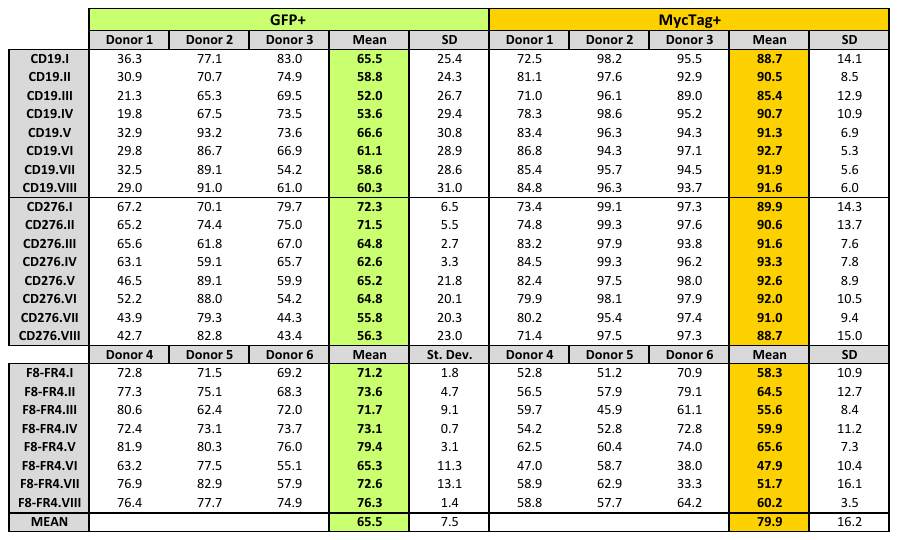

### Supplementary Table S6. Mean CD4^+^:CD8^+^ ratio at day 14 of production.

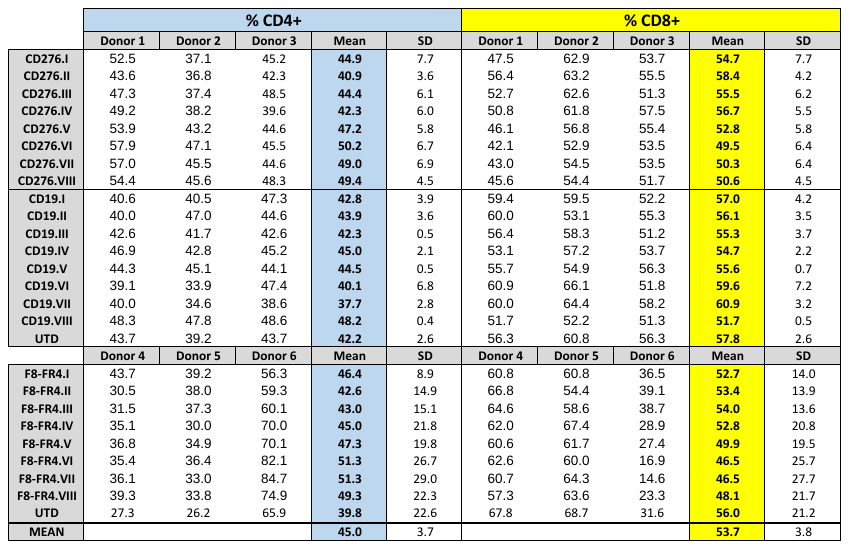

### Supplementary Table S7. Mean CAR expression assessed at d14 by GFP and MycTag expression.

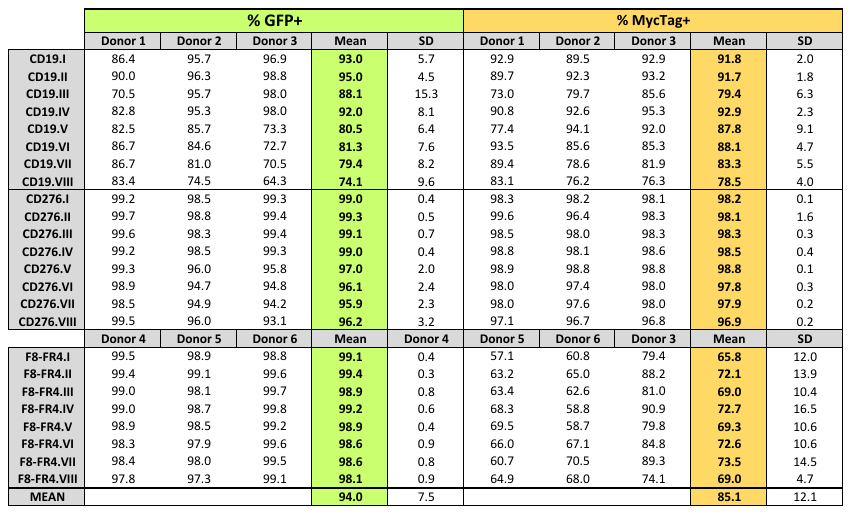

### Supplementary Table S8. GFP and MycTag expression in one representative donor

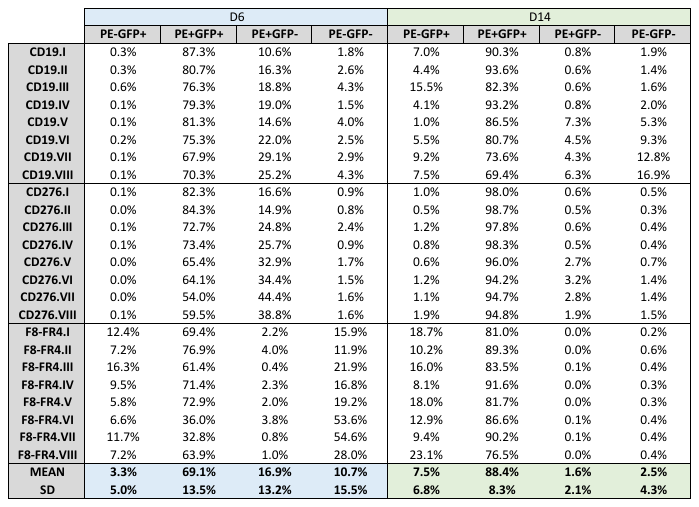

### Supplementary Table S9. Killing capacity significance compared to UTD T cells.

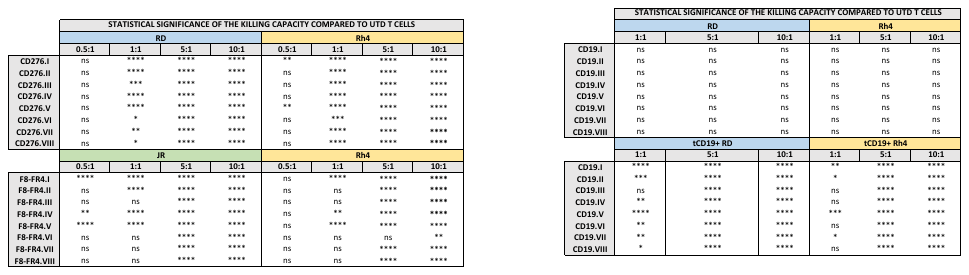

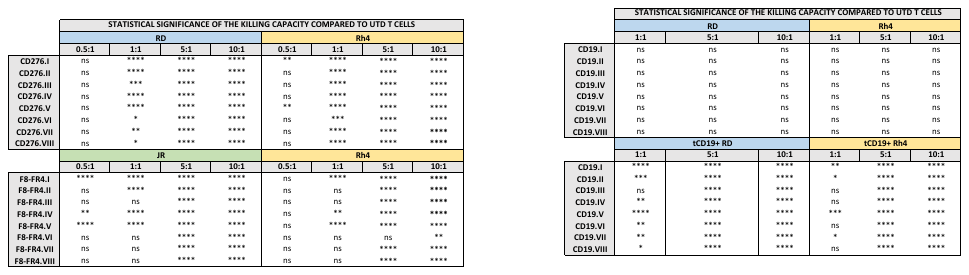

The statistical significance of the differences between experimental and control groups was assessed using Dunnett’s multiple comparison test following a two-way ANOVA (p>0.05 (ns), p ≤ 0.05 (*), p ≤ 0.01 (**), p ≤ 0.001 (***), p ≤ 0.0001 (****)).

### Supplementary Table S10. Summary of memory phenotype measurements for three donors.

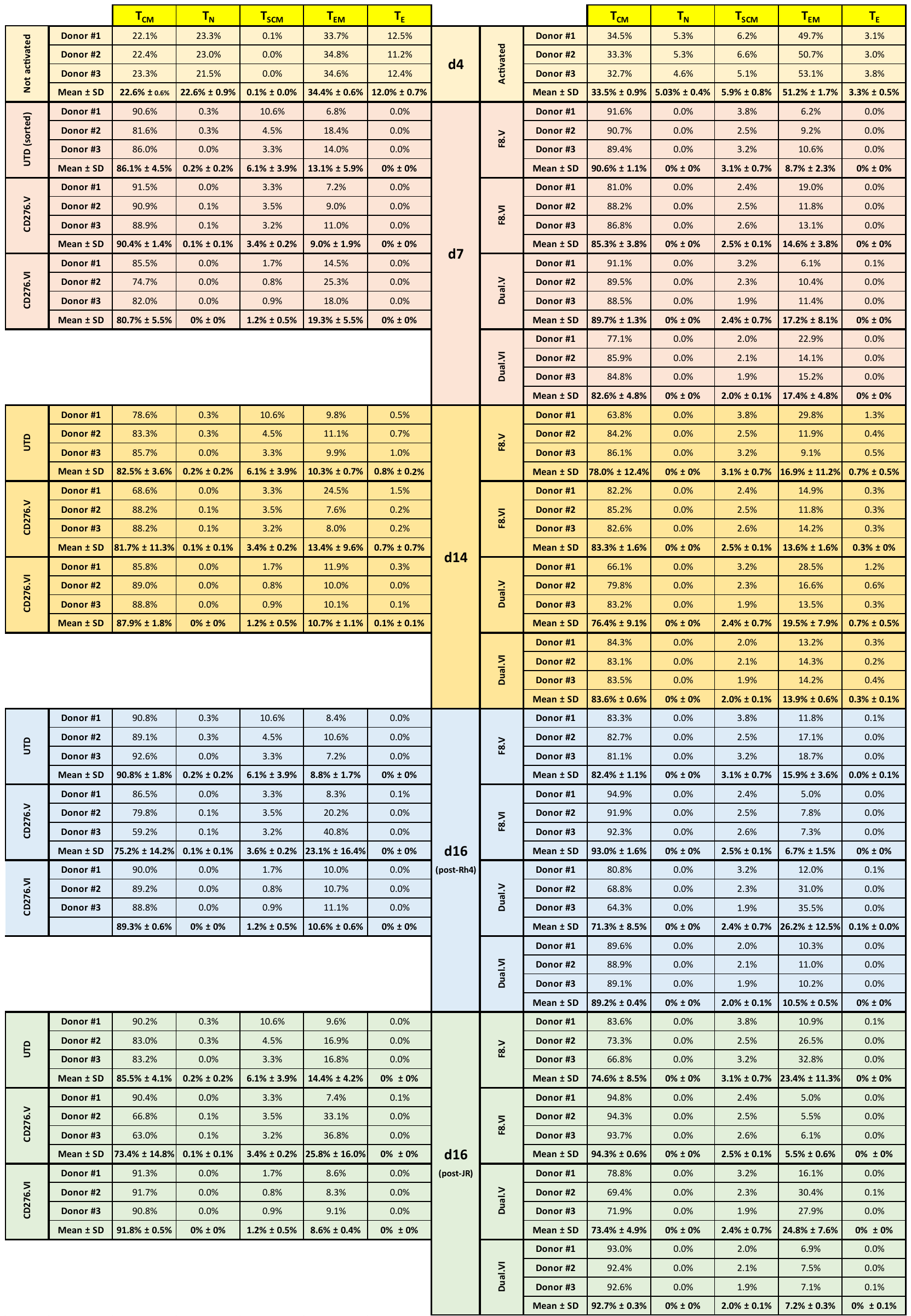

### Supplementary Table S11. Summary of exhaustion phenotype measurements for three donors.

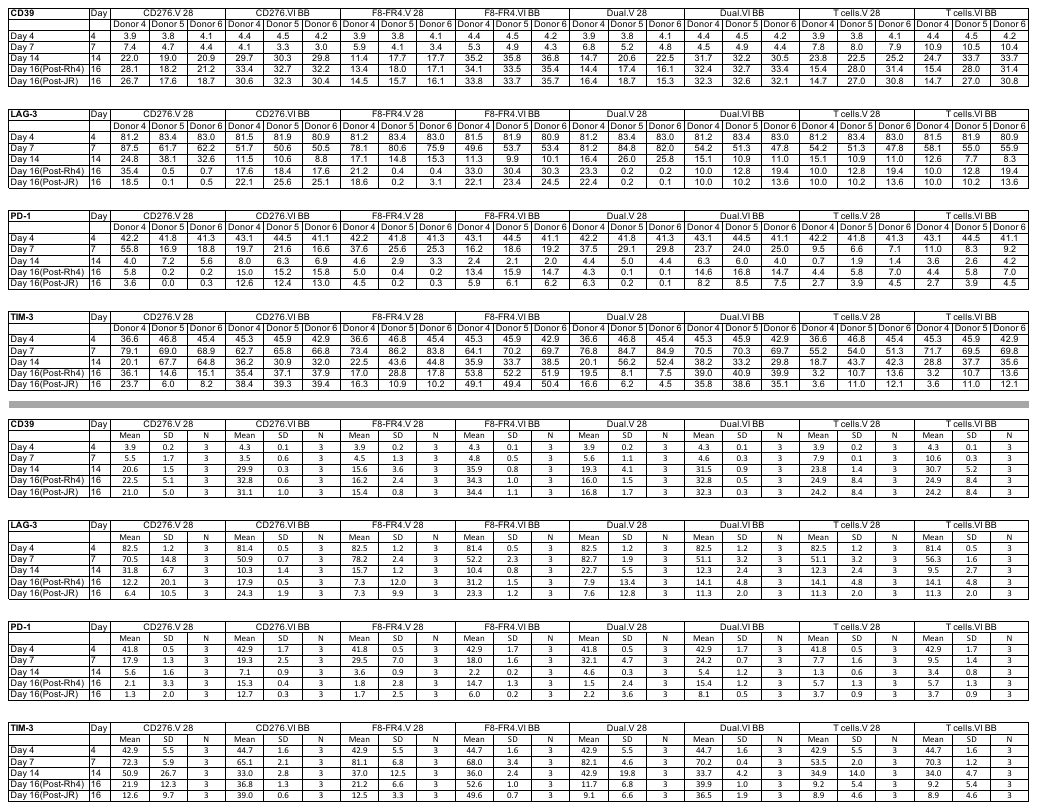

**Supplementary Figures:**

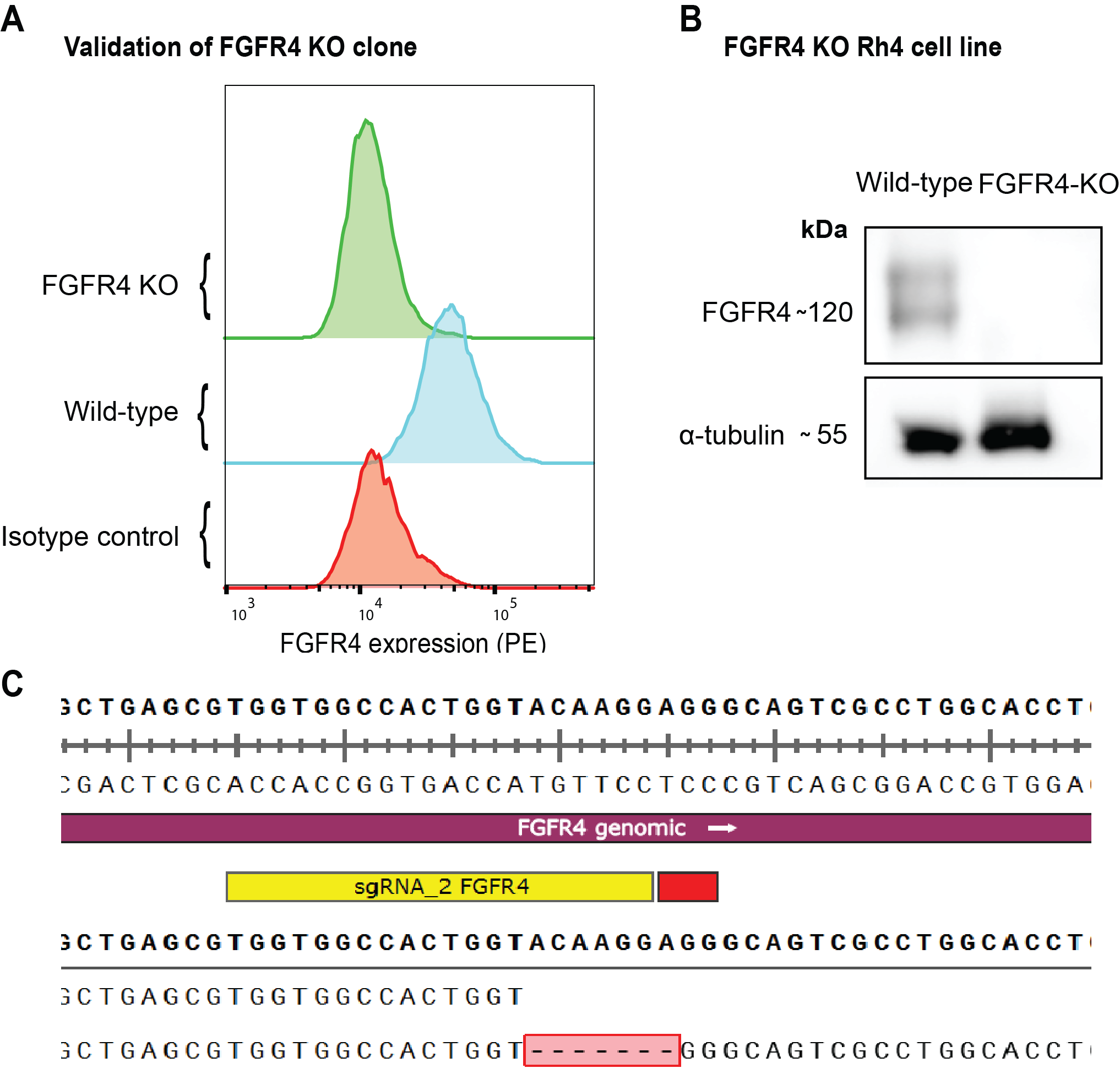

### Supplementary Fig. S1. Validation of FGFR4 KO Rh4 cell line.

(**A**) Flow Cytometry detection of FGFR4 in FGFR4 KO clone (green), compared to Rh4 wild-type (cyan) and isotype control (red). (**B**) FGFR4 expression levels were evaluated by WB. No detection of FGFR4 was visible in the second lane, corresponding to the FGFR4 KO clone. (**C**) Sequencing of PCR amplified fragments cloned by TOPO TA cloning confirmed the introduction of a 7 bp-gap with consequent frameshift mutation. This change in the sequence in exon 6 results in no expression of FGFR4.

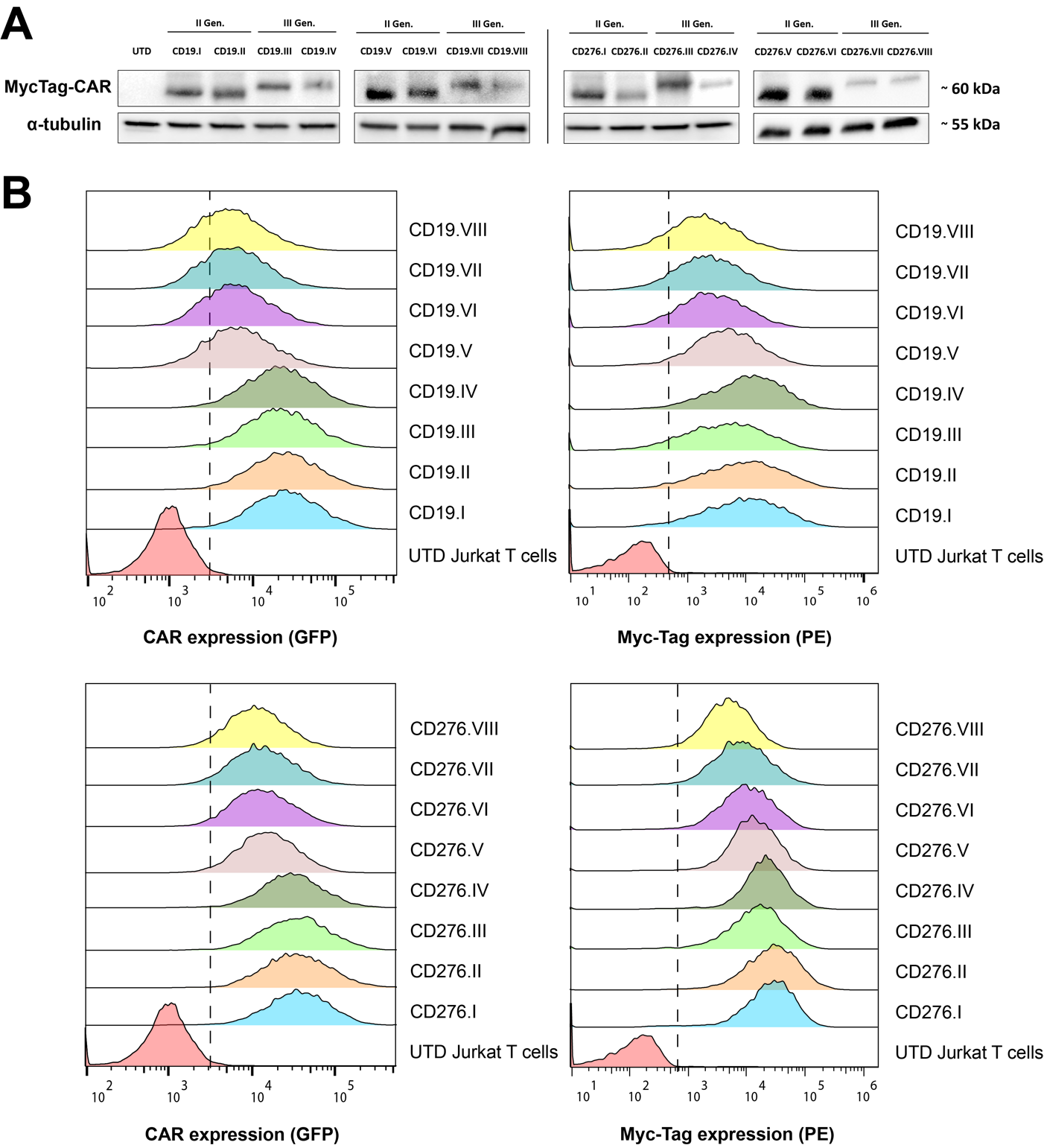

### Supplementary Fig. S2. Expression levels of the different CARs on Jurkat T cells.

(**A)** Western blotting analysis showed high CAR expression levels in all the constructs of interest by MycTag detection. The respective bands confirmed a predicted CAR size of ca. 60 kDa with a visible shift between the second- and third-generation CARs. (**B)** FACS analysis on living GFP-expressing CAR Jurkat T cells performed with an anti-MycTag PE-conjugated antibody confirmed high and homogenous surface expression of the CARs.

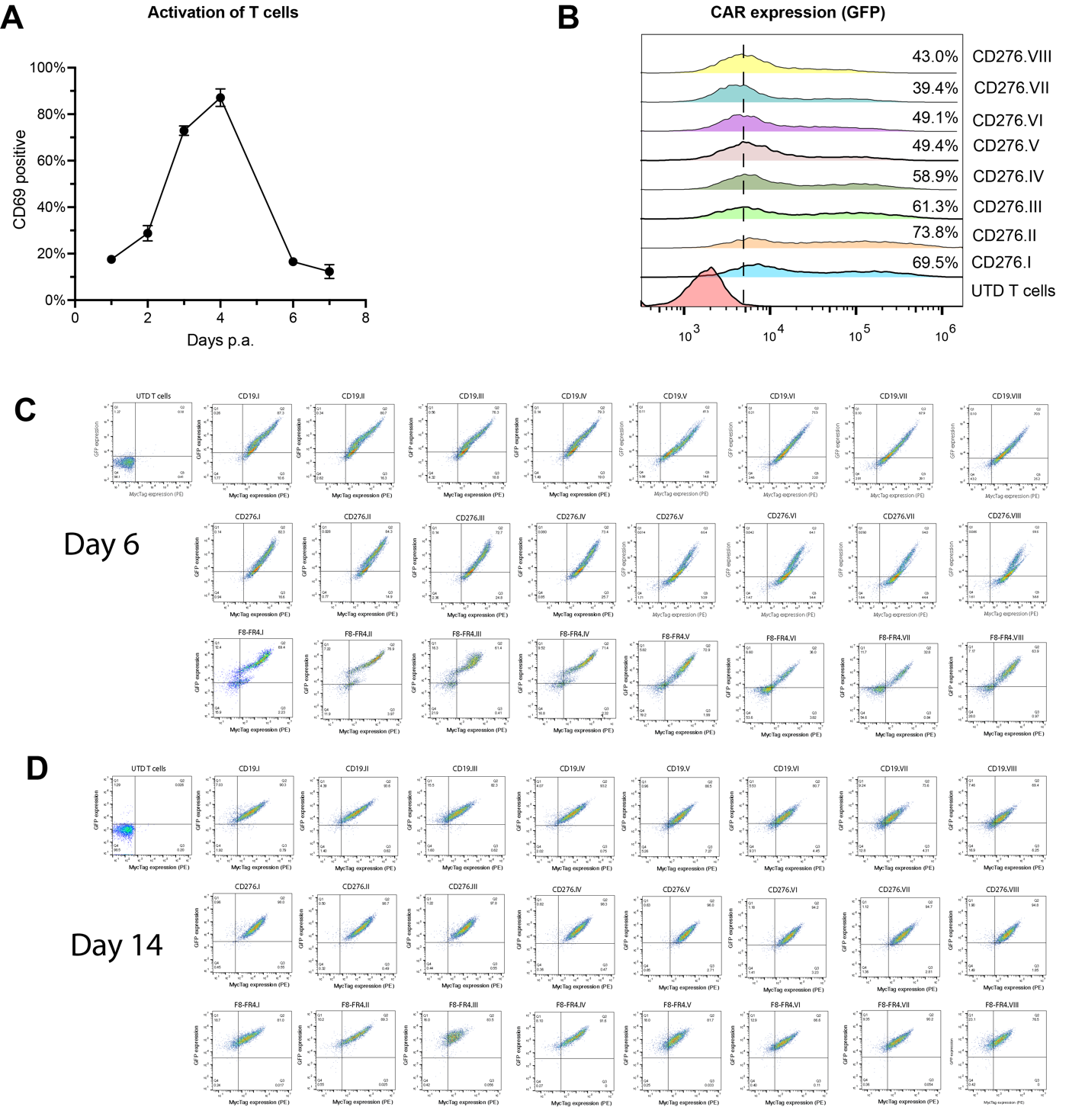

### Supplementary Fig. S3. Activation profile of T cells and lentiviral infection efficiency.

(**A**) T cells were purified from PBMCs and incubated on day 1 with anti-CD3/CD28 activators. During the initial T cell expansion, CD69 marker expression was monitored and resulted at the highest levels on day 4. (**B**) T cells were infected with CAR-expressing lentiviruses on day 4 and transduction efficiency was >40% in most of the tested CAR T cells. Infection efficiency and CAR expression efficiency monitored by GFP expression and MycTag expression, shown for one representative donor at Day 6 (**C**), and at day 14 (**D**).

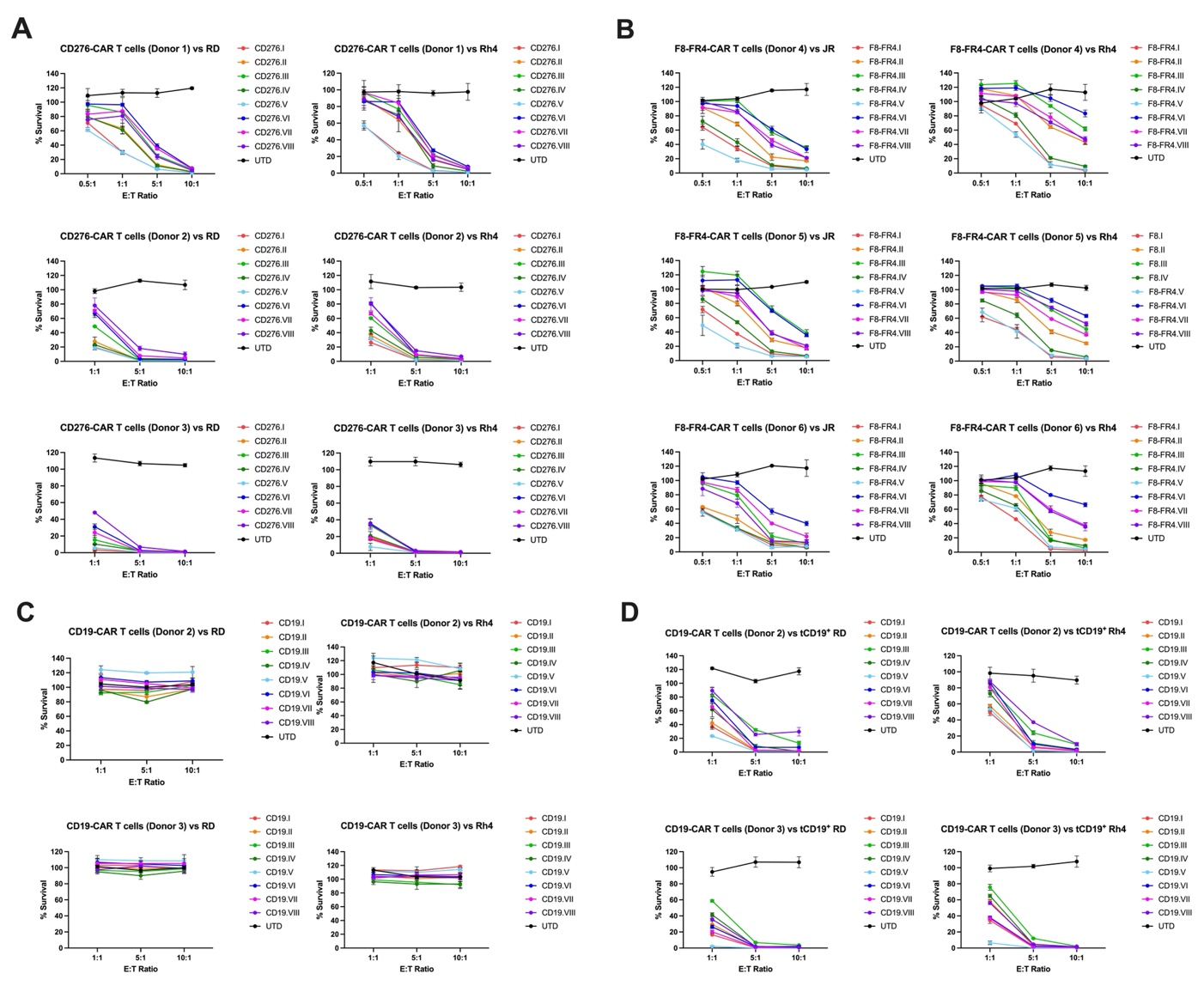

### Supplementary Fig. S4. Evaluation of killing capacity by CD276-, F8-FR4-, and CD19-directed CAR T cells after co-incubation with RD, Rh4 and JR cell lines.

(**A**) CD276-CAR T cells were co-incubated for 48h with fLuc^+^ RD and Rh4 at different E:T ratios showed potent killing of RMS cells in the three donors. CD276.V (cyan) was selected as the most potent CAR construct. (**B**) F8-FR4-CAR T cells from three different donors were co-incubated with fLuc^+^ JR and Rh4 showed high killing capacity at E:T ratios of 5:10 and 10:1. Similarly to the results observed with CD276 CAR T cells, F8-FR4.V CAR T cells outperformed the other experimental groups. (**C**) CD19-CAR T cells, used here as negative control, were co-incubated with fLuc^+^ RD and Rh4 and confirmed no unspecific killing of RMS cells. (**D**) CD19-CAR T cells, used here as positive control, were co-incubated with tCD19^+^ fLuc^+^ RD and Rh4. High killing efficacy was confirmed for all CD19-CAR constructs.

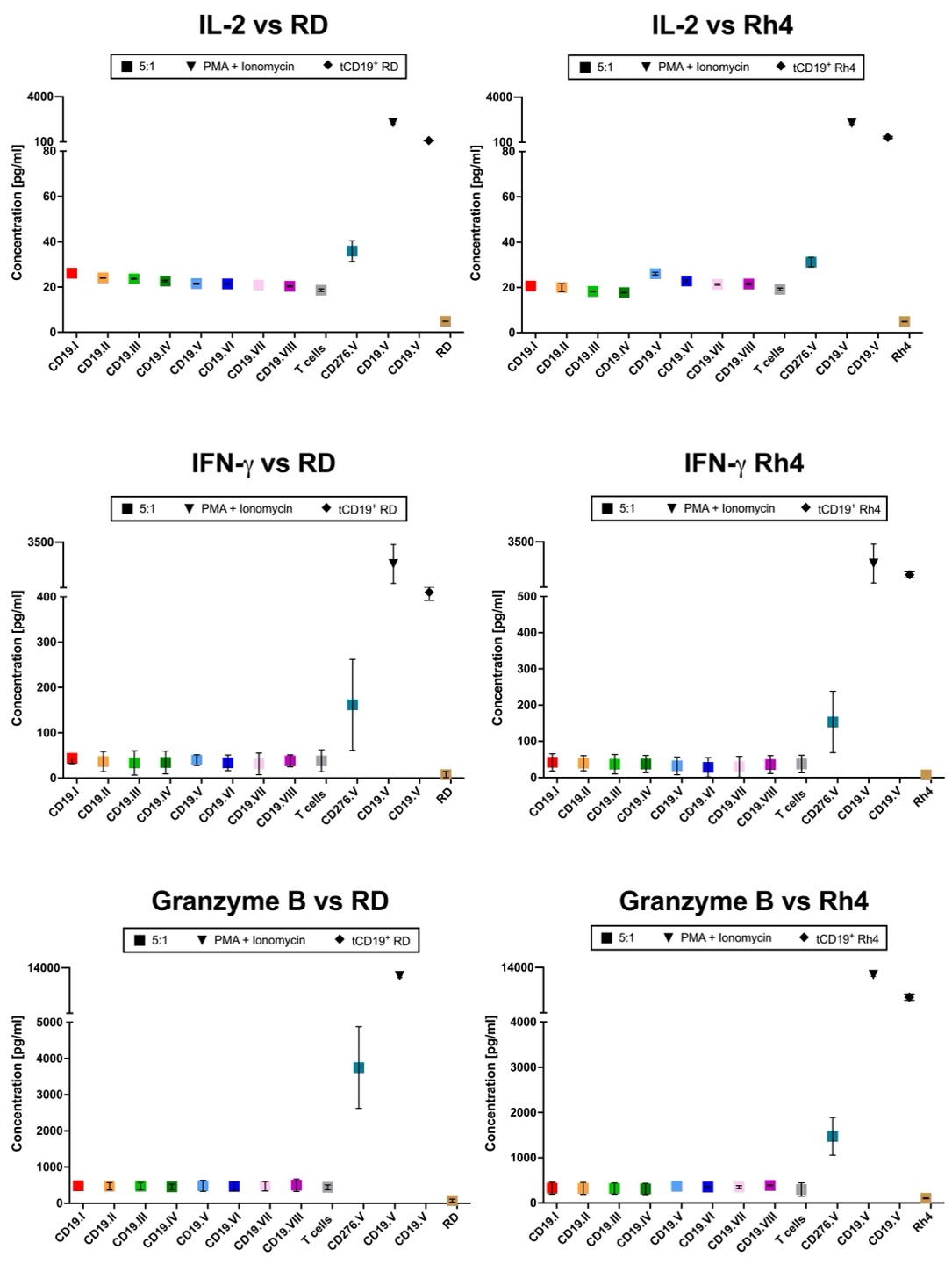

### Supplementary Fig. S5. Cytokine release by CD19-CAR T cells after 24h co-incubation with RD and Rh4 cells.

Concentrations of IL-2, IFN-γ, and Granzyme B in the supernatant released during co-incubation of CD19-CAR T cells with RD (left panels) and Rh4 cells (right panels) at the E:T ratio of 5:1 were measured by ELISA after 24h. For CD19-CAR constructs background levels of cytokines were detected. As positive control CD276.V-CAR T cells were used, as well as CD19.V-CAR T cells were also incubated with RD and Rh4 cells overexpressing a truncated version of CD19 (tCD19).

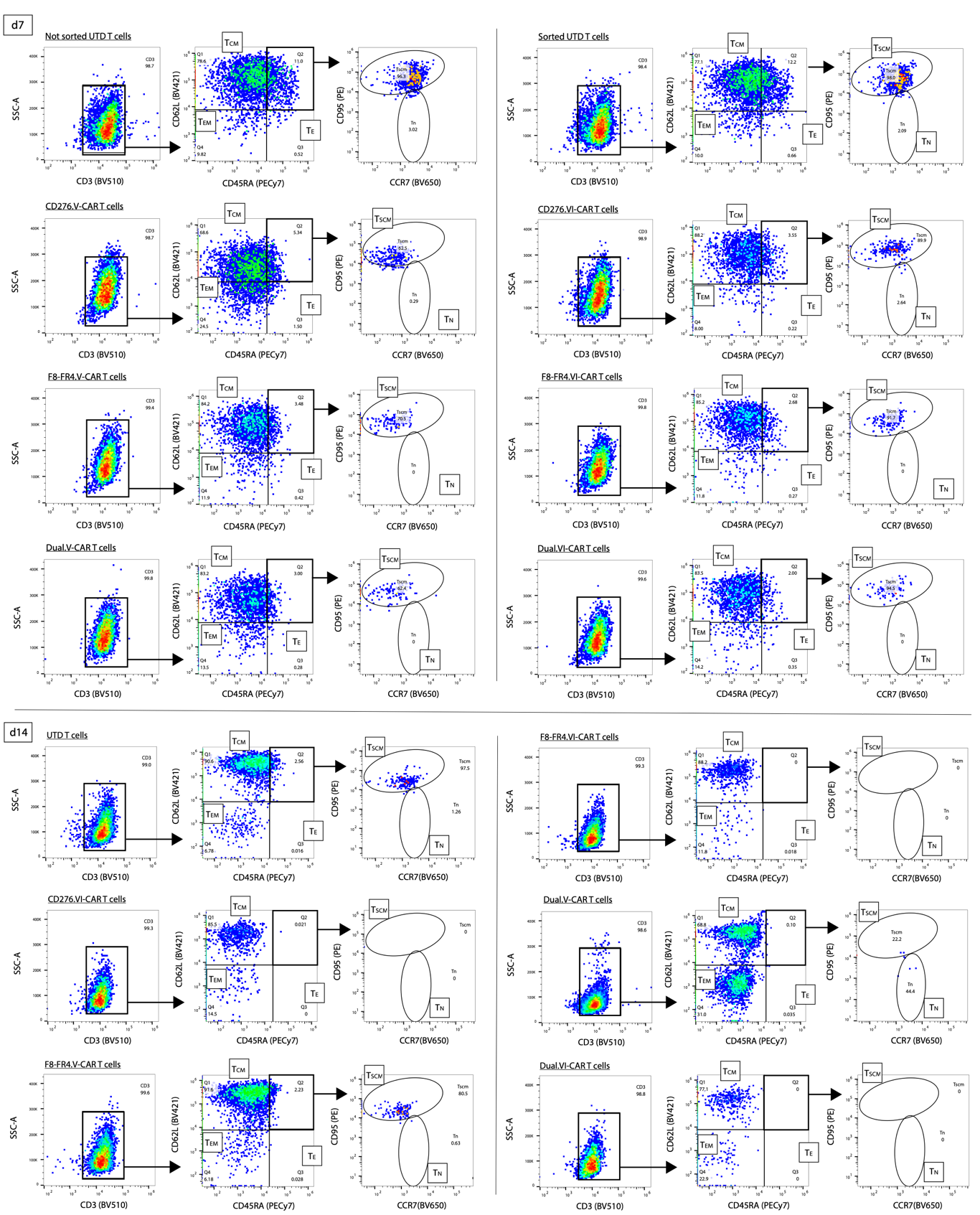

### Supplementary Fig. S6. Phenotypic characterization of CAR T cells on day 7 and day 14 before co-incubation.

Flow Cytometry during CAR T cells manufacturing was used to quantify percentages of cell memory (TCM), effector memory (TEM) and effector (TE) T cells on CD3 positive cells by CD45RA and CD62L staining. Naïve (TN) and stem cell-like (TSCM) T cells were quantified on CD3^+^/CD45RA^+^/CD62L^+^ based on CD95 and CCR7 staining. On day 14, before co-incubation assays activated CAR T cells showed very high percentage of TCM cells and lower percentages of the other cell populations.

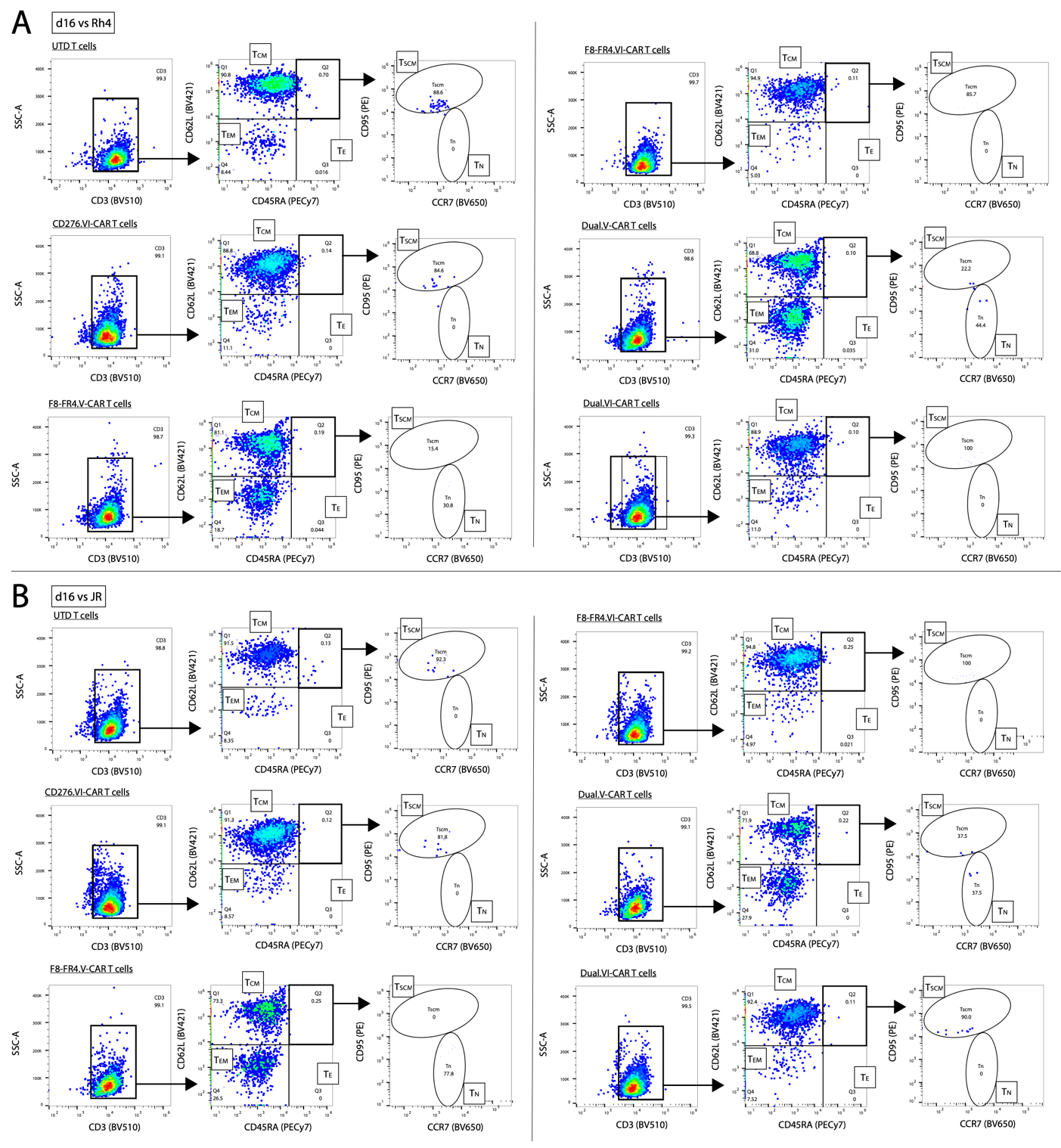

### Supplementary Fig. S7. Phenotypic characterization of CAR T cells on day 16 after co-incubation with Rh4 and JR cells.

Flow Cytometry during CAR T cells manufacturing was used to quantify percentages of cell memory (TCM), effector memory (TEM) and effector (TE) T cells on CD3 positive cells by CD45RA and CD62L staining. Naïve (TN) and stem cell-like (TSCM) T cells were quantified on CD3^+^/CD45RA^+^/CD62L^+^ based on CD95 and CCR7 staining. (**A**) On day 16, after co-incubation assays with Rh4 cells, almost 60% of activated CAR T cells had a TCM phenotype, and ca. 40% a TEM phenotype. (**B**) On day 16, after cytotoxicity experiments with JR, almost 70% were TCM cells, whereas ca. 30% were TEM cells.

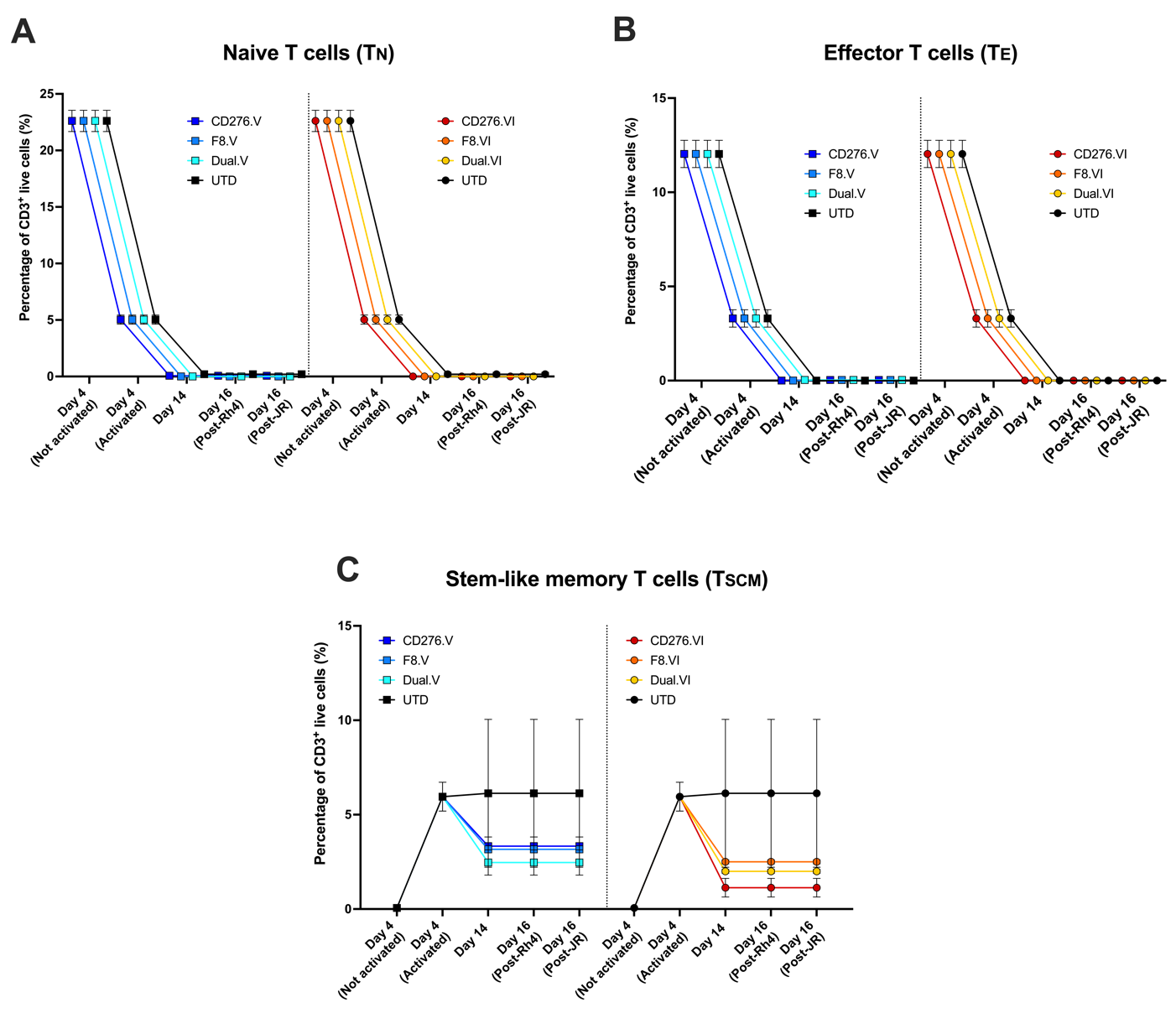

### Supplementary Fig. S8. Phenotypic characterization of CAR T cells during expansion and co-incubation experiments.

Flow Cytometry during CAR T cells manufacturing was used to quantify percentages of cell memory (TCM), effector memory (TEM) and effector (TE) T cells on CD3 positive cells by CD45RA and CD62L staining. Naïve (TN) and stem cell-like (TSCM) T cells were quantified on CD3^+^/CD45RA^+^/CD62L^+^ based on CD95 and CCR7 staining. (**A**) On day 4 before infection, almost 25% of not-activated T cells were TN. As expected, T cells activated for 4 days showed a lower TN population, and there were no naïve CAR T cells detected from day 14. (**B**) On day 4 before infection, less than 15% of not-activated T cells were TE. T cells activated for 4 days showed a lower TE population, and there were no effector CAR T cells detected from day 14. (**C**) ~5% of activated T cells were TSCM on day 4. Population decreased by ca. 2-fold during expansion and co-incubation experiments.

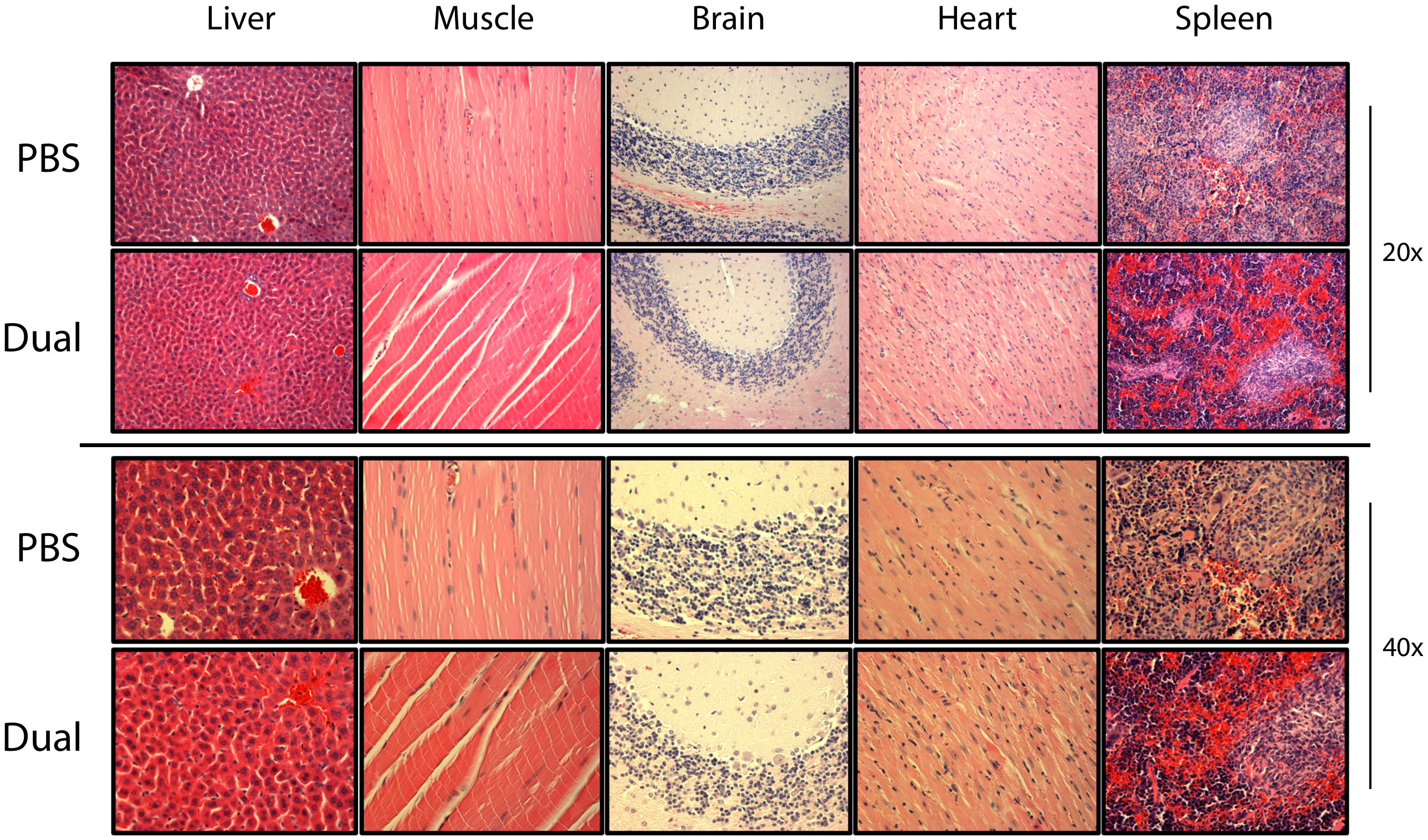

### Supplementary Fig. S9. No visible toxicity detected in normal mouse tissues by IHC.

IHC analyses performed on normal tissues show no apparent side effects due to the CAR T cell treatment. Hematoxylin and Eosin staining performed in untreated and CAR T cell-treated mice exhibited no evident side effects in normal tissues, such as liver, muscle, brain, heart, and spleen.
